## supporting_material.pdf for "Sequential water and headgroup merger: Membrane poration paths and energetics from MD simulations"

(Dated: June 13, 2020)

**CONTENTS**

PMF Recovery 2

Implementation Details 6

Gizmo Parameterization 7

Gizmo Simulations: PMF ensembles and committor seed ensembles 10

Committor Dataset 14

Committor Regression Model Scores 16

References 16

### PMF RECOVERY

In this section, we describe the process to recover unbiased PMFs for the membrane (and solvent) given an ensemble of gizmo biased simulations. Our scheme describes the membrane and gizmo as two coupled systems, recovers their joint probability distribution via the weighted histogram analysis method (WHAM)[1], marginalizes over coupling terms, and then uses maximum likelihood to recover the independent, unbiased distribution for the poration CVs of interest.

Consider a system of particles, such as a solvated lipid membrane, with coordinates $\mathbf{x}$ and potential energy $V_x(\mathbf{x})$ in the canonical ensemble at inverse temperature $\beta = 1/(k_B T)$. For any observable $a(\mathbf{x})$, for example a reaction coordinate for pore opening, the free energy $F(a) = -k_B T \ln p(a)$ is defined via the probability density

$$p(a) = \int_X e^{-\beta V_x(\mathbf{x})} \delta(a - a(\mathbf{x})) d\mathbf{x} / Z_x, \tag{S.1}$$

where $Z_x = \int_X e^{-\beta V_x(\mathbf{x})} d\mathbf{x}$ is the partition function.

If a second set of particles is added, e.g. of a pore gizmo, with coordinates $\mathbf{y}$ and potential energy $V_y(\mathbf{y})$, the combined system comprises two subsystems 'x' and 'y'. For the case of an additive total potential energy of the combined system,

$$V(\mathbf{x}, \mathbf{y}) = V_x(\mathbf{x}) + V_y(\mathbf{y}), \tag{S.2}$$

the subsystems are uncoupled, non-interacting, and have independent statistics. We refer to this case as the reference system, $V(\mathbf{x}, \mathbf{y})$ as the reference potential, and $Z = \int_X \int_Y e^{-\beta V(\mathbf{x},\mathbf{y})} d\mathbf{x} d\mathbf{y}$ as the reference partition function.

Next, consider the case at hand, with three observables, $a(\mathbf{x})$, $b(\mathbf{x}, \mathbf{y})$, and $c(\mathbf{y})$, that are used to define an extended ('biased') potential energy function, which contains coupling terms between the two subsystems x and y:

$$\widetilde{V}^{(l)}(\mathbf{x}, \mathbf{y}) = V_x(\mathbf{x}) + V_y(\mathbf{y}) + V_a^{(l)}(a(\mathbf{x})) + V_b^{(l)}(b(\mathbf{x}, \mathbf{y})) + V_c^{(l)}(c(\mathbf{y})). \tag{S.3}$$

Here, the tilde indicates biasing and $(l)$ indicates the simulation window for cases where multiple $\widetilde{V}^{(l)}$ are simulated. Often, and also in the main text, if $a$ is a coordinate of interest that is not energetically biased, then $V_a^{(l)} = 0$. Here, the $V_b^{(l)}$ term energetically couples the two subsystems. In our membrane-gizmo system, $b(\mathbf{x}, \mathbf{y})$ is the Lennard-Jones potential between the membrane and the gizmo, and $V_b^{(l)}$ is a linear scaling function to apply different coupling strengths to different simulations. As a result, the statistics of systems x and y are no longer independent. Like $a(\mathbf{x})$, $c(\mathbf{y})$ can also be used to energetically bias the system (e.g. the harmonic biasing restraint of the gizmo belt). In a more general application, additional coordinates beyond $a, b, c$ can also be included in this formalism, either to impose biasing or as collective variables of mechanistic interest. With this potential $\widetilde{V}^{(l)}$ the biased joint probability density of the observables is given by

$$\widetilde{p}^{(l)}(a, b, c) = \int_X \int_Y e^{-\beta \widetilde{V}^{(l)}(\mathbf{x},\mathbf{y})} \delta_a \delta_b \delta_c d\mathbf{x} d\mathbf{y} / \widetilde{Z}^{(l)}, \tag{S.4}$$

where $\widetilde{Z}^{(l)} = \int_X \int_Y e^{-\beta \widetilde{V}^{(l)}(\mathbf{x},\mathbf{y})} d\mathbf{x} d\mathbf{y}$ and where $[\delta_a, \delta_b, \delta_c] = [\delta(a - a(\mathbf{x})), \delta(b - b(\mathbf{x}, \mathbf{y})), \delta(c - c(\mathbf{y}))]$. The integrand of (S.4) is nonzero only when $\delta_a \delta_b \delta_c \neq 0$, implying that the corresponding exponentiated energy terms are constants that can come outside the integrand, giving

$$\widetilde{p}^{(l)}(a, b, c) = w^{(l)}(a, b, c) \int_X \int_Y e^{-\beta V^{(l)}(\mathbf{x},\mathbf{y})} \delta_a \delta_b \delta_c d\mathbf{x} d\mathbf{y} / \widetilde{Z}^{(l)}, \tag{S.5}$$

where we defined a biasing weight $w^{(l)}(a, b, c) = e^{-\beta(V_a^{(l)}(a)+V_b^{(l)}(b)+V_c^{(l)}(c))}$. The analogous probability density in the uncoupled reference system (for which $w(a, b, c) = 1$) is

$$p(a, b, c) = \int_X \int_Y e^{-\beta V(\mathbf{x},\mathbf{y})} \delta_a \delta_b \delta_c d\mathbf{x} d\mathbf{y} / Z. \tag{S.6}$$

Combining equations (S.5) and (S.6) yields

$$p(a, b, c) = \frac{\widetilde{p}^{(l)}(a, b, c) \widetilde{Z}^{(l)}}{w^{(l)}(a, b, c) Z}. \tag{S.7}$$

Eq (S.7) permits sampling a biased distribution $\widetilde{p}^{(l)}$, with coupled subsystems x and y, and then recovering the unbiased joint distribution $p(a, b, c)$ from which $p(a)$ can be recovered. Equation (S.7) is also the basis for multi-state reweighting methods such as WHAM and MBAR [1, 2] that are used to recover an unbiased distribution from an ensemble of simulations, each using a different biasing potential (such as $\widetilde{V}^{(l)}$) or temperature and each, typically, tuned to sample a small region of configuration space.

Here, we numerically estimate $p(a, b, c)$ by applying WHAM to the ensemble of gizmo biased simulations. For this, we discretize coordinates $a, b, c$ into bins with indices $i, j, k$, respectively. In this notation, $\widetilde{p}_{ijk}^{(l)}$ is the integral of $\widetilde{p}^{(l)}(a, b, c)$ over the voxel indexed by $i, j, k$, and is approximated here by the sample fraction $\widetilde{N}_{ijk}^{(l)} / \widetilde{N}^{(l)}$ within this voxel.

For ideal sampling, $p_i$ is obtained by marginalizing $p_{ijk}$ (the distribution recovered by WHAM). Here, although the sampling within individual windows is well converged with respect to unrestrained degrees of freedom (see the autocorrelation analysis in the Gizmo Simulations section), we know that the ensemble does not cover the full $i, j, k$ space. By construction, the gizmo state is coupled to the membrane state, which is precisely how it controls pore formation.

To proceed, we marginalize only over the coupling coordinate $b$ ($j$ indices) to obtain $p_{ik} = \sum_j p_{ijk}$. This requires our simulation ensemble to only have sufficient sampling of the conditional distribution $p_{j|ik}$, which is a lesser requirement than sampling the full joint distribution. Because $p_{ik}$ is not fully sampled, marginalization yields a "masked" joint distribution $\hat{p}_{ik} \propto p_{ik} m_{ik}$, where $m_{ik} \in \{0, 1\}$ is a binary mask that denotes whether bin $ik$ was sampled by the simulated ensemble. This mask also breaks normalization. Recall, from the continuous case above, that

$$\begin{aligned} p(a,c) &= \int_X \int_Y e^{-\beta V(\mathbf{x},\mathbf{y})} \delta_a \delta_c d\mathbf{x} d\mathbf{y} / Z \\ &= \left( \int_X e^{-\beta V_x(\mathbf{x})} \delta_a d\mathbf{x} / Z_x \right) \left( \int_Y e^{-\beta V_y(\mathbf{y})} \delta_c d\mathbf{y} / Z_y \right) \\ &= p(a)p(c), \end{aligned} \tag{S.8}$$

implying that $p_{ik} = p_i p_k$ here in the discrete case, such that

$$\hat{p}_{ik} \propto p_i p_k m_{ik}. \tag{S.9}$$

For a fixed index $k$, $p_k$ is constant and can be omitted. Using Eq (S.9) and $\gamma_{ik} = m_{ik} N_{ik} / \hat{p}_{ik}$ yields, after proper normalization,

$$\frac{N_{ik}}{N_k} = \frac{p_i \gamma_{ik}}{Z_k}, \tag{S.10}$$

where $N_k = \sum_i N_{ik}$, $Z_k = \sum_i p_i \gamma_{ik}$, and $N_k$ and $N_{ik}$ are totals over all windows.

Equation (S.10) fits the previously established scenario for biased sampling (Eq (S.4)). For "window" $k$, the unknown distribution $p_i$ was sampled with the multiplicative bias $\gamma_{ik}$, yielding the bin counts $N_{ik}$. Because each $k$ window only samples a segment of the bins in $i$ (similar to umbrella sampling), we derive a maximum likelihood estimate for $p_i$ (and $Z_k$), which, as it turns out, yields the WHAM equations [1].

To this aim, we express the probability of all observed bin counts $\{N_{ik}\}$ conditioned on unknowns $\{p_i\}$ and $\{Z_k\}$, and assuming uncorrelated observations, as

$$P(\{N_{ik}\} | \{p_i\}, \{Z_k\}) = \prod_k \prod_i (p_i \gamma_{ik} / Z_k)^{N_{ik}}. \tag{S.11}$$

The corresponding log likelihood is

$$L = \sum_k \sum_i N_{ik} (\ln p_i + \ln \gamma_{ik} - \ln Z_k). \tag{S.12}$$

Setting $\frac{\partial L}{\partial p_i} = 0$ we obtain

$$\frac{\partial L}{\partial p_i} = 0 = \sum_k N_{ik} (1/p_i) - \sum_k N_k \gamma_{ik} / Z_k, \tag{S.13}$$

and solving for $p_i$ gives

$$p_i = \frac{\sum_k N_{ik}}{\sum_k N_k \gamma_{ik}/Z_k}, \tag{S.14}$$

where, (as defined earlier)

$$Z_k = \sum_i p_i \gamma_{ik}. \tag{S.15}$$

Together, (S.14) and (S.15) can be solved iteratively, and are equivalent to the WHAM equations [1]. Finally, from $p_i$ we obtain the discretized free energy estimate $F_i = -k_B T \ln p_i$, i.e. the PMF.

#### Implementation Details

The correspondences between the observables $a$, $b$, and $c$, in Eq. (S.3) and the terms of the biased potential energy from the main text are listed in Table S1. Briefly, $a_1$ and $a_2$ are CVs for the lipid and solvent, and do not contribute to the potential energy. For computing 1D PMFs, $a_2$ and $\phi_2$ are omitted from the analysis. $b_1$ and $b_2$ comprise the coupling between the gizmo and the lipid-solvent system. $c$ is the gizmo belt's collective radial coordinate $\xi$.

In all simulations, we used $\lambda^{(l)}_{\text{WX}}$=1, meaning the water-repulsive belt atoms were not switched off (instead, the belt radius was varied), however the energy $V^{\text{WX}}_{\text{LJ}}$ was tracked to allow proper marginalization over it. The chain-tail repulsion scaling factor $\lambda^{(l)}_{\text{CX}}$ was varied between 0 and 1 to drive pore opening. As suggested by Tan et al [3], and because $V^{\text{CX}}_{\text{LJ}}$ varies by many orders of magnitude due to the hard core Lennard-Jones repulsion, we used a geometric rather than constant bin spacing for $b_1$.

TABLE S1: Observables used to compute PMFs from gizmo biased simulations

| observable type | observable | biasing energy term |
|---|---|---|
| lipid-solvent CV | $a_1(\mathbf{x}) = \phi_1(\mathbf{x})$ | $0 * \phi_1(\mathbf{x})$ |
| lipid-solvent CV | $a_2(\mathbf{x}) = \phi_2(\mathbf{x})$ | $0 * \phi_2(\mathbf{x})$ |
| gizmo-system coupling | $b_1(\mathbf{x},\mathbf{y}) = V^{\text{CX}}_{\text{LJ}}(\mathbf{x},\mathbf{y})$ | $\lambda^{(l)}_{\text{CX}} V^{\text{CX}}_{\text{LJ}}(\mathbf{x},\mathbf{y})$ |
| gizmo-system coupling | $b_2(\mathbf{x},\mathbf{y}) = V^{\text{WX}}_{\text{LJ}}(\mathbf{x},\mathbf{y})$ | $\lambda^{(l)}_{\text{WX}} V^{\text{WX}}_{\text{LJ}}(\mathbf{x},\mathbf{y})$ |
| gizmo belt | $c(\mathbf{y}) = \xi(\mathbf{y})$ | $k/2(\xi(\mathbf{y}) - \xi^{(l)}_0)^2$ |

### GIZMO PARAMETERIZATION

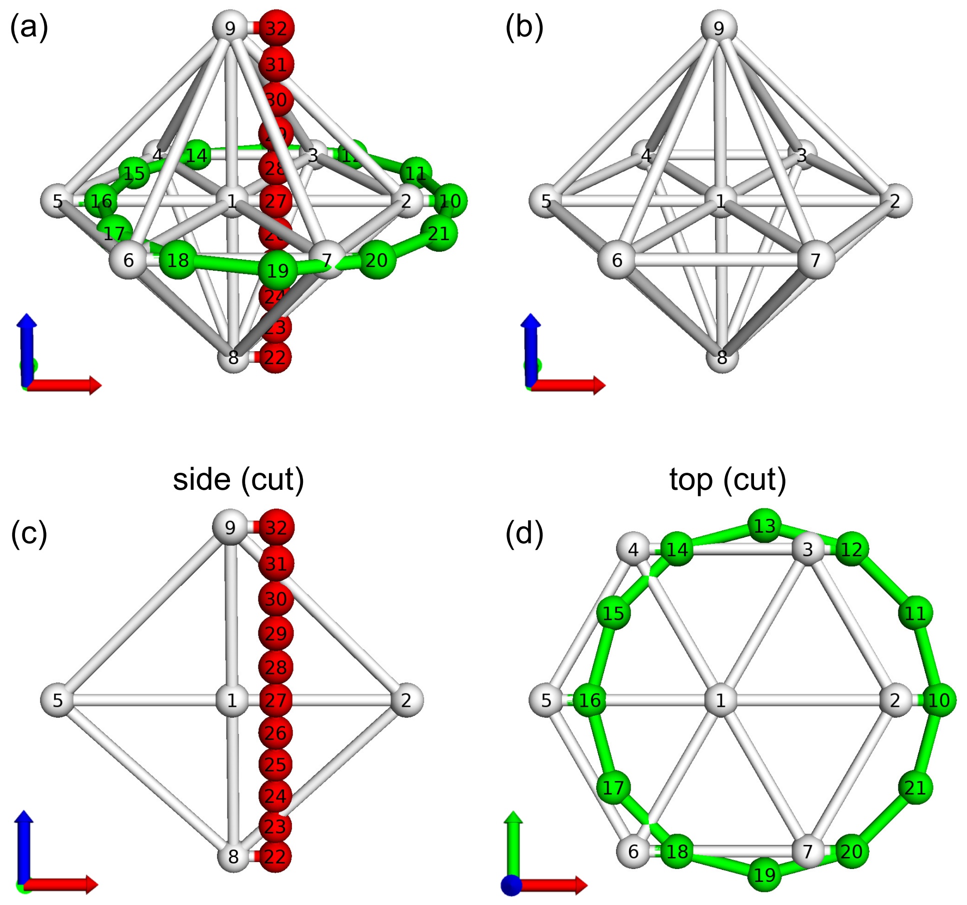


FIG. S1: Four views of the H-gizmo. (a) All atoms. (b) The GG skeleton. (c) Side cutaway view with the belt hidden. (d) Top cutaway view with the chain hidden. In all panels, GG/CX/WX atoms are colored white/red/green. For illustration purposes, the atoms are not drawn to scale. Also for illustration purposes the chain and belt are offset horizontally to show the bonds that attach them to the GG skeleton (i.e. atom pairs (9, 32), (4, 14)). Aside from these bonds, the gizmo is shown in its energetic ground state (all bonds at zero energy)

The gizmo (Fig. S1 and Tab. S2) is built from three non-standard atom types — GG, CX, and WX — as described in the main text. The H-gizmo structure consists of a skeleton of 9 GG atoms arranged in a hexagonal bypyramid (Fig. S1(b)), an 11 atom CX chain along its six-fold rotation axis (Fig. S1(c)), and a 12 atom WX belt that circumscribes its hexagonal perimeter (Fig. S1(d)). The I-gizmo only differs by having the three centermost CX chain beads replaced with GG beads (atoms 26, 27, 28). The terminal beads of the CX chain are bound to the skeleton's axial vertices (atoms 8 and 9). The WX belt is bound, by alternating atoms, to the six equatorial GG atoms of the skeleton. All bonds, shown in Fig. S1, are implemented as harmonic springs. The chain uses angle bending potentials to balance its flexibility. The belt has six WX atoms (even indices) that are bound to the skeleton and hence have a restricted range of motion compared to the other six WX atoms (odd indices). Additional angle restraints were used to keep these odd indexed WX atoms arranged in a hexagonal shape and to prevent large displacements out of the horizontal plane.

The parameters for bond stretching and angle bending potentials are given in Tab. S2. Because the gizmo has six-fold rotation symmetry about its central axis, the table only lists the unique potential parameters and not the symmetric copies. For example, the bond potential between atoms 1 and 3 (omitted) has the same parameters as that for atoms 1 and 2 (included).

TABLE S2: Parameters for gizmo bond stretching and bond angle bending, both of which use harmonic potentials. All units are those used in the Gromacs forcefield specification (this includes the use of both degrees and radians for angle bending terms).

| gizmo bond stretching potentials | | | |
|---|---|---|---|
| atoms | $r_0$ [nm] | $k$ [kJ/mol/nm$^2$] | description |
| 1,2 | 2 | 1250 | skeleton center to perimeter |
| 2,3 | 2 | 1250 | skeleton perimeter |
| 1,9 | 2 | 1250 | skeleton center to apex |
| 2,9 | 2.83 | 1250 | skeleton perimeter to apex |
| 10,11 | 1.04 | 100 | belt to belt |
| 2,10 | 0 | 250 | belt to skeleton attachment |
| 22,23 | 0.4 | 1250 | chain to chain |
| 22,8 | 0 | 1250 | chain to skeleton attachment |

| gizmo bond angle bending potentials | | | |
|---|---|---|---|
| atoms | $\theta_0$ [deg] | $k$ [kJ/mol/rad$^2$] | description |
| 22,23,24 | 180 | 25 | chain bending |
| 11,13,15 | 120 | 100 | odd belt atoms, hexagonal |
| 11,1,13 | 60 | 200 | odd belt atoms, hexagonal |
| 11,1,17 | 180 | 200 | odd belt atoms, hexagonal |
| 10,1,9 | 90 | 200 | even belt atoms, in-plane |
| 11,1,9 | 90 | 200 | odd belt atoms, in-plane |

This Supporting Information includes a zipped directory of the files used to implement the H- or I-gizmo in a Gromacs simulation (tested with Gromacs versions 5.x). These files include:

1. Gromacs topology and forcefield files (itp and top)
2. An essential dynamics input file (edi) that defines the belt breathing mode
3. Coordinate files (pdb) for the gizmo and the simulated 128 lipid DMPC patch

The included forcefield is a combination of the Berger lipid forcefield [4], downloaded from the Tieleman Lab (`http://wcm.ucalgary.ca/tieleman/downloads`), and the GROMOS 53A6 forcefield [5], combined using the Lemkul Lab's prescription (`http://www.mdtutorials.com/gmx/membrane_protein/index.html`). To this, the gizmo particle definitions (CX, WX, GG) were added. Although no membrane proteins are considered in this study, it would be possible using this combined forcefield.

### GIZMO SIMULATIONS: PMF ENSEMBLES AND COMMITTOR SEED ENSEMBLES

Figures S2 and S3 illustrate the four PMF ensembles used in this study, combining both gizmo types (H- and I-) and both starting pore states (closed and open). The gizmo simulations were set along a path through ($\lambda_{\mathrm{CX}}$, $\xi_0$) control parameter space shown in (a) and (d). This is a similar to schemes using repulsive field potentials, with incrementally adjusted control parameters, to direct similar membrane conformation changes [6]. Here, for the H-Gizmo ensembles, the path from closed (lower left) to open (upper right) in Figure S2 (a) and (d) consists of three segments: 1, chain growth until the TS; 2, belt opening to allow pore expansion; and 3, additional chain growth to push the pore open slightly beyond its metastable radius. For the I-Gizmo ensembles, shown in Figure S3(a) and (d), reaching the TS required higher $\lambda_{\mathrm{CX}}$ values (due the different imposed bias) so only two segments, chain growth and belt opening, were simulated. In all cases, the first 50ns of each window was discarded to equilibration prior to computing statistics (including PMFs).

As the time series raster plots in (b) and (e) show, total simulation length varied between windows. The windows exhibiting slow, bi-modal switching dynamics (indicative of TS crossing) were extended to 2$\mu$s, whereas windows sampling only one side of the TS were run for 200-500ns. The corresponding autocorrelation times, shown in (c) and (f), were computed using the pymbar package [2, 7] (`https://github.com/choderalab/pymbar`), and show that longer simulations were warranted to obtain sufficient statistical sampling ($\approx$10x the autocorrelation time).

Figure S4 illustrates the two committor ensembles used to specifically target the TS region and generate structures for subsequent committor analysis. In both the H-Gizmo (a-c) and I-Gizmo (d-f) cases, the control parameters ($\lambda_{\mathrm{CX}}$, $\xi_0$) were chosen to intersect the paths of the PMF ensembles and to facilitate bi-stable switching (i.e. repeated TS crossing). The H-Gizmo ensemble illustrates that a range of gizmo parameters can facilitate bi-stable switching, whereas the I-Gizmo ensemble was closer to the PMF pathway. Each of these trajectories was strided to 1ns prior to selecting points for committor calculations.

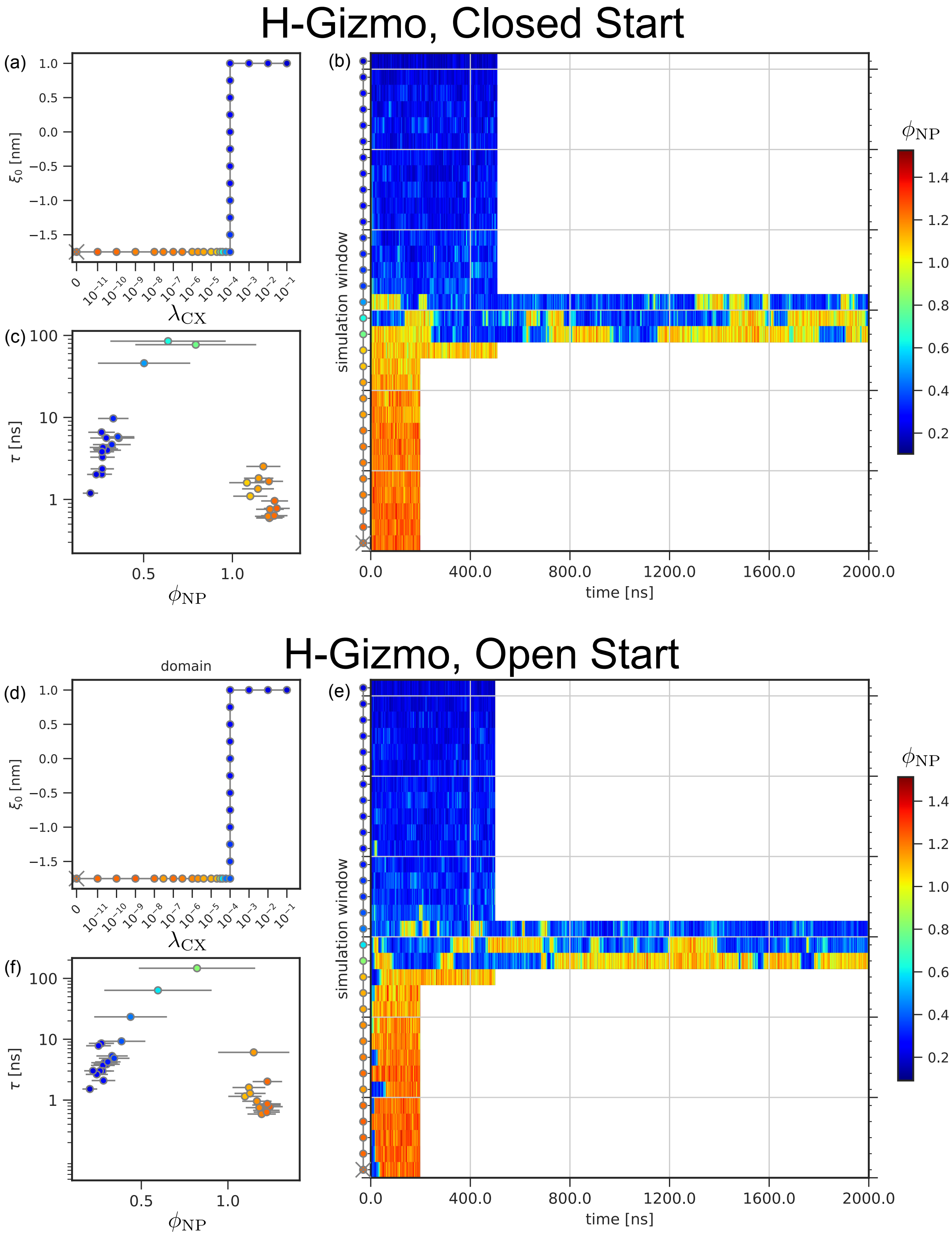


FIG. S2: Summary of the PMF ensembles using H-Gizmos and initiated from closed (a-c) or open (d-f) pore states. Panels (a) and (d) show the control parameters $\lambda_{\mathrm{CX}}$ and $\xi_0$ for the windows in the simulated ensemble. Panels (b) and (e) show a raster time series of $\phi_{\mathrm{NP}}$ for each simulation window. Panels (c) and (f) show scatter plots of autocorrelation times $\tau$ versus mean values for the $\phi_{\mathrm{NP}}$ timeseries, computed using data from 50ns onwards. The grey horizontal error bars indicate $\pm$ one standard deviation. The colorbars in (b) and (e) apply to the dots and rasters in all panels, and the dots are colored by the mean value of $\phi_{\mathrm{NP}}$.

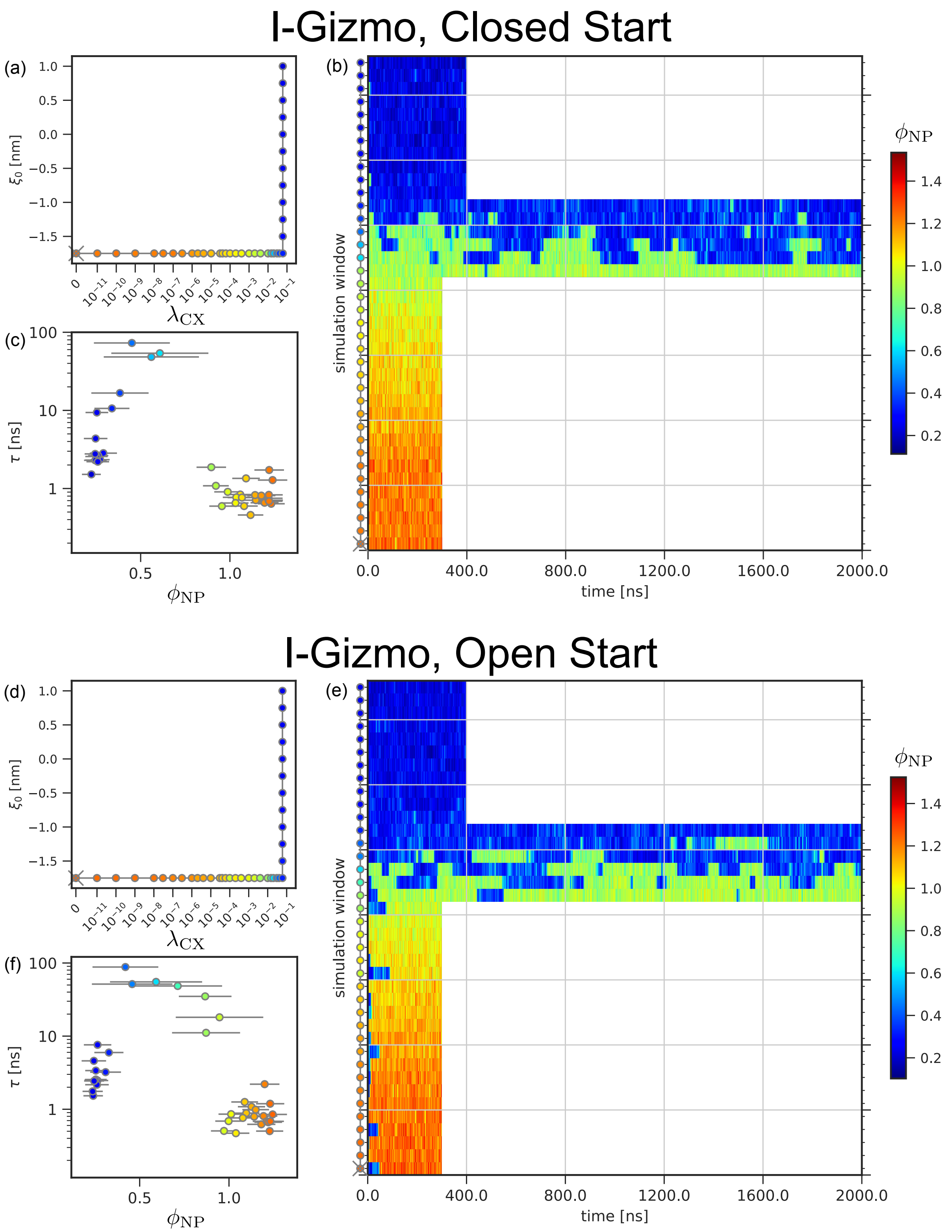


FIG. S3: The PMF ensembles using I-Gizmos and initiated from closed (a-c) or open (d-f) pore states. The plots are formatted as described in Fig S2

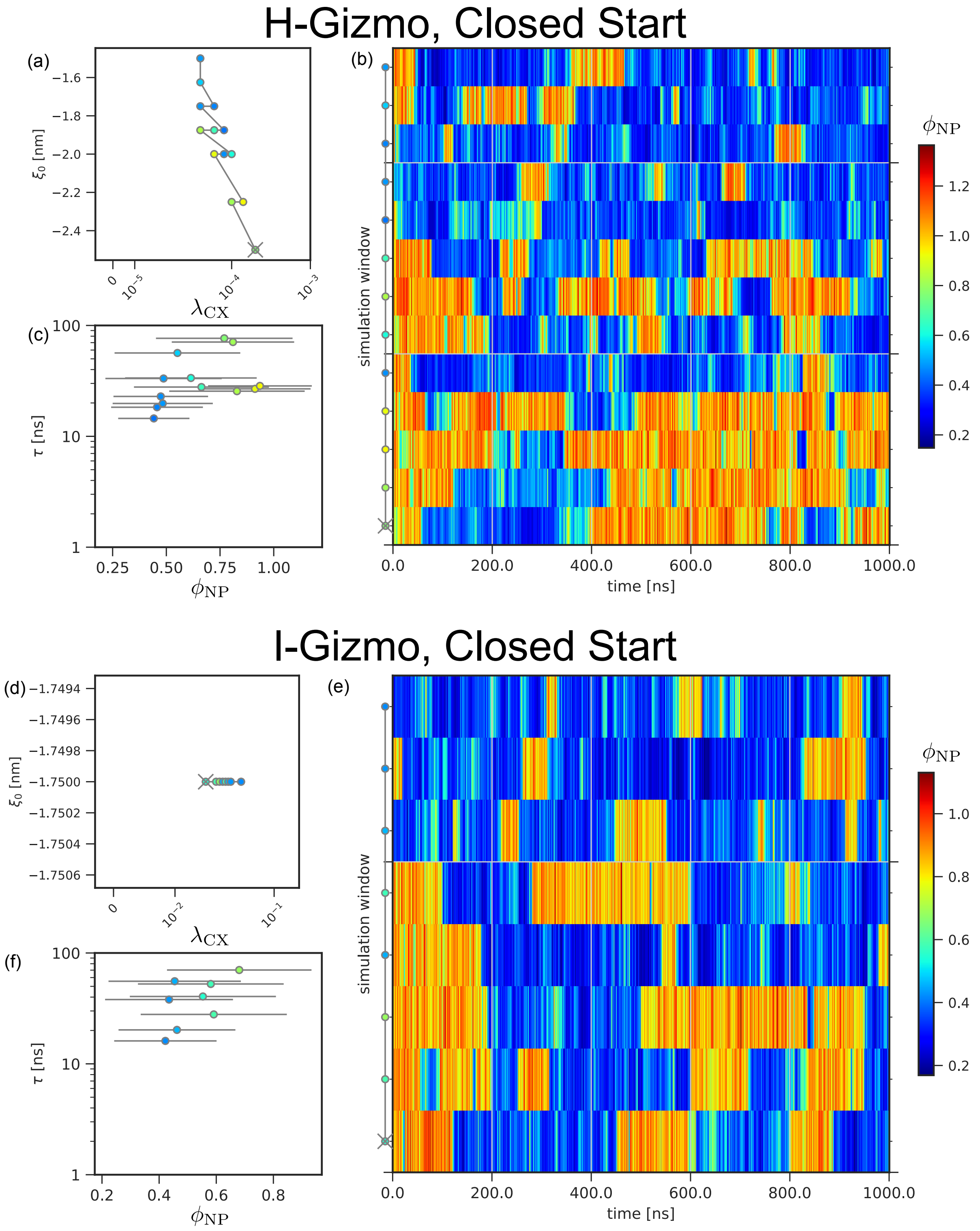


FIG. S4: The committor ensembles using H-Gizmos (a,b,c) and I-Gizmos (d,e,f), and initiated from closed pore states. The plots are formatted as described in Fig S2

### COMMITTOR DATASET

The committor datasets $\mathcal{C}^{(\mathrm{H})}$ and $\mathcal{C}^{(\mathrm{I})}$ were built by selecting structures close to the TS from the trajectories described above (see Fig. S4). Table S3 lists the parameters of these committor datasets.

The structure pool was grown in four iterations. In the first iteration, we made an initial (hand) estimate of the $c(\mathbf{x}) = 0.5$ dividing plane for the full simulation ensemble projected onto the 2D space of CVs $\mathrm{r}_{10}^{\mathrm{NP}}$ and $\mathrm{r}_{10}^{\mathrm{OW}}$. Then, we added to the structure pool: (i) any point within a threshold distance of the plane, (ii) any pair of time-adjacent points that crossed the plane and (iii) any points that were time-adjacent to the points in (i) and (ii). Then, $\hat{c}(\mathbf{x})$ estimates were computed for these structures (using $N_c = 8$ for the first iteration, and $N_c = 16$ afterwards). In later iterations we also included (iv) points that were time-adjacent to points with $0 < \hat{c}(\mathbf{x}) < 1$. Each subsequent iteration consisted of computing a new regression estimate $f_c = 0.5$, collecting points matching criteria (i-iv), and computing $\hat{c}(\mathbf{x})$ for the new points.

After the datasets $\mathcal{C}^{(\mathrm{H})}$ and $\mathcal{C}^{(\mathrm{I})}$ were built, and prior to computing the regression models for RC scoring, each committor dataset was split into training ($\mathcal{C}_{\mathrm{train}}$) and cross validation ($\mathcal{C}_{\mathrm{xval}}$) subsets by taking structures before and after a time cutoff. In addition, because the datasets had an excess of points with $\hat{c}(\mathbf{x}) = 0$ and 1, each dataset was sub-sampled to have an approximately flat distribution in $p(\hat{c}(\mathbf{x}))$. This prevented the $R^2$ values from disproportionately reflecting the fit at the $\hat{c}(\mathbf{x})$ extremes.

TABLE S3: Parameters of the committor datasets ($\mathcal{C}^{(\mathrm{H})}$ and $\mathcal{C}^{(\mathrm{I})}$) and their parent gizmo simulation ensembles.

| parameter | $\mathcal{C}^{(\mathrm{H})}$ | $\mathcal{C}^{(\mathrm{I})}$ |
|---|---|---|
| Windows in ensemble | 13 | 8 |
| Window length [ns] | 500 | 1000 |
| Time cutoff (train/xval)[ns] | 300 | 700 |
| Training set size | 694 | 378 |
| Validation set size | 505 | 176 |
| $\phi_{\mathrm{NP}}^{(\mathrm{A})}$ (closed) | 1.2 | 1.2 |
| $\phi_{\mathrm{NP}}^{(\mathrm{B})}$ (open) | 0.35 | 0.35 |

To compute $\hat{c}(\mathbf{x}, N_c)$ for a given $\mathbf{x}$, we used $N_c$ velocity re-seeded simulations ("shots"), each 6ns long, with the gizmo-system coupling switched fully off. Each shot was projected onto $\phi_{\mathrm{NP}}$ and absorbing boundaries at $\phi_{\mathrm{NP}}^{(\mathrm{A})}$=1.2 and $\phi_{\mathrm{NP}}^{(\mathrm{B})}$=0.35 were used to classify the shot as having reached end state $A$ (closed) or $B$ (open). Shots that reached neither absorbing boundary within 6ns (15.7% and 13.2% of all shots for the H- and I-gizmo datasets) were not considered for computing $\hat{c}(\mathbf{x}, N_c)$.

### COMMITTOR REGRESSION MODEL SCORES

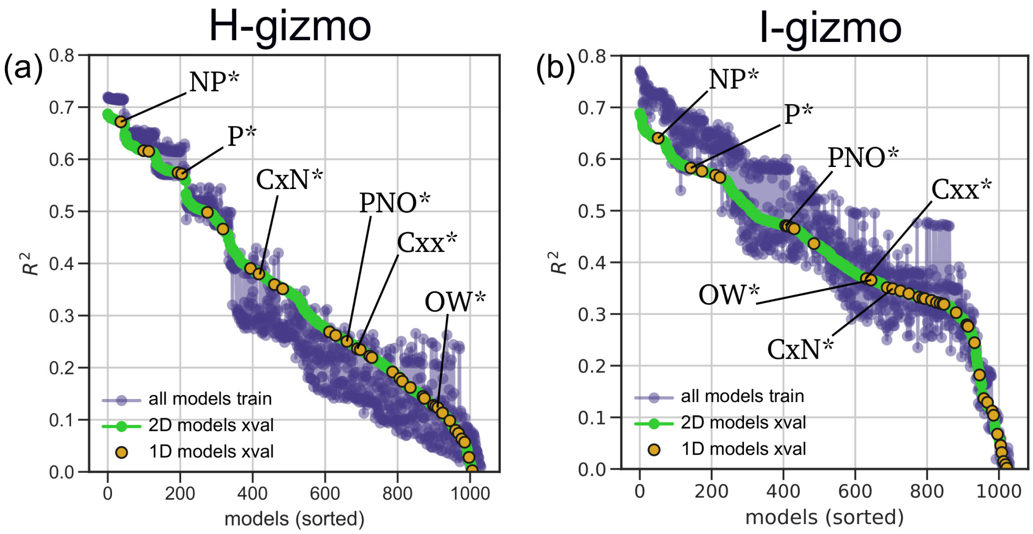


FIG. S5: Comparison of committor regression model scores $R^2$ computed using the $\mathcal{C}^{(\mathrm{H})}$ (a) and $\mathcal{C}^{(\mathrm{I})}$ (b) datasets. Both 1D and 2D models are shown and are co-sorted. The labeled orange dots indicate the top scoring 1D model for each atom group.


*

†



[1] S. Kumar, J. M. Rosenberg, D. Bouzida, R. H. Swendsen, and P. A. Kollman, Journal of Computational Chemistry **13**, 1011 (1992).

[2] J. D. Chodera, W. C. Swope, J. W. Pitera, C. Seok, and K. A. Dill, Journal of Chemical Theory and Computation **3**, 26 (2007).

[3] Z. Tan, E. Gallicchio, M. Lapelosa, and R. M. Levy, The Journal of Chemical Physics **136**, 144102 (2012).

[4] O. Berger, O. Edholm, and F. Jähnig, Biophysical Journal **72**, 2002 (1997).

[5] C. Oostenbrink, A. Villa, A. E. Mark, and W. F. van Gunsteren, Journal of Computational Chemistry **25**, 1656 (2004).

[6] S. Kawamoto and W. Shinoda, Soft Matter **10**, 3048 (2014).

[7] M. R. Shirts and J. D. Chodera, The Journal of Chemical Physics **129**, 124105 (2008).
