## Supplementary material for "Sequential water and headgroup merger: Membrane poration paths and energetics from MD simulations": pore-gizmo.zip: bpj.tex

\documentclass[lineno,twocolumn,endfloat,biblatex]{biophys-new}
% \documentclass{biophys}
\usepackage[utf8]{inputenc}
\usepackage{graphicx}
\usepackage[colorlinks,allcolors=cyan!70!black]{hyperref}

% Uncomment if using biblatex
% \addbibresource{sample.bib}

\usepackage{lipsum}

\title{New Biophysical Journal Template}
\runningtitle{Biophysical Journal Template} %% For page header

\author[1,*]{First author}
\author[2]{Second author}
\runningauthor{Author1 and Author2} %% For page header

\affil[1]{Institution A, Address A}
\affil[2]{Institution B, Address B}

\corrauthor[*]{}

% \papertype{Letters}
\papertype{Article}
% \papertype{Computational Tools}


\begin{document}

\begin{frontmatter}

\begin{abstract}
Each manuscript must be accompanied by an informative abstract of no more than 300 words. Abstracts should describe the substance of the manuscript in language non-specialists can understand, and must make clear the biological significance of the research. Reference citations are not allowed in the Abstract of a manuscript. Computational Tools and Letters articles are limited to five pages in length.
\end{abstract}

\begin{sigstatement}
Each manuscript must also have a statement of significance or no more than 120 words. Each manuscript must also have a statement of significance or no more than 120 words. Each manuscript must also have a statement of significance or no more than 120 words.
\end{sigstatement}
\end{frontmatter}

\section*{Introduction}


Thanks for using Overleaf to write your article. Your introduction goes here! Some examples of commonly used commands and features are listed below, to help you get started. Leave a blank line between blocks of text to start a new paragraph---use \verb|\\| for separating tabular rows or hard line-breaks only. Abbreviations should be defined in the text at first mention.

Please also take note of the \verb|\section*{...}| titles in this template: they are the required sections in a regular research Article manuscript. 

In particular, the main text of regular Articles and Computational Tools manuscripts must be structured with the following sections: \textbf{Introduction}, \textbf{Materials and Methods}, \textbf{Results}, \textbf{Discussion (or Results and Discussion)}, \textbf{Conclusion}.

Theoretical manuscripts may include just a \textbf{Methods} section and do not require \textbf{Materials}.

No particular organization structure is required for Letters.

If your manuscript is accepted, the Biophysical production team will re-format the references for publication. \emph{It is not necessary to format the reference list yourself to mirror the final published form.}

\section*{Materials and Methods}

Capitalize trade names and give manufacturers' full names and addresses (city and state). 


\subsection*{Sectioning commands}

Use \verb|\section*{...}| and \verb|\subsection*{...}| to create first- and second-level headings. Sed ut perspiciatis unde omnis iste natus error sit voluptatem accusantium doloremque laudantium, totam rem aperiam, eaque ipsa quae ab illo inventore veritatis et quasi architecto beatae vitae dicta sunt explicabo. 

\subsection*{Figures and Tables}

Use the table and tabular commands for basic tables --- see Table \ref{tab:widgets}, for example. \href{http://tablesgenerator.com}{TablesGenerator.com} is a handy tool for designing tables and generating the \LaTeX{} \texttt{tabular} code, which you can copy and paste into your article here.

You can upload a figure (JPG, PNG or PDF) using the PROJECT menu (Files\ldots > Add files). To include it in your document, use the \verb|graphicx| package and the \verb|\includegraphics| command as in the code for Figure \ref{fig:view}. 

In addition, you can use \verb|\ref{...}| and \verb|\label{...}| commands for cross-references.

\begin{table}[hbt!]
\caption{An example table}
\label{tab:widgets}
\centering

\begin{threeparttable}

\begin{tabular}{c l r}
\hline
Code & Item & Quantity \\\hline
W1 & Widgets\tnote{a} & 42 \\
G35 & Gadgets & 13\tnote{b} \\
\hline
\end{tabular}

\begin{tablenotes}
\item[a] This is a table note.
\item[b] This is another table note.
\end{tablenotes}

\end{threeparttable}

\end{table}

\begin{figure}[hbt!]
\centering
\includegraphics[width=0.6\linewidth]{example-image}
\caption{A figure example.}
\label{fig:view}

\end{figure}

\section*{Results}

\LaTeX{} is great at typesetting mathematics:

Let $X_1, X_2, \ldots, X_n$ be a sequence of independent and identically distributed random variables with $\text{E}[X_i] = \mu$ and $\text{Var}[X_i] = \sigma^2 < \infty$, and let
\begin{equation}
\label{eq:CLT}
S_n = \frac{X_1 + X_2 + \cdots + X_n}{n}
      = \frac{1}{n}\sum_{i}^{n} X_i
\end{equation}
denote their mean. Then as $n$ approaches infinity, the random variables $\sqrt{n}(S_n - \mu)$ converge in distribution to a normal $\mathcal{N}(0, \sigma^2)$. Thus concludes the explanation about Eq.~\ref{eq:CLT}.


You can make lists with automatic numbering \dots

\begin{enumerate}
\item Like this,
\item and like this.
\end{enumerate}

\dots or bullet points \dots

\begin{itemize} 
\item Like this,
\item and like this.
\end{itemize}

\dots or with words and descriptions \dots

\begin{description}
\item[Word] Definition
\item[Concept] Explanation
\item[Idea] Text
\end{description}

An example quotation:

\begin{quote}
Lorem ipsum dolor sit amet, consectetur adipiscing elit, sed do eiusmod tempor incididunt ut labore et dolore magna aliqua. Ut enim ad minim veniam, quis nostrud exercitation ullamco laboris nisi ut aliquip ex ea commodo consequat.
\end{quote}


\section*{Discussion}

\LaTeX{} formats citations and references automatically using the bibliography records in your .bib file, which you can edit via the project menu. Use the \verb|\cite| command to insert a citation, like this: \cite{Chen_Nicholson00} Multiple citations can be given as \cite{Stiles_Bartol01,el-Kareh_etal93,Callaghan91}. You can use either BibTeX or biblatex: see the following subsections.

If your manuscript is accepted, the Biophysical production team will re-format the references for publication. \emph{It is not necessary to format the reference list yourself to mirror the final published form.}

\subsection*{Using bibtex} 
This is the default. Specify your \texttt{.bib} file with \verb|\bibliography{sample}| (the extension is unnecessary) near the end of your manuscript, where you want the references list to appear.

\subsection*{Using biblatex} 
Pass the \texttt{biblatex} option to the \verb|\documentclass| declaration, then specify your \texttt{.bib} file name in the \emph{preamble}: \verb|\addbibresources{sample.bib}| (the extension is necessary). Write \verb|\printbibliography| near the end of your manuscript where you want the references to appear.

\section*{Conclusion}

Sed ut perspiciatis unde omnis iste natus error sit voluptatem accusantium doloremque laudantium, totam rem aperiam, eaque ipsa quae ab illo inventore veritatis et quasi architecto beatae vitae dicta sunt explicabo. 

\section*{Author Contributions}

Author1 designed the research. Author2 carried out all simulations, analyzed the data. Author1 and Author2 wrote the article. 

\section*{Acknowledgments}

We thank G. Harrison, B. Harper, and J. Doe for their help.

% Uncomment if using bibtex (default)
\bibliography{sample}

% Uncomment if using biblatex
% \printbibliography

\section*{Supplementary Material}

An online supplement to this article can be found by visiting BJ Online at \url{http://www.biophysj.org}.

\end{document}
