## Supplementary material for "Sequential water and headgroup merger: Membrane poration paths and energetics from MD simulations": pore-gizmo.zip: cover.tex

%\documentclass[aps,amsmath,amssymb,floatfix,superscriptaddress]{revtex4-1} % APS journal style (preprint or reprint)
%\documentclass[a4paper,10pt]{letter}
\documentclass[a4paper,12pt]{letter}

\usepackage[top=3cm]{geometry}

%\nonstopmode            % plow through errors/notes during latex2pdf

%abstract:    communicating message of paper
%coverletter: communicating impact/relevance/interest of paper
% why BPJ? To address biological questions

\begin{document}
\noindent
Dear Editor

Please consider the enclosed manuscript "Water defects precede headgroup merger in membrane poration: Paths and energetics from Gizmo biased MD simulations" for publication in Biophysical Journal.

Small prepore defects in lipid membranes have been of interest for decades because their formation is a primary step of many important biological processes, such as synaptic transmission, fertilisation, virus or phage entry into the host cell, and vesicle mediated cellular transport processes. Despite ample experimental data, their nucleation mechanism is not fully understood at the atomic level. Molecular simulations have provided some insight, but are hampered largely by the lack of optimal mechanistic structural descriptors (reaction coordinates, RCs). This limitation, which, crucially, also precludes an accurate characterisation of the underlying energetics and kinetics via free energy calculations, in turn arises from the use of empirical RCs due to the absence of a systematic approach for their construction.

Here, we address this shortcoming and report new structural and energetics insight into the opening and closing of small pore defects. Using the ideal theoretical RC -- the committor -- to determine RCs that nearly optimally describe the poration mechanism allowed us to compute accurate free energies for the full preporation process. We quantitatively compare hydrophobic versus indentation mechanisms, resolve the sequential actions of water and head group penetration at the transition state, and report the energetic stability of the metastable prepore basin -- thereby explaining previous indirect experimental evidence.

Beyond the relevance of our results for the field of topological membrane transformations and membrane fusion, we think our study should be of broader interest to the readers of the Biophysical Journal not only because of the biological ubiquity of pore defects, but also because it provides a general framework for the widespread challenge of determining reaction coordinates and mechanisms for diffusive, permutationally frustrated systems.  

The following people would be suitable reviewers in our opinion, owing to their expertise in membrane physics and simulation methods.


\noindent
Wouter den Otter, Twente\\
Michael Klein, Temple\\
Peter Tieleman, Calgary\\
Drew Bennet, Lawrence Livermore National Lab\\
Mark Stevens, Sandia National Lab\\


\noindent 
Sincerely,\\

\noindent
Greg Bubnis\\
Helmut Grubm\"{u}ller


\end{document}
