## Supplementary material for "Sequential water and headgroup merger: Membrane poration paths and energetics from MD simulations": pore-gizmo.zip: main.tex

\documentclass[aps,amsmath,reprint,amssymb,prl,floatfix,superscriptaddress]{revtex4-1} % APS journal style (preprint or reprint)

%\documentclass[a4paper,12pt]{article}
%\documentclass[a4paper,10pt]{article}
\usepackage{graphicx}   % Include figure files
\usepackage{dcolumn}    % Align table columns on decimal point
\usepackage{bm}         % bold math
\usepackage{xcolor}     % textcolor
\usepackage{hyperref}   % add hypertext capabilities
%\usepackage{float}
\nonstopmode            % plow through errors/notes during latex2pdf

\newcommand{\wt}{\widetilde}
\newcommand{\wh}{\widehat}
\newcommand{\mb}{\mathbf}

%\usepackage[left=1.5cm, right=4cm, top=2cm, bottom=2cm]{geometry}

%\bibliographystyle{abbrv}
%\bibliographystyle{chem-acs}

%\bibliographystyle{apsrev4-1}


%\usepackage[backend=biber, style=phys,]{biblatex}
%\addbibresource{refs.bib}

\def\abt105{Department of Theoretical and Computational Biophysics, Max-Planck-Institute for Biophysical Chemistry, G\"{o}ttingen, Germany}
\def\weill{Weill Institute for Neurosciences and Department of Neurology, University of California San Francisco, San Francisco California 94158, USA}

\begin{document}
\title{Water defects precede headgroup merger in membrane poration: Paths and energetics from Gizmo biased MD simulations}

\author{Greg Bubnis}
\affiliation{\abt105}
\affiliation{\weill}
\author{Helmut Grubm\"{u}ller}
\affiliation{\abt105}
\date{\today}

\begin{abstract}
Membrane topology changes such as poration, stalk formation, and hemi-fusion rupture are essential to cellular function, but their molecular details, energetics, and kinetics are still not fully understood. Here we present a unified energetic and mechanistic picture of metastable pore defects in tensionless lipid membranes. We used an exhaustive committor analysis to test and select optimal reaction coordinates and also to determine the nucleation mechanism. These reaction coordinates were used to calculate free energy landscapes that capture the full process and end states. The identified barriers agree with the committor analysis. To enable sufficient sampling of the complete transition path for our atomistic simulations, we developed a novel "gizmo" potential biasing scheme. The simulations suggest that the essential step in the nucleation is the initial merger of lipid headgroups at the nascent pore center. To facilitate this event, an indentation pathway is energetically preferred to a hydrophobic defect. Continuous water columns that span the indentation were determined to be on-path transients that precede the nucleation barrier. This study gives a quantitative description of the nucleation mechanism and energetics of small metastable pores and illustrates a systematic approach to uncover the mechanisms of diverse cellular membrane remodeling processes.
\end{abstract}

\maketitle

\section{Statement of Significance}
The primary steps and nucleation of lipid membrane pore formation are key to membrane fusion, viral infection, and vesicular cellular transport. Despite decades experimental and theoretical studies, the underlying mechanisms are still not fully understood at the atomic level. Using a committor-based reaction coordinate and atomistic simulations, we report new structural and energetics insight into the full poration process.  We find that the pore nucleates via an elastic indentation rather than by forming a hydrophobic defect. Subsequently, water pierces the thinned slab as a prerequisite for the following axial merger of the first lipid headgroups from opposite monolayers, which precedes and best characterizes the transition state. We also identify a metastable prepore basin, thereby explaining previous indirect experimental evidence.

%========================================================
\section{Introduction}
%========================================================
Lipid membranes can undergo a number of topological remodelling processes --- such as endo- and exocytosis, vesiculation, viral entry, and fertilization --- that involve the formation or closure of aqueous pore defects. Pore nucleation is key to transmembrane transport \cite{neumann_gene_1982}, ion permeation \cite{deamer_permeability_1986, fathizadeh_ion_2019}, antimicrobial peptide function \cite{reddy_antimicrobial_2004, shai_mode_2002}, and bilayer equilibration \cite{gurtovenko_molecular_2007, bruckner_flip-flop-induced_2009} and is the essential step of synaptic transmission \cite{burger_greasing_2000, jahn_membrane_1999, risselada_free_2014}. Despite the evidence for small, metastable aqueous pores from conductivity \cite{abidor_electric_1979} and tension \cite{evans_dynamic_2003} experiments, as well as molecular simulations \cite{ting_metastable_2018, hub_probing_2017, leontiadou_molecular_2004}, the atomistic details, driving forces, and kinetics of their formation and closure are not entirely understood \cite{kirsch_membrane_2016, awasthi_simulations_2016}.

In the long established continuum description for pores, namely the Litster model \cite{litster_stability_1975}, a pore's energy is governed by the membrane surface tension and the pore rim line tension, both of which depend on the pore radius. This model is suited for already-formed pores but does not explain the large energies required to create the topological defect, nor can it describe metastable pore defects in the absence of applied tension \cite{leontiadou_molecular_2004, ting_metastable_2018, evans_dynamic_2003, abidor_electric_1979}. In a more recent continuum description, a nascent pore is treated explicitly as a hydrophobic cylinder of solvent-exposed lipid tails, with both a height and radius \cite{akimov_pore_2017}. Here we focus on the formation of metastable prepores (henceforth pores) that arise in a tensionless bilayer, using atomistic molecular dynamics (MD) simulations to fully resolve the process.

Specifically, we will address the following questions. Is the poration pathway hydrophobic, with a penetrating water column surrounded by lipid tails, or hydrophilic, involving an indentation where lipid head groups submerge to shield water/tail interactions? In fact, "water wires" that span the hydrophobic slab have been observed in several recent simulation studies \cite{bennett_atomistic_2014, grafmuller_free_2014, ting_metastable_2018}, but whether or not this step is rate-limiting is unclear. In one study, the energetics of pore formation were found to be insensitive to "bundling" four waters together, suggesting that the precise organisation of water matters little \cite{bennett_water_2011}. We will therefore address the question of whether or not these water wires are the energetic transition state, and whether single file water columns are sufficient to nucleate the pore. To address these questions, we aim to identify optimal reaction coordinates (RCs) in terms of collective variables (CVs) to properly describe the progress and energetics of the nucleation mechanism. (The terms CV and RC are both used throughout this study, with the intention that CV indicates a collective coordinate that may or may not be a useful RC, whereas a RC is tasked to measure reaction progress)

Whereas spontaneous pores can form on $\mu s$ timescales for simulations of short-tailed lipids \cite{bennett_atomistic_2014}, biasing potentials acting along a chosen RC are usually required to access relevant structural intermediates, in particular the transition state (TS). Typically an {\em ad hoc} RC $\xi(\mathbf{x})$ --- defined as a function of the combined coordinate vector $\mathbf{x}$ of a suitable set of relevant atoms ---  is used as a biasing coordinate to compute a free energy profile (or potential of mean force, PMF) $G(\xi)$ via umbrella sampling or related methods \cite{torrie_nonphysical_1977, smirnova_free-energy_2015}. The proper choice of RCs is therefore critical for obtaining correct and well-converged energetics, and also serves to characterise the reaction mechanism in structural terms. Indeed, for diverse membrane remodeling processes (stalk nucleation, vesiculation, hemifusion) the fluid disorder of the membranes makes it far from obvious which collective motions (i.e. RCs) are best suited as mechanistic descriptors and for biasing simulations towards transient intermediates.

%the limitations of most established RCs have been documented.. 
A recent comparison of poration RCs documented that many established and intuitively plausible RCs for poration were poor biasing coordinates due to prohibitively slow convergence and resulting hysteresis effects \cite{awasthi_simulations_2016}. Further, and quite generally, even for well-converged simulations, sub-optimal choice of RCs may artificially lower --- or even hide --- important energy barriers if different intermediates of the reaction progress are projected onto similar values along the RC \cite{kopelevich_one-dimensional_2013, fathizadeh_ion_2019}. One consequence is that in such cases, the TS region may not be properly described by the RCs, and the position of the obtained PMF barrier may differ markedly from the true TS.  

More recently, a poration RC based on the fraction of cylindrical slabs (spanning the membrane) occupied by hydrophilic particles (water, lipid headgroups) has been proposed, which enabled converged free energy calculations suggesting a free energy barrier and TS leading to the metastable (pre)pore state \cite{hub_probing_2017}. Despite this advance, the observed barrier was sensitive to several tuning parameters, such that it remained unclear which combination provides an accurate result, and the subsequent pore could not be well resolved.

% more general studies of optimal reaction coordinates 
For a more systematic approach, we exploit the optimal RC for any conformational transition from a conformational state $A$ to a state $B$, which is  --- by definition --- the committor $c(\mathbf{x}) \equiv P(B|\mathbf{x})$, i.e., the probability that an unrestrained trajectory seeded from $\mathbf{x}$ reaches state B before state A (cf.\ Fig.~\ref{figure:FIG-02-ts-schematic}b). This concept traces back to Onsager \cite{onsager_initial_1938} and has been more recently reviewed \cite{li_recent_2014, peters_reaction_2016, banushkina_optimal_2016}. In this framework, $c(\mathbf{x})$ optimally reflects the reaction progress, and all configurations $\mathbf{x}$ with $c(\mathbf{x})=0.5$ comprise the TS between the basins of attraction of $A$ and $B$. Computing $c(\mathbf{x})$ is, however, often computationally prohibitive, as it involves starting sufficiently many trajectories from every point $\mathbf{x}$ along the transition path ensemble. In addition, $c(\mathbf{x})$ is not an intuitive structural descriptor and thus offers no structural insight by itself. The real utility of $c(\mathbf{x})$ stems from the fact that a good RC must be a strong committor correlate in the TS region. It follows then that putative RCs can be quantitatively compared by how well they correlate to $c(\mathbf{x})$.

Indeed, this concept has been applied to several other systems \cite{ma_automatic_2005, peters_obtaining_2006, quaytman_reaction_2007, du_transition_1998} as well to as a highly coarse grained lipid system \cite{marti_molecular_2004}. For these systems, RCs were successfully identified which correlated strongly with $c(\mathbf{x})$ and, hence, were considered close to optimal. Notably, once the expensive computation of $c(\mathbf{x})$ is achieved for a sufficient number of points, any number of candidate RCs can be evaluated and ranked via  post-processing at only little computational cost, and without further simulations. The appeal of this strategy is that, rather than relying on intuition, the selection of an optimal RC becomes an optimisation problem with a defined cost function. This framework also allows for building more powerful descriptors via linear \cite{peters_obtaining_2006} or non-linear \cite{ma_automatic_2005} combinations of the initial CV pool, although overly complex hybrid coordinates (e.g. derived via a deep neural network), bear the risks of overfitting and of becoming uninterpretable. Yet, for a very broad range of systems, the approach provides a principled method to identify RCs that are sufficiently close to optimal and thus provide quantitative insight into reaction paths and energetics.

Here we apply such a committor-driven scheme to study pore nucleation in a solvated DMPC lipid bilayer, which is known to form metastable prepores \cite{bennett_water_2011, awasthi_simulations_2016}. To efficiently sample states where $c(\mathbf{x})\approx 0.5$, we will first develop a novel biasing scheme using a membrane embedded "gizmo" which energetically biases the system towards intermediate states and functionally resembles a lipid scramblase \cite{kalienkova_stepwise_2019, nakao_effects_2018}. Using gizmo biasing, multi-$\mu$s simulations will be used to sample TS-crossing events as input for subsequent (unbiased) $c(\mathbf{x})$ calculations. The obtained $c(\mathbf{x})$ estimates will then be used as a regression target to score and rank putative RCs from a diverse combinatoric pool including CVs for water, lipid head groups, and lipid tails, as well as pairwise combinations thereof. A second set of gizmo biased simulations will serve to sample the full poration pathway and recover unbiased PMFs, projected onto optimal RCs determined from the committor analysis. Taken together, the PMF along the optimal RC will serve to address the above mechanistic questions.

%========================================================
\section{Theory}
%========================================================
\subsection{Biasing Potential}
%========================================================
To thoroughly sample the poration process, and in particular the TS, we have developed a pseudomolecular "gizmo" embedded within the membrane, much like (but not intended to resemble) a membrane protein. This gizmo imposes a potential energy bias onto adjacent lipid and water molecules so as to direct the membrane towards open, transition, or closed pore states.

%==== Fig. 1: gizmo designs
\begin{figure*}[htbp] %\centering
\includegraphics[width=6.25in]{fig-01-gizmos.eps}
\caption{\small{(color online) Pore gizmo designs. (a) A vertical chain of (red) CX particles drives pore opening by creating a packing defect in the hydrophobic slab. (b) A belt of (green) WX particles drives pore closure by pinching off the water column. The belt is drawn with six WX particles for clarity, whereas the actual belt has twelve. The faces in (a-b) indicate in which compartments the CX/WX particles are non-interacting (smiley) and purely repulsive (frowns), such that these particles can occupy one bulk region of the system without introducing any perturbation. (c-d) When combined to form a gizmo, the CX chain and WX belt allow reversible control of pore formation. (c) The "hydrophobic" H-gizmo uses a chain of 11 CX beads to create a hydrophobic packing defect spanning the bilayer. (d) The "indentation" I-gizmo has its three centermost beads switched off so that two partial chains penetrate the membrane from both sides, creating an indentation defect.}}
\label{figure:FIG-01-gizmo}
\end{figure*}

To this aim, the gizmo (sketched in Fig.~\ref{figure:FIG-01-gizmo} and fully described in the Supporting Material and Fig~.S1) was designed with two special-purpose structural elements for driving pore opening and closure, respectively. To drive pore opening, we used a "chain" of lipid-tail repellent (CX) beads (red) that is aligned to the bilayer normal to create a packing defect in the hydrophobic region (light grey) of the membrane (Fig.~\ref{figure:FIG-01-gizmo}(a)). To drive closure, we used a circular "belt" of water repellent (WX) beads (green) lying in the membrane midplane to constrict and pinch off the water column (blue) (Fig.~\ref{figure:FIG-01-gizmo}(b)). The chain and belt are flexible and are bound to a stiff frame comprised of ghost particles (GG) which interact via bonded interactions, but not with any other particles of the simulation system (i.e., no van der Waals or electrostatic interactions).

The gizmo was implemented via the potential energy of the combined system, 
\begin{equation}
\label{eq_potential}
\tilde{V}(\mathbf{x,y}) = V_x(\mathbf{x}) + V_{xy}(\mathbf{x,y}) + V_y(\mathbf{y}) + V_{\xi}(\xi(\mathbf{y})),
\end{equation}
where $V_x(\mathbf{x})$ is the force field for the lipids and solvent system, $V_y(\mathbf{y})$ that for the gizmo, $V_{xy}(\mathbf{x,y}) = \lambda_{\text{CX}} V_{\text{CX}}(\mathbf{x,y}) + \lambda_{\text{WX}} V_{\text{WX}}(\mathbf{x,y})$ couples the system and gizmo via  repulsive CX/WX interactions, and a harmonic restraint $V_{\xi}(\xi(\mathbf{y})) = \frac{1}{2}k( \xi(\mathbf{y}) - \xi_0 )^2$ controls the WX belt radius of the gizmo with a collective radial breathing coordinate $\xi$. In the above equations,  $\mathbf{x}$ denotes the (combined) atomic coordinates vector of the lipids and solvent, and $\mathbf{y}$ the (combined) atomic coordinates vector of the gizmo. 

The repulsive potentials $V_{\text{CX}}$ and $V_{\text{WX}}$ were implemented via Lennard-Jones potentials ($\sigma$=$1.1\,$nm and $\epsilon$=0.01\,kJ/mol). The tilde over $\tilde{V}(\mathbf{x,y})$ indicates that the lipid-solvent system is coupled to, and biased by, the gizmo. The relative strengths of the gizmo's chain and belt biases were tuned via $\lambda_{\text{CX}}$, which scales the CX-tail repulsion that drives opening, and $\xi_0$, which controls the WX belt radius via a harmonic restraint that is constricted to drive closure ($\lambda_{\text{WX}}$ was fixed to 1). Crucially, except for these biasing potential contributions (i.e. $V_{xy}$), the beads do not interact with the physical system of lipids and solvent (see Fig.~\ref{figure:FIG-01-gizmo}). Accordingly, the potential energy remains otherwise unperturbed and the gizmo potential can be instantly switched off without creating packing artefacts, as is required for the committor simulations. This setup also enabled us to derive PMFs for pore opening and closure for the lipid-solvent system from the biased MD ensemble as described in the Supporting Material. 

The gizmo has a few additional features worth noting. First, in the open state, it functions as a lipid scramblase, stabilizing a pore indefinitely and allowing lipids to flip-flop and equilibrate. Also, due to its shape and hydrophobicity, the WX belt keeps the gizmo embedded and properly oriented in the membrane interior. Lastly, the gizmo's central atom (a GG atom) provides a convenient reference position for the pore center, which is useful for deriving localized CVs for the lipids and solvent.

We used two different gizmo potentials to efficiently sample two different pore formation paths. The "hydrophobic" H-gizmo (Fig.~\ref{figure:FIG-01-gizmo}(c)) promotes a hydrophobic defect --- where a narrow water column pierces and hydrates the hydrophobic slab and then head groups pivot inwards --- by a chain of 11 CX beads which creates a tail packing defect fully spanning the membrane. In contrast, the "indentation" I-gizmo (Fig.~\ref{figure:FIG-01-gizmo}(d)) drives indentation and thinning of the bilayer without directly perturbing lipid chain packing at the bilayer midplane. This effect is achieved by a chain that has its three centermost CX beads replaced by GG beads (acting as spacers) such that two CX chain segments, with four beads each, penetrate from opposite sides.  The two gizmos also served to assess how much the resulting RCs and PMFs depend on the particular properties of the gizmos.

We used independent ensembles of gizmo biased simulations to collect input structures for committor analysis and to sample the full poration pathway for PMF calculations. As summarized in Table \ref{TAB00}, these ensembles used different initial structures to test convergence, and different gizmo types to allow us to compare hydrophobic and indentation mechanisms. For all of the computed ensembles, we ensured sufficient TS sampling by using extended simulations (up to 2 $\mu$s when necessary) targeted to the TS region. This ensured that the PMF calculations are well converged, irrespective of starting structure, and that a sufficiently large number of statistically uncorrelated starting structures is available in the vicinity of the TS for the 
subsequent committor analysis. Full details of these ensembles are provided in the Supporting Material (see Figs.~S2-4).

\begin{table}[htbp] 
\centering
   \caption{\small{Simulation ensembles used in this study}}
   \label{TAB00}
   \begin{tabular}{l|l|l}
    \hline
    Ensemble type & Gizmo & Starting state\\
    \hline
    PMF  & H & A (closed)\\
    PMF  & H & B (open)\\
    PMF  & I & A (closed) \\
    PMF  & I & B (open)\\
    Committor & H & A (closed)\\
    Committor & I & A (closed)\\
    \hline
   \end{tabular}
\end{table}

%========================================================
\subsection{Collective variables to describe pore formation}
%========================================================
To obtain structural insight into how lipids and solvent reorganize when crossing the TS, we assembled a set of CVs that capture diverse structural changes during pore nucleation. The pool of CVs was chosen to quantify the merger and depletion of specific atom groups $\alpha$ (see Tab.~\ref{TAB01}) relative to the pore center, defined by the center $\mathbf{y}_0$ of the gizmo (i.e. providing a local coordinate system). To study how different groups of atoms collectively reorganize, we defined atom groups that combine and isolate individual components of the system (e.g., lipid head groups with and without water oxygen). To compare the effects of local versus non-local collective motions, we varied the cutoff number $N$ of atoms closest to $\mathbf{y}_0$ used to compute the CVs. Finally, to distinguish different (an)-isotropic symmetries, we implemented isotropic, axial (z-axis), and lateral (xy plane) collective coordinates.

The isotropic mean radius $\text{r}^{\alpha}_N$ was computed as
\begin{eqnarray}
\label{rgop}
\text{r}^{\alpha}_N = \frac{1}{N}\sum_{i=1}^{N} || \mathbf{x}_i^\alpha - \mathbf{y}_0 || ,\
\end{eqnarray}
where $\mathbf{x}_i^\alpha$ is the position of atom $i$ belonging to the atom group $\alpha$.

To measure lateral merger or spreading of atoms, the 'xy' variant $\text{rxy}^{\alpha}_N$ was computed via
\begin{eqnarray}
\label{rgxyop}
\text{rxy}^{\alpha}_N = \frac{1}{N}\sum_{i=1}^{N} || \mathbf{x}_i^\alpha - \mathbf{y}_0 ||_\text{xy} ,\
\end{eqnarray}
where $||..||_\text{xy}$ denotes the distance between points projected onto the xy plane.

The axial variant $\text{rz}^\alpha_N$ was computed by projecting and sorting atoms on the membrane-normal z-axis, to capture the penetration of the lipid slab. When all system coordinates are projected onto the z-axis, thermal fluctuations can create non-local projection artefacts,  giving the false appearance of membrane penetration. These include "hanging droplets" (described in \cite{hub_probing_2017}), where two laterally distant indentations partially penetrate the bilayer, and low-wavenumber thermal undulations \cite{kopelevich_one-dimensional_2013}. To avoid these effects, we preselected the 100 atoms closest to $\mathbf{y}_0$ prior to projection and sorting.

From this set, $\text{rz}^\alpha_N$ was computed as the maximum over $N$-tuples (i.e., a set of $N$ successive, z-sorted atoms) of the average axial distance between the atoms and their mean position
\begin{eqnarray}
\label{rgzop}
\!\!\!\!\!\text{rz}^{\alpha}_N = \!\!\max_{j \in 1,\ldots,101-N} \left( \frac{1}{N}\sum_{i=j}^{j+N-1} || \mathbf{x}_i^\alpha - \bar{\mathbf{x}}_{j,N}^\alpha  ||_\text{z} \right) ,\
\end{eqnarray}
where $||..||_\text{z}$ denotes the distance between points projected onto the z-axis and $\bar{\mathbf{x}}_{j,N}^\alpha = N^{-1}\sum_{i=j}^{j+N-1} \mathbf{x}_i^\alpha$ is the moving average (of width $N$). This CV measures the largest gap (depletion) in the vertical column, without requiring the gap to be centered at any reference ($\mathbf{y}_0$ or bilayer midplane) and will therefore detect a gap that is centered above or below the bilayer midplane. 

%$\big< asdf \big>$
\begin{table}[htbp] 
\centering
   \caption{\small{Atom subsets ($\alpha$) used for r, rxy, and rz CVs.}}
   \label{TAB01}
   \begin{tabular}{l|l}
    \hline
    group label ($\alpha$) & group description \\
    \hline
    OW  &  water oxygens                 \\
    P   &  lipid phosphorus              \\
    NP  &  lipid nitrogen and phosphorus \\
    PNO &  NP $\cup$ OW                   \\
    CxN &  (terminal) lipid tail carbons \\
    Cxx &  (all) lipid carbons           \\
    \hline  
   \end{tabular}
\end{table}

%========================================================
\subsection{Committor regression models}
%========================================================
%==== Fig. 2: committor regression
\begin{figure}[htbp] %\centering
\includegraphics[width=0.8\columnwidth]{fig-02-ts-schematic-vertical.eps}
\caption{\small{Schematics showing (a) a commitor regression model $f_c$ which consists of a linear combination of input features with a logistic activation. (b) A 2D TS model $\mathcal{T}_{(\phi_1, \phi_2)}$ for the CVs $\phi_1(\mathbf{x})$ and  $\phi_2(\mathbf{x})$. The two free energy basins A and B are separated by the $f_c$=0.5 isoline. The three-point MEP estimate $\bm{\phi}^{\text{MEP}} = [\bm{\phi}^{0.2}, \bm{\phi}^{0.5}, \bm{\phi}^{0.8}]$ is drawn as cyan, red, and blue dots. The solid black line is a linear fit to $\bm{\phi}_{\text{MEP}}$, constrained to pass through $\bm{\phi}^{0.5}$, and the closed and open circles at the ends indicate the directions of the closed and open pore states, respectively. Because $a$ is a linear function of input CVs, all $f_c=const$ isolines are parallel.}  }
\label{figure:FIG-02-ts-schematic}
\end{figure}

A committor regression scheme was used to find optimal RCs among the pool of the just described CVs and to build low-dimensional TS models.

Given a configuration $\mathbf{x}$, a committor estimate $\hat{c}(\mathbf{x}|N_c)$ was computed from $N_c=16$ unrestrained (i.e., without gizmo potential), velocity-randomized trajectories of $6\,$ns length, each spawned from $\mathbf{x}$. By counting how many trajectories reached the A and B basins first ($N_A$ and $N_B$, respectively), we obtained $\hat{c}(\mathbf{x}) = N_B/(N_A+N_B)$; trajectories that reached neither boundary were discarded.   We denote a dataset of configurations and their committor estimates as $\mathcal{C}$ = $\left \{ (\mathbf{x}, \hat{c}(\mathbf{x}|N_c)) \right \}$.

To predict $\hat{c}(\mathbf{x})$ given CVs $(\phi_1(\mathbf{x}), \phi_2(\mathbf{x}), \ldots)$, we fit a logistic regression model $f_c$ (cf.\ Fig.~\ref{figure:FIG-02-ts-schematic}(a)),
\begin{eqnarray}
\label{linearFit}
\hat{c}(\mathbf{x}_i) = f_c(\mathbf{x}_i) + \epsilon_i = \frac{1}{1+e^{-a(\mathbf{x}_i)}} + \epsilon_i\,,
\end{eqnarray}
% NOTE: \hat{c} is an imperfect target since c = \hat{c} + error
where $a(\mathbf{x}_i) = \sum_{j}^{} w_j \phi_j(\mathbf{x}_i)$ is a linear combination of CVs (including a constant offset $\phi_0=1$) and $\epsilon_i$ is the residual error associated with $\mathbf{x}_i$. The logistic function was chosen because $\hat{c}(\mathbf{x})$ is expected to have a  sharp transition between its limiting values 0 and 1. The weights for each model $(w_0, w_1, ...)$ were fit by gradient descent to minimise the mean squared error, $\text{MSE} = \left< \epsilon_i^2 \right>$, for a training dataset $\mathcal{C}_\text{train}$ (equivalent to maximising the coefficient of determination $R^2 = 1-\langle\epsilon_i^2\rangle/\sigma^2_{\hat{c}(\mathbf{x})}$). The $R^2$ value for a set of (omitted) cross-validation data $\mathcal{C}_\text{xval}$ was used as the model's score.

Each regression model $f_c$ was used with $\mathcal{C}_\text{train}$ to build a low-dimensional TS model $\mathcal{T}_{(\phi_1, \phi_2, ..)}$ (cf.\ Fig.~\ref{figure:FIG-02-ts-schematic}(b)) in the corresponding CV space. This minimal TS description comprises the CV basis $(\phi_1, \phi_2, ..)$, the regression model $f_c$, and an estimate of the minimum free energy path (MEP) that connects the A and B basins. Whereas the $f_c$ isosurfaces are directly accessible by solving $f_c=0.5$, $f_c$ does not specify where the separatrix is most likely crossed (saddle point), nor does it specify the orientation of the MEP \cite{li_recent_2014}. In fact, the MEP may not be orthogonal to the committor isosurfaces in the chosen coordinate basis. Therefore, a partial MEP segment was estimated with a three point "string" $\bm{\phi}_{\text{MEP}} = [\bm{\phi}^{0.2}, \bm{\phi}^{0.5}, \bm{\phi}^{0.8}]$, where $\bm{\phi}^{0.2}$, $\bm{\phi}^{0.5}$, and $\bm{\phi}^{0.8}$ are averages of points in $\mathcal{C}_\text{train}$ binned by $\hat{c}(\mathbf{x})$ into the intervals $(0,0.4)$, $[0.4,0.6]$, and $(0.6,1.0)$, respectively. With this definition, $\bm{\phi}^{0.5}$ estimates the free energy saddle point. Because the points in $\mathcal{C}_\text{train}$ are drawn from $\tilde{V}(\mathbf{x},\mathbf{y})$, the MEP segment is only an estimate of the true MEP. We tested this estimate by comparing the committor derived $\mathcal{T}$ models with the computed PMFs for the unbiased system.

To test to what extent the H and I gizmos bias sampling to the same TS regions, we built two separate committor datasets $\mathcal{C}^{(H)}$ and $\mathcal{C}^{(I)}$ using H and I-gizmo biased simulations. Further details on the committor datasets are given in the Supporting Material.

%========================================================
\section{Methods}
%========================================================
%== MD boiler plate
MD simulations of the membrane-solvent-gizmo system were carried out essentially as if the gizmo were a transmembrane protein, using Gromacs 5.05 \cite{abraham_gromacs_2015}. The simulated system was a patch of 128 Berger DMPC lipids \cite{berger_molecular_1997} that was solvated with SPC water \cite{berendsen_interaction_1981} to a 43:1 water to lipid ratio using the MemGen web server \cite{knight_memgen_2015}. The gizmo was placed approximately at the membrane center. The simulation box used periodic boundary conditions and had dimensions 6.2 x 6.2 x 7.9 nm.

In all simulations, a temperature of 323 K was set with independent, velocity-rescale, thermostats \cite{bussi_canonical_2007} for solvent, lipids, and the gizmo, all using a coupling of 2.0 ps. A pressure of 1.0 bar, in NPT simulations, was maintained via  semi-isotropic weak coupling \cite{berendsen_molecular_1984} using a time constant of 1.0 ps. Center of mass motion was removed from the lipids, gizmo and solvent groups independently. The center atom of the gizmo was position restrained in the horizontal (x and y) dimensions with harmonic spring constants of 1000 kJ/(mol-nm$^2$). Electrostatics were computed with particle mesh Ewald \cite{essmann_smooth_1995} using a real space cutoff of 1.2 nm. A 1.2 nm cutoff was also used for van der Waals interactions.

Prior to production runs, the system was prepared using a steepest descent minimization followed by brief NVT and NPT runs (100 ps apiece) using a 2 fs timestep. Production runs used a 4 fs timestep. All NVT and NPT simulations used a stochastic integrator \cite{goga_efficient_2012}. Water bonds and angles were constrained by the SETTLE algorithm \cite{miyamoto_settle_1992}, and all other bonds were constrained using LINCS \cite{hess_lincs_1997}.

The Lennard-Jones interactions between the gizmo chain (CX beads) and lipid tails were scaled by using an alchemical mutation to turn CX beads into (non-interacting) GG beads, with $\lambda_{\text{CX}}$ implemented via the "vdw-lambdas" free energy option in Gromacs. The restraint potential for the gizmo belt radial breathing mode $V_\xi$ (see Eq.~\ref{eq_potential}) was implemented using the essential dynamics options in Gromacs. The Supporting Material includes example files for including a Gizmo in a Gromacs simulation.

%========================================================
\section{Results}
%========================================================
\subsection{Committor guided search for optimal RCs}

%==== FIGURE 3: committor regression models
\begin{figure*}[htbp] %\centering
\includegraphics[width=6.0in]{fig-03-regr-Hgizmo-newleg.eps}
\caption{\small{Committor regression models derived using $\mathcal{C}^{(H)}$, the H-gizmo committor dataset. (a)-(c) Top scoring 1D models for CVs using the OW (a), CxN (b) and NP (c) atom groups. Training and test datasets are drawn as purple and green dots, respectively, and a vertical gaussian jitter was added to the (discrete) $\hat{c}(\mathbf{x})$ values. The solid black lines show $f_c$ and have closed and open caps indicating the directions of the closed and open pore states, respectively. The dashed lines are $\pm$ two standard deviations from $f_c$, assuming 16 ($N_c$) samples drawn from a binomial distribution. (d) All regression models, 1D and 2D, co-sorted by $R^2$. The labeled orange points are the best 1D models for each atomgroup listed in Table \ref{TAB00}. (e) The regression fit for the 2D model using CVs $\phi_\text{CxN}$ and $\phi_\text{NP}$, with the same formatting as (a-c). (f) 2D view of the model in (e), showing the training dataset and the resulting TS model $\mathcal{T}_{(\phi_\text{CxN}, \phi_\text{NP})}^{(\text{H})}$. Individual points (small markers) are colored by $\hat{c}(\mathbf{x})$, with cyan, red, and blue dots belonging to the intervals (0, 0.4), [0.4, 0.6], and (0.6, 1.0) respectively. Grey squares and diamonds indicate $\hat{c}(\mathbf{x})$ = 0 or 1, respectively. Large markers are averages of the respective subsets and the ellipses are one standard deviation contours.}}
\label{figure:FIG-03-regression}
\end{figure*}

To test how accurately a single RC describes TS crossing, we first used the above sketched scheme to train and score 1D models (setting $\phi_2=0$) prior to building 2D models. These 1D models (derived from $\mathcal{C}^{(H)}$) yielded scores as high as $R^2=0.67$. The atom group was found to be highly predictive of $R^2$ (Fig.~\ref{figure:FIG-03-regression}), with only NP and P based CVs having $R^2>$ 0.38. The best CxN CVs had weaker predictive power ($R^2\approx0.35$) and the CVs involving water (PNO and OW) had $R^2 \leq 0.3$. In Fig.~\ref{figure:FIG-03-regression}, three examples of regression models are shown for the top scoring NP, CxN, and OW based CVs. 

This sorting suggests that TS passage is best characterized by localized headgroup (NP) merger. Lipid tail (CxN) expulsion is also involved, but to a much lesser extent, and the presence or absence of water (OW) in the nascent pore is, unexpectedly, not predictive of barrier crossing (at least by itself). The top ranked CV, with $R^2=0.67$, was $\text{rz}_{4}^{\text{NP}}$, which measures the axial merger of the four N or P atoms penetrating the bilayer. Presumably, this captures a pair of lipid headgroups merging from opposite sides. This CV is henceforth abbreviated as $\phi_\text{NP}$. We also abbreviate $\text{r}_{20}^{\text{CxN}}$ and $\text{rz}_{10}^{\text{OW}}$, the top ranked CxN and OW CVs, as $\phi_\text{CxN}$ and $\phi_\text{OW}$, respectively. 

According to this ranking, the PNO CVs, which group together the charged N and P atoms as well as water oxygens, are less predictive than NP and P alone. This result suggests that water and headgroup merger are not tightly coupled at the TS; it also suggests that the mechanism might proceed in a step-wise fashion.

Next, we considered 2D models, testing whether any linear combinations of two CVs were more predictive than $\phi_\text{NP}$ alone. As summarized in Fig.~\ref{figure:FIG-03-regression}, a few 2D models slightly outperformed $\phi_\text{NP}$. These improved models all contain $\phi_\text{NP}$ paired with another CV, however the increase in $R^2$ (at most $R^2=0.69$ as opposed to $R^2=0.67$) was modest.

The high ranking of NP and P regression models is consistent between $\mathcal{C}^{(H)}$ and $\mathcal{C}^{(I)}$ datasets, and the absolute $R^2$ values differ slightly (see Fig.~S5 for a side-by-side comparison). For the other atom groups with lower $R^2$ scores (OW, CxN, Cxx and PNO), there are differences in the relative rankings between the two datasets which we attribute to the gizmos facilitating similar, but not identical, pathways. These pathways were probed further by computing free energy landscapes.

%========================================================
\subsection{Free Energy Landscapes}
%========================================================
Next, we constructed free energy surfaces by projecting the PMF ensembles onto the optimized CVs constructed in the previous section. First, to examine headgroup merger and tail depletion, we used CVs  $\phi_\text{NP}$ and $\phi_\text{CxN}$. Later, to compare headgroup and water penetration we use $\phi_\text{NP}$ and the analogous water penetration coordinate. (PMF projection and unbiasing required a modified version of the weighted histogram analysis method (WHAM)\cite{kumar_weighted_1992}, described fully in the Supporting Material). We also tested how computed PMFs vary with gizmo design (H versus I) and tested for convergence using ensembles that were started from closed ('A') and open ('B') states.

Fig.~\ref{figure:FIG-04-pmf-NP-CxN} shows the free energy landscapes as a function of the two CVs $\phi_\text{NP}$ and $\phi_\text{CxN}$. In both the 2D and marginal PMFs, two clear minima can be seen, corresponding to the unperturbed membrane and the metastable prepore at a relative energy of +10 $k_{\rm B}T$. The barrier heights are between 14 and 17 $k_{\rm B}T$, which is consistent with the range of previously reported values for the same system \cite{bennett_atomistic_2014, awasthi_simulations_2016, grafmuller_free_2014}. The PMFs also resolve the full metastable prepore that has been experimentally predicted for decades \cite{abidor_electric_1979, evans_dynamic_2003}.

%== line tension estimate (this was hand drawn/eyeballed)
%After crossing the barrier, if $\phi_\text{CxN}$ is taken to measure radius for the expanding prepore, then a linear fit of $G(\phi_\text{CxN} > 1.5 nm)$ gives a line tension of $\approx 11$ pN, in line with previous estimates [].

%==================
%==== FIGURE 4: PMFs NP vs CxN
\begin{figure*}[htbp] %\centering
\includegraphics[width=6.2in]{fig-04-pmfs-NP-CxN.eps}
\caption{\small{2D (a-b) and 1D (c-d) PMFs of pore nucleation. (a),(b) PMFs computed using the I-gizmo (a) and H-gizmo (b), with ensembles seeded from the closed (A) state. Contours have 1 $k_{\rm B}T$ spacing. The insets show snapshots of pre-barrier structures, with the orange/yellow stars indicating their positions in ($\phi_\text{CxN}$, $\phi_\text{NP}$) space. In the snapshots, water is drawn as a stick with oxygen and hydrogen atoms in red and white, respectively. Lipid headgroup N and P atoms are drawn as blue and orange spheres, respectively, and the lipid tail carbons are illustrated as a grey slab that has been cut away. (c-d) Marginal 1D PMFs. For each RC, 1D PMFs are shown for four cases: both gizmo types (I and H) and both starting structures (A and B). In all panels, the $\mathcal{T}$ overlays are as described in Fig.~\ref{figure:FIG-02-ts-schematic}. Severely undersampled bins (bin count $\le$ 20), which make the PMF perimeters jagged, were not plotted, however this has no effect on the energetics or barrier heights shown.}}
\label{figure:FIG-04-pmf-NP-CxN}
\end{figure*}

In all cases considered in Fig.~\ref{figure:FIG-04-pmf-NP-CxN}, the quantitative agreement between A/B pairs suggests that the PMFs are sufficiently converged and do not suffer from marked hysteresis effects. An autocorrelation analysis (detailed in the Supporting Material) showed that intermediate windows in each ensemble exhibited slow, two-state switching corresponded to TS crossing, with autocorrelation times of 50-100ns. As this switching was clearly the sampling bottleneck, we extended these windows to 2$\mu s$ to ensure convergence. The resulting PMFs all show a net free energy change $\Delta G_{AB}$ of 9-11 $k_{\rm B}T$ irrespective of the particular choice of gizmo, CV, and starting structure. Only the barrier heights vary, which is to be expected.

The PMFs in Fig.~\ref{figure:FIG-04-pmf-NP-CxN} exhibit some clear differences that shed light on the nucleation mechanism. The location of energy barriers, and to some extent their heights, depend on the CV choice and the gizmo design. The I-gizmo simulations give barriers that are lower and later than the H-gizmo simulations. Snapshots shown in Fig.~\ref{figure:FIG-04-pmf-NP-CxN} ((a-b) insets), reveal the difference in paths. In (a) the membrane has partially thinned (I-gizmo), but in (b) a hydrophobic column already connects the opposite sides (H-gizmo). This result suggests that such hydrophobic defects are energetically unfavorable and unlikely to initiate nucleation, whereas an indentation pathway is energetically preferred by 3-4 $k_{\rm B}T$. 

Fig.~\ref{figure:FIG-04-pmf-NP-CxN} also shows that the computed energy barriers somewhat depend on the chosen CV, which underscores the well known fact that proper choice of RC is crucial. The marginal PMFs, shown in Fig.~\ref{figure:FIG-04-pmf-NP-CxN} (c-d), show that $\Delta G^{\ddagger}(\phi_\text{NP})$ is always higher than $\Delta G^{\ddagger}(\phi_\text{CxN})$, suggesting that $\phi_\text{NP}$ is better at distinguishing the A and B basins, and that state densities partially overlap when projected onto $\phi_\text{CxN}$. This projection overlap for $\phi_\text{CxN}$ is evident in the 2D PMFs (Fig.~\ref{figure:FIG-04-pmf-NP-CxN} (a-b)) as well. Given that $\phi_\text{CxN}$ primarily captures in-plane, radial depletion of lipids, this corroborates that pore radius is a suboptimal RC for pore nucleation, but is better suited to track pore expansion, such as would occur with applied surface tension.

Based on these findings, we take the PMF $G(\phi_\text{NP})$, for the I-gizmo, to be the most accurate (single RC) estimate of the true PMF. Here, $\phi_\text{NP}$ resolves the TS at values between 0.6 and 0.75, which is well before the headgroups fully reach the pore center, and before the RC saturates, at $\phi_\text{NP} \approx$ 0.2. The PMF $G(\phi_\text{NP})$ indicates a barrier for prepore closure of around 5 $k_{\rm B}T$. This value, to our knowledge, has not been reported previously because the full PMF, including the entire metastable basin, was not resolved with a single RC. However, a previous study reported that for twenty independent, unbiased simulations of DMPC pores, zero closed within 500ns \cite{awasthi_simulations_2016}, which suggests that the closure barrier is substantial.

Our previous committor analysis provides an independent estimate of the transition state and an additional control to determine which PMF results are the most accurate. Accordingly, we have overlaid the corresponding TS models, onto the PMFs in Fig.~\ref{figure:FIG-04-pmf-NP-CxN}. For the I-gizmo ensemble (panel (a) and marginals (c-d)), the TS model features align well with the projected PMF, and the $f_c(\mathbf{x})=0.5$ isosurface (black dashed line) cuts the PMF nearly at its saddle point. This effect is also seen in the marginals (panels (c-d)), where $\bm{\phi}^{0.5}$ (red dots) are near the PMF maxima (blue and green curves). In contrast, the PMF barriers are higher and much earlier for the H-gizmo ensembles and do not align well with $\bm{\phi}^{0.5}$. This result suggests that forming a hydrophobic column is energetically too costly, and thus provides additional evidence for the indentation path. 

%========================================================
\subsection{The role of water}
%========================================================
%== interplay of water and headgroups: the sequence of water and headgroup merger
Despite our finding that water CVs correlate poorly with the committor, one would expect water to partially hydrate the lipid headgroups that submerge to create the prepore. Thus, to examine this idea and to better resolve the sequence of events and the role of water, we computed a 2D PMF using a water CV ($\text{rz}_{4}^{\text{OW}}$) paired with $\phi_\text{NP}$. The CV $\text{rz}_4^{\small{\text{OW}}}$ was chosen because it measures the axial merger of the first (four) water molecules analogous to how $\phi_\text{NP}$ ($\text{rz}_{4}^{\text{NP}}$) measures the merger of headgroups.

%==== FIGURE 5 PMFs NP vs OW
\begin{figure*}[htbp] %\centering
\includegraphics[width=6.0in]{fig-05-pmfs-NP-OW.eps}
\caption{\small{ Free energy landscapes for headgroup and water CVs, $\phi_\text{NP}$ and $\text{rz}_{4}^{\text{OW}}$. The formatting is as in Fig.~\ref{figure:FIG-04-pmf-NP-CxN}. The $\mathcal{T}$ overlays in (c) and (d) are from the H-gizmo committor ensemble} }
\label{figure:FIG-05}
\end{figure*}

Fig.~\ref{figure:FIG-05} shows that, as was the case for $\phi_\text{CxN}$, $\text{rz}_{4}^{\text{OW}}$ does not cleanly distinguish the A and B basins and gives rise to projection overlaps, thus rendering it a poor RC by itself. The marginal PMFs in panel (d) do not reveal this overlap directly, but the PMF maximum and $\bm{\phi}^{0.5}$ (red dot in the $\mathcal{T}$ overlay) are in poor agreement. Indeed, in Fig.~\ref{figure:FIG-05} (a-b), the MEPs (cyan, red, and blue dots) are nearly orthogonal to $\text{rz}_{4}^{\text{OW}}$, suggesting that the water penetration is largely orthogonal to TS crossing. In addition, $\text{rz}_{4}^{\text{OW}}$ is a poor committor correlate ($R^2=0.32$ and $R^2=0.07$ for $\mathcal{C}^{(I)}$ and $\mathcal{C}^{(H)}$ datasets, respectively).

In contrast to the linear MEPs seen in Fig.~\ref{figure:FIG-04-pmf-NP-CxN}, both of the 2D PMFs in Fig.~\ref{figure:FIG-05} (a),(b) exhibit a kinked pathway in the TS region. The kink results from $\text{rz}_{4}^{\text{OW}}$ saturating at its lowest values prior to the TS. The MEP overlay also shows this kink at the cyan dot, $\bm{\phi}^{0.2}$. 

Taken together, these findings sugget a "water first" mechanism, where a small number of waters reach the pore center prior to the headgroups and prior to $\bm{\phi}^{0.5}$. The subsequent barrier crossing, $\bm{\phi}^{0.2} \rightarrow \bm{\phi}^{0.5} \rightarrow \bm{\phi}^{0.8}$, follows a straight line in the projected space, primarily along the $\phi_\text{NP}$ coordinate. 

%== ts model snapshots: best agreement and disagreement
To further examine the sequential steps of nucleation and the roles of hydration and headgroups we used the trained TS model for the I-gizmo, $\mathcal{T}^{(I)}_{(\phi_\text{NP}, \text{rz}_{4}^{\text{OW}})}$, to select representative structures at different stages of nucleation. Fig.~\ref{figure:FIG-06} shows those frames (points $\mathbf{x}$ from $\mathcal{C}^{(I)}$) from just before, at, and just after the TS (rows (c),(e),(g), respectively) which agree best with $f_c$ (residuals $|\epsilon_i| \leq0.1$). The pre-barrier frames (row (c)) all yield $\hat{c}(\mathbf{x})<0.2$ and illustrate the diversity of water defects that pierce the membrane prior to the TS. This finding is consistent with the PMFs and kinked MEP described above, indicating that hydration defects precede the TS. Next, as the barrier is crossed, the merger of lipid headgroups ($\phi_\text{NP}$ decreasing) is also clearly visible by comparing rows (c),(e), and (g). Within these rows, there are variations in hydration, shearing/skewing, pyramidal indentations, and axial asymmetry, suggesting that this variability is orthogonal to the MEP and thus does not correlate with the progress of the reaction. The snapshots show that the submerged headgroups are hydrated, as expected; however, the size and shape of the water clusters appear too variable for water to serve as a precise measure of reaction progress \cite{fathizadeh_ion_2019}.

%==== FIGURE 6: TS snapshots Igizmo
\begin{figure*}[htbp] %\centering
\includegraphics[width=6.25in]{fig-06-tsm-snapshots-Igizmo.png}
\caption{\small{Configurations showing TS crossing, taken from the $\mathcal{C}^{(I)}$ dataset. The regression model (a) and the corresponding TS model (b) for the CVs $\phi_\text{NP}$ and $\text{rz}_{4}^{\text{OW}}$. In (a), $f_c$ is shown as a solid line and the dotted lines are $\pm$ two standard deviations, assuming 16 ($N_c$) samples drawn from a binomial distribution. In (b) the solid and dashed lines correspond to those in Fig.~\ref{figure:FIG-05}(a). In (a) and (b), the committor dataset is shown as grey dots. The colored dots indicate configurations that are illustrated in rows (c)-(g). These illustrated configurations were chosen based on $\hat{c}(\mathbf{x})$ values and residual errors. Rows (c), (e), and (g) show configurations from before, at, and after the TS respectively, with small residual errors. Configurations in rows (d) and (f) have large negative and positive error residuals, respectively.}}
\label{figure:FIG-06}
\end{figure*}

For comparison, we also plotted snapshots from cases where $\hat{c}(\mathbf{x})$ is severely under- or over-predicted in rows (d) and (f), respectively. The structures in rows (d)-(f) are visually similar and no striking structural differences stand out to suggest other predictive CVs.

A considerable amount of the $\hat{c}(\mathbf{x})$ variance here stems from using imprecise ($N_c=16$) committor estimates. Whereas upwards of 100 shots are needed to converge a single $\hat{c}(\mathbf{x})$ estimate, this is largely due to statistical uncertainty. Indeed, binomial deconvolution methods can be used to estimate committor histograms rigorously, at 10x reduced cost\cite{peters_using_2006}. Here, as Figs.~\ref{figure:FIG-06}(a) and \ref{figure:FIG-03-regression}(c,e) show, the observed variance in $\hat{c}(\mathbf{x}|N_c=16)$ is indeed comparable to the statistical uncertainty (dotted lines above and below the regression curve $f_c$). This suggests that, for the purpose of ranking CVs, only low resolution committor estimates are required, provided a sufficient number of uncorrelated configurations near the TS. (For more information about the comittor datasets see the Supporting Material.)

%=================================================
\section{Discussion}
%=================================================
We have used a combination of atomistic MD simlations, free energy calculations, and committor analysis to probe the mechanism of metastable prepore formation in a lipid membrane.

% NARRATIVE: patchwork of studies looking at this and that (trying for only a condensed enumeration of examples here)
Several previous simulation studies of poration have proposed and employed one or a few possible RCs, considering specific atom groups and CVs in isolation. These approaches include restraining a single lipid headgroup relative to the bilayer midplane \cite{bennett_water_2011, awasthi_simulations_2016, tieleman_lipids_2006, grafmuller_free_2014, nishizawa_molecular_2013}, biasing hydrophilic atoms to occupy a stack of cylindrical slabs spanning the membrane \cite{hub_probing_2017}, biasing water to occupy a cylindrical column \cite{mirjalili_density-biased_2014}, and growing a lateral depletion of lipid centers of mass \cite{wohlert_free_2006}.

% NARRATIVE: our systematic study
Here we adopted a more general approach and constructed a combinatoric pool of CVs to systematically vary the different factors relevant to the mechanism, including the atom group, geometric bias, and locality (via $N$, the number of atoms in the CV definition). These CVs were then assessed systematically as RCs, using a committor regression scheme. For comparison, hybrid RCs built from pairwise linear combinations of individual CVs were also tested, but not found to be substantially superior.

% NARRATIVE: RC quality largely depends on atom group choice
Our results showed that the achievable RC quality largely depends on the choice of atom group. Headgroup (NP and P) based CVs yielded the highest correlation with the committor and in this sense represented the best descriptors for the RC in the vicinity of the TS. In contrast, and somewhat unexpectedly, CVs based only on the position of water molecules (the OW family) provided remarkably poor RCs. Also, the PNO CVs (grouping N,P, and (water) O atoms)  described the RC less accurately than NP alone, suggesting that these hydrophilic components have distinct mechanistic roles in the TS crossing. Lipid tail based CVs (CxN) were also less predictive than headgroups.

% NARRATIVE: atom groups and geometry
Among the headgroup RCs (and among all RCs tested), $\text{rz}_{4}^{\text{NP}}$ ($\phi_\text{NP}$), which essentially measures the gap between two penetrating lipid headgroups, provided the best description of the commitor RC. Interestingly, none of the combinations of $\phi_\text{NP}$ and CVs involving larger numbers (10--20) of atoms, which measure larger collective motion, substantially outperformed $\phi_\text{NP}$ alone. This result suggests that barrier crossing is more localized than one might have expected.

% a few specific comparisons..
This particular RC $\phi_\text{NP}$ somewhat resembles the hydrophobic belt height suggested by Akimov et al. \cite{akimov_pore_2017}. The fact that our systematic and unbiased search highlighted this RC, corroborates its mechanistic relevance and utility in continuum modeling. Our RC $\phi_\text{NP}$ is also similar to the slab occupancy metric \cite{hub_probing_2017} referred to above. Both RCs essentially measure the gap between the penetrating hydrophilic material, but for different groups of atoms, which turned out to be an important distinction. This RC, as well as the other RCs considered here, also provide a rather smoother measure of merger progress, which does not require spatial discretization and thereby avoids Poisson noise due to near-zero bin counts.

Overall, our committor analysis and RC ranking underscores the importance of headgroup merger, both as a mechanistic descriptor and biasing coordinate. However, regarding the precise molecular mechanism, it is crucial to systematically define and asses RCs against the, by definition, optimal committor RC. Unexpectedly, water turned out to be rather weakly correlated with the progress of pore formation; further, the degree of hydration fluctuates markedly during poration, which, taken together, renders hydration a rather poor RC. Water based CVs (by themselves), therefore, seem to provide less suitable biasing coordinates for portion and, possibly, similar systems and processes such as membrane fusion. Lateral depletion coordinates, in contrast, such as $\phi_\text{CxN}$, are better for describing pore expansion than nucleation.

To obtain sufficient sampling of the hard-to-reach TS region, we developed a new biasing scheme, the pore gizmo. The intended effect of this gizmo bias potential, exerted via repulsive beads, is similar in spirit to other schemes using repulsive plates \cite{kawamoto_free_2014}, hydrophilic beads \cite{risselada_expansion_2014}, depletion coordinates \cite{wohlert_free_2006}, or constrictive "cuff potentials" \cite{fuhrmans_coarse-grained_2015}. What distinguishes the gizmo, and, as it turned out, renders it particularly suitable for the present purpose, is its flexibility, which provides the combined system with sufficient flexibility to enhance sampling and thereby to also probe asymmetric states. In particular, the axial flexibility of the CX chain allows for proper sampling of asymmetric or skewed tail packing defects.

% mechanism as a whole
Using our gizmo biasing potential enabled us to obtain sufficient sampling to study, by atomistic simulations, the full poration pathway and the mechanistic steps of a DMPC lipid bilayer prior to nucleation. Two different biasing schemes were used to direct a membrane thinning (indentation) path and a hydrophobic defect path, both leading to open pores. By computing PMFs, projected onto the optimal RCs determined before, and by comparing snapshots close to the TS, a sequential penetration of water and headgroups was resolved.

Taken together, our simulations and committor-based RCs suggest the following mechanism. A pore nucleates via an elastic indentation that is energetically preferred to forming a hydrophobic defect. After thinning, but just prior to the TS, water pierces the thinned slab. This water defect is required but does not suffice to finally nucleate the pore. Instead, it is the axial merger of the first lipid headgroups from opposite monolayers ($\phi_\text{NP}$) that precedes --- and best characterizes --- the subsequent nucleation and thus crossing of the TS.

This mechanism could be biologically relevant for the structure and rupture of the hemifusion diaphragm (HD) \cite{kozlovsky_lipid_2002, nikolaus_direct_2010}. In this context, the HD perimeter is a three-way membrane junction that is predicted to stabilize, and potentially lower the nucleation barrier for, transient pore defects \cite{risselada_free_2014}. These rim pores are likely on-path precursors to rupture, and therefore might also explain flickering prior to HD opening \cite{chanturiya_flickering_1997}. A similar combination of enhanced sampling and committor analysis could enable one to study this and other biologically relevant topological membrane remodelling mechanisms, such as fusion stalk formation. 

Similar to lipids, intrinsically disordered proteins (IDPs) can regulate transport, but are challenging to control and characterize structurally. For example, there is an ongoing discussion on how the IDPs that form the interior of nuclear pore complexes (Phe–Gly nucleoporins or FG-Nups) are organized and spatially distributed within the nuclear pore in order to achieve selectivity for karyopherins \cite{ketterer_dna_2018}. Gizmo based enhanced sampling may enable one to energetically distinguish between these hypotheses.

In a broader context, future studies that seek to uncover atomistic mechanisms and to compute energetics with $k_BT$ precision particularly for permutationally frustrated systems such as membranes, solvent surface layers, or IDPs, will benefit from hybrid schemes that combine committor analysis based RC optimisation and free energy calculations using gizmo biased enhanced sampling.

\section{Acknowledgements}
The authors gratefully acknowledge helpful discussions with Martin Mechelke, Maxim Igaev, Vytautas Gapsys, Jochen Hub, Neha Awasthi, and H. Jelger Risselada and financial support from the DFG Grant No. SFB 803.B2.

\section{Author Contributions}
GB and HG designed research. GB performed research and analyzed data. GB and HG wrote the manuscript.

\section{Supporting References}
References \cite{chodera_use_2007, tan_theory_2012, oostenbrink_biomolecular_2004, shirts_statistically_2008} appear in the Supporting Material.

\section{References}

\bibliography{refs}

%\printbibliography

\end{document}
