## Supplementary material for "Sequential water and headgroup merger: Membrane poration paths and energetics from MD simulations": pore-gizmo.zip: refs.bib

@article{pronk_gromacs_2013,
	title = {{GROMACS} 4.5: a high-throughput and highly parallel open source molecular simulation toolkit},
	volume = {29},
	issn = {1367-4803, 1460-2059},
	shorttitle = {{GROMACS} 4.5},
	url = {http://bioinformatics.oxfordjournals.org/content/29/7/845},
	doi = {10.1093/bioinformatics/btt055},
	abstract = {Motivation: Molecular simulation has historically been a low-throughput technique, but faster computers and increasing amounts of genomic and structural data are changing this by enabling large-scale automated simulation of, for instance, many conformers or mutants of biomolecules with or without a range of ligands. At the same time, advances in performance and scaling now make it possible to model complex biomolecular interaction and function in a manner directly testable by experiment. These applications share a need for fast and efficient software that can be deployed on massive scale in clusters, web servers, distributed computing or cloud resources.
Results: Here, we present a range of new simulation algorithms and features developed during the past 4 years, leading up to the GROMACS 4.5 software package. The software now automatically handles wide classes of biomolecules, such as proteins, nucleic acids and lipids, and comes with all commonly used force fields for these molecules built-in. GROMACS supports several implicit solvent models, as well as new free-energy algorithms, and the software now uses multithreading for efficient parallelization even on low-end systems, including windows-based workstations. Together with hand-tuned assembly kernels and state-of-the-art parallelization, this provides extremely high performance and cost efficiency for high-throughput as well as massively parallel simulations.
Availability: GROMACS is an open source and free software available from http://www.gromacs.org.
Contact: erik.lindahl\{at\}scilifelab.se
Supplementary information: Supplementary data are available at Bioinformatics online.},
	language = {en},
	number = {7},
	urldate = {2014-01-15},
	journal = {Bioinformatics},
	author = {Pronk, Sander and Páll, Szilárd and Schulz, Roland and Larsson, Per and Bjelkmar, Pär and Apostolov, Rossen and Shirts, Michael R. and Smith, Jeremy C. and Kasson, Peter M. and Spoel, David van der and Hess, Berk and Lindahl, Erik},
	month = apr,
	year = {2013},
	pages = {845--854},
	file = {Full Text PDF:/home/gbubnis/Zotero/storage/8HCADZGS/Pronk et al. - 2013 - GROMACS 4.5 a high-throughput and highly parallel.pdf:application/pdf;Snapshot:/home/gbubnis/Zotero/storage/8R9GVFFB/845.html:text/html}
}

@article{risselada_expansion_2014,
	title = {Expansion of the fusion stalk and its implication for biological membrane fusion},
	volume = {111},
	issn = {0027-8424, 1091-6490},
	url = {http://www.pnas.org/content/111/30/11043},
	doi = {10.1073/pnas.1323221111},
	abstract = {Over the past 20 years, it has been widely accepted that membrane fusion proceeds via a hemifusion step before opening of the productive fusion pore. An initial hourglass-shaped lipid structure, the fusion stalk, is formed between the adjacent membrane leaflets (cis leaflets). It remains controversial if and how fusion proteins drive the subsequent transition (expansion) of the stalk into a fusion pore. Here, we propose a comprehensive and consistent thermodynamic understanding in terms of the underlying free-energy landscape of stalk expansion. We illustrate how the underlying free energy landscape of stalk expansion and the concomitant pathway is altered by subtle differences in membrane environment, such as leaflet composition, asymmetry, and flexibility. Nonleaky stalk expansion (stalk widening) requires the formation of a critical trans-leaflet contact. The fusion machinery can mechanically enforce trans-leaflet contact formation either by directly enforcing the trans-leaflets in close proximity, or by (electrostatically) condensing the area of the cis leaflets. The rate of these fast fusion reactions may not be primarily limited by the energetics but by the forces that the fusion proteins are able to exert.},
	language = {en},
	number = {30},
	urldate = {2015-02-23},
	journal = {Proceedings of the National Academy of Sciences},
	author = {Risselada, Herre Jelger and Bubnis, Gregory and Grubmuller, Helmut},
	month = jul,
	year = {2014},
	keywords = {diaphragm, hemagglutinin, hemi, neuronal, SNARE},
	pages = {11043--11048},
	file = {Full Text PDF:/home/gbubnis/Zotero/storage/Q7WWJVGW/Risselada et al. - 2014 - Expansion of the fusion stalk and its implication .pdf:application/pdf;Snapshot:/home/gbubnis/Zotero/storage/V3AN3M5A/11043.html:text/html}
}

@article{bennett_water_2011,
	title = {Water {Defect} and {Pore} {Formation} in {Atomistic} and {Coarse}-{Grained} {Lipid} {Membranes}: {Pushing} the {Limits} of {Coarse} {Graining}},
	volume = {7},
	issn = {1549-9618},
	shorttitle = {Water {Defect} and {Pore} {Formation} in {Atomistic} and {Coarse}-{Grained} {Lipid} {Membranes}},
	url = {http://dx.doi.org/10.1021/ct200291v},
	doi = {10.1021/ct200291v},
	abstract = {Defects in lipid bilayers are important in a range of biological processes, including interactions between antimicrobial peptides and membranes, transport of polar molecules (including drugs) across membranes, and lipid flip?flop from one monolayer to the other. Passive lipid flip?flop and the translocation of polar molecules across lipid membranes occur on a slow time scale because of high-energy intermediates involving water defects and pores in the membrane. Such defects are an interesting test case for coarse-grained models because of their relatively small characteristic size at the level of water molecules and the complex environment of water and polar head groups in a low-dielectric membrane interior. Here we compare coarse-grained simulations with the MARTINI model with the standard MARTINI water and two recently developed coarse-grained polarizable water models to atomistic simulations. Although in several cases the MARTINI model reproduces the correct free energies, there are structural differences between the atomistic and coarse-grained models. The polarizable water model improves the free energies but only moderately improves the structures. Atomistic test simulations in which water molecules are artificially tethered to each other in groups of four, the resolution of MARTINI, suggest that the limiting factor is not the size of the coarse-grained particles but rather the simple interaction potential and/or the entropy lost in coarse graining the system. By increasing the attractive interaction between the lipids? headgroup and water, we did observe pore formation but at the expense of the correct equilibrium properties of the bilayers.},
	number = {9},
	urldate = {2015-01-22},
	journal = {Journal of Chemical Theory and Computation},
	author = {Bennett, W.F. Drew and Tieleman, D. Peter},
	year = {2011},
	pages = {2981--2988},
	file = {ACS Full Text PDF w/ Links:/home/gbubnis/Zotero/storage/8NVD2UVW/Bennett and Tieleman - 2011 - Water Defect and Pore Formation in Atomistic and C.pdf:application/pdf;ACS Full Text Snapshot:/home/gbubnis/Zotero/storage/MMKHEKDS/ct200291v.html:text/html}
}

@article{mirjalili_density-biased_2014,
	title = {Density-{Biased} {Sampling}: {A} {Robust} {Computational} {Method} for {Studying} {Pore} {Formation} in {Membranes}},
	volume = {11},
	issn = {1549-9618},
	shorttitle = {Density-{Biased} {Sampling}},
	url = {http://dx.doi.org/10.1021/ct5009153},
	doi = {10.1021/ct5009153},
	abstract = {A new reaction coordinate to bias molecular dynamics simulation is described that allows enhanced sampling of density-driven processes, such as mixing and demixing two different molecular species. The methodology is validated by comparing the theoretical entropy of demixing two ideal gas species and then applied to induce deformation and pore formation in phospholipid membranes within an umbrella sampling framework. Comparison with previous biased simulations of membrane pore formation suggests overall quantitative agreement, but the density-based biasing potential results in a different, more realistic transition pathway than that in previous studies.},
	number = {1},
	urldate = {2015-01-22},
	journal = {Journal of Chemical Theory and Computation},
	author = {Mirjalili, Vahid and Feig, Michael},
	year = {2014},
	pages = {343--350},
	file = {ACS Full Text PDF w/ Links:/home/gbubnis/Zotero/storage/KVEBSPEJ/Mirjalili and Feig - 2014 - Density-Biased Sampling A Robust Computational Me.pdf:application/pdf;ACS Full Text Snapshot:/home/gbubnis/Zotero/storage/DNGV7AMS/ct5009153.html:text/html}
}

@article{tiwary_spectral_2016,
	title = {Spectral gap optimization of order parameters for sampling complex molecular systems},
	volume = {113},
	copyright = {©  . http://www.pnas.org/preview\_site/misc/userlicense.xhtml},
	issn = {0027-8424, 1091-6490},
	url = {https://www.pnas.org/content/113/11/2839},
	doi = {10.1073/pnas.1600917113},
	abstract = {In modern-day simulations of many-body systems, much of the computational complexity is shifted to the identification of slowly changing molecular order parameters called collective variables (CVs) or reaction coordinates. A vast array of enhanced-sampling methods are based on the identification and biasing of these low-dimensional order parameters, whose fluctuations are important in driving rare events of interest. Here, we describe a new algorithm for finding optimal low-dimensional CVs for use in enhanced-sampling biasing methods like umbrella sampling, metadynamics, and related methods, when limited prior static and dynamic information is known about the system, and a much larger set of candidate CVs is specified. The algorithm involves estimating the best combination of these candidate CVs, as quantified by a maximum path entropy estimate of the spectral gap for dynamics viewed as a function of that CV. The algorithm is called spectral gap optimization of order parameters (SGOOP). Through multiple practical examples, we show how this postprocessing procedure can lead to optimization of CV and several orders of magnitude improvement in the convergence of the free energy calculated through metadynamics, essentially giving the ability to extract useful information even from unsuccessful metadynamics runs.},
	language = {en},
	number = {11},
	urldate = {2019-05-01},
	journal = {Proceedings of the National Academy of Sciences},
	author = {Tiwary, Pratyush and Berne, B. J.},
	month = mar,
	year = {2016},
	keywords = {caliber, collective variables, enhanced sampling, spectral gap, timescale separation},
	pages = {2839--2844},
	file = {Full Text PDF:/home/gbubnis/Zotero/storage/E6Y9NPXZ/Tiwary and Berne - 2016 - Spectral gap optimization of order parameters for .pdf:application/pdf;Snapshot:/home/gbubnis/Zotero/storage/MY4TB4DA/2839.html:text/html}
}

@article{bussi_canonical_2007,
	title = {Canonical sampling through velocity rescaling},
	volume = {126},
	issn = {0021-9606, 1089-7690},
	url = {http://scitation.aip.org/content/aip/journal/jcp/126/1/10.1063/1.2408420},
	doi = {10.1063/1.2408420},
	abstract = {The authors present a new molecular dynamics algorithm for sampling the canonical distribution. In this approach the velocities of all the particles are rescaled by a properly chosen random factor. The algorithm is formally justified and it is shown that, in spite of its stochastic nature, a quantity can still be defined that remains constant during the evolution. In numerical applications this quantity can be used to measure the accuracy of the sampling. The authors illustrate the properties of this new method on Lennard-Jones and TIP4P water models in the solid and liquid phases. Its performance is excellent and largely independent of the thermostat parameter also with regard to the dynamic properties.},
	number = {1},
	urldate = {2014-04-16},
	journal = {The Journal of Chemical Physics},
	author = {Bussi, Giovanni and Donadio, Davide and Parrinello, Michele},
	month = jan,
	year = {2007},
	keywords = {Molecular dynamics, Classical ensemble theory, Equations of motion, Ice, Water energy interactions},
	pages = {014101},
	file = {Full Text PDF:/home/gbubnis/Zotero/storage/824JWUS3/Bussi et al. - 2007 - Canonical sampling through velocity rescaling.pdf:application/pdf;Snapshot:/home/gbubnis/Zotero/storage/KDEAQ3EI/1.html:text/html}
}

@article{chernomordik_hemifusion_1995,
	title = {The hemifusion intermediate and its conversion to complete fusion: regulation by membrane composition},
	volume = {69},
	issn = {0006-3495},
	shorttitle = {The hemifusion intermediate and its conversion to complete fusion},
	url = {http://www.sciencedirect.com/science/article/pii/S0006349595799660},
	doi = {10.1016/S0006-3495(95)79966-0},
	abstract = {To fuse, membranes must bend. The energy of each lipid monolayer with respect to bending is minimized at the spontaneous curvature of the monolayer. Two lipids known to promote opposite spontaneous curvatures, lysophosphatidylcholine and arachidonic acid, were added to different sides of planar phospholipid membranes. Lysophosphatidylcholine added to the contacting monolayers of fusing membranes inhibited the hemifusion we observed between lipid vesicles and planar membranes. In contrast, fusion pore formation depended upon the distal monolayer of the planar membrane; lysophosphatidylcholine promoted and arachidonic acid inhibited. Thus, the intermediates of hemifusion and fusion pores in phospholipid membranes involve different membrane monolayers and may have opposite net curvatures, Biological fusion may proceed through similar intermediates.},
	number = {3},
	urldate = {2016-04-11},
	journal = {Biophysical Journal},
	author = {Chernomordik, L. and Chanturiya, A. and Green, J. and Zimmerberg, J.},
	month = sep,
	year = {1995},
	pages = {922--929},
	file = {ScienceDirect Full Text PDF:/home/gbubnis/Zotero/storage/TJKQHJ49/Chernomordik et al. - 1995 - The hemifusion intermediate and its conversion to .pdf:application/pdf;ScienceDirect Snapshot:/home/gbubnis/Zotero/storage/FDHQ37MF/S0006349595799660.html:text/html}
}

@article{bennett_atomistic_2014,
	title = {Atomistic {Simulations} of {Pore} {Formation} and {Closure} in {Lipid} {Bilayers}},
	volume = {106},
	issn = {0006-3495},
	url = {http://www.cell.com/article/S0006349513057457/abstract},
	doi = {10.1016/j.bpj.2013.11.4486},
	abstract = {Cellular membranes separate distinct aqueous compartments, but can be breached by transient hydrophilic pores. A large energetic cost prevents pore formation, which is largely dependent on the composition and structure of the lipid bilayer. The softness of bilayers and the disordered structure of pores make their characterization difficult. We use molecular-dynamics simulations with atomistic detail to study the thermodynamics, kinetics, and mechanism of pore formation and closure in DLPC, DMPC, and DPPC bilayers, with pore formation free energies of 17, 45, and 78 kJ/mol, respectively. By using atomistic computer simulations, we are able to determine not only the free energy for pore formation, but also the enthalpy and entropy, which yields what is believed to be significant new insights in the molecular driving forces behind membrane defects. The free energy cost for pore formation is due to a large unfavorable entropic contribution and a favorable change in enthalpy. Changes in hydrogen bonding patterns occur, with increased lipid-water interactions, and fewer water-water hydrogen bonds, but the total number of overall hydrogen bonds is constant. Equilibrium pore formation is directly observed in the thin DLPC lipid bilayer. Multiple long timescale simulations of pore closure are used to predict pore lifetimes. Our results are important for biological applications, including the activity of antimicrobial peptides and a better understanding of membrane protein folding, and improve our understanding of the fundamental physicochemical nature of membranes.},
	language = {English},
	number = {1},
	urldate = {2015-01-22},
	journal = {Biophysical Journal},
	author = {Bennett, W. F. Drew and Sapay, Nicolas and Tieleman, D. Peter},
	month = jul,
	year = {2014},
	pages = {210--219},
	file = {Full Text PDF:/home/gbubnis/Zotero/storage/RRAQQJIZ/Bennett et al. - 2014 - Atomistic Simulations of Pore Formation and Closur.pdf:application/pdf;Snapshot:/home/gbubnis/Zotero/storage/CHN4VWCJ/S0006-3495(13)05745-7.html:text/html}
}

@article{berryman_free_2013,
	title = {Free {Energies} by {Thermodynamic} {Integration} {Relative} to an {Exact} {Solution}, {Used} to {Find} the {Handedness}-{Switching} {Salt} {Concentration} for {DNA}},
	volume = {9},
	issn = {1549-9618},
	url = {http://dx.doi.org/10.1021/ct3005968},
	doi = {10.1021/ct3005968},
	abstract = {Sets of free energy differences are useful for finding the equilibria of chemical reactions, while absolute free energies have little physical meaning. However finding the relative free energy between two macrostates by subtraction of their absolute free energies is a valuable strategy in certain important cases. We present calculations of absolute free energies of biomolecules, using a combination of the well-known Einstein molecule method (for treating the solute) with a conceptually related method of recent genesis for computing free energies of liquids (to treat the solvent and counterions). The approach is based on thermodynamic integration from a detailed atomistic model to one which is simplified but analytically solvable, thereby giving the absolute free energy as that of the tractable model plus a correction term found numerically. An example calculation giving the free energy with respect to salt concentration for the B- and Z-isomers of all-atom duplex DNA in explicit solvent and counterions is presented. The coexistence salt concentration is found with unprecedented accuracy.},
	number = {1},
	urldate = {2016-04-14},
	journal = {Journal of Chemical Theory and Computation},
	author = {Berryman, Joshua T. and Schilling, Tanja},
	month = jan,
	year = {2013},
	pages = {679--686},
	file = {ACS Full Text PDF w/ Links:/home/gbubnis/Zotero/storage/R5CJCCMK/Berryman and Schilling - 2013 - Free Energies by Thermodynamic Integration Relativ.pdf:application/pdf;ACS Full Text Snapshot:/home/gbubnis/Zotero/storage/UW4JIZBF/ct3005968.html:text/html}
}

@article{kumar_weighted_1992,
	title = {{THE} weighted histogram analysis method for free-energy calculations on biomolecules. {I}. {The} method},
	volume = {13},
	copyright = {Copyright © 1992 John Wiley \& Sons, Inc.},
	issn = {1096-987X},
	url = {http://onlinelibrary.wiley.com/doi/10.1002/jcc.540130812/abstract},
	doi = {10.1002/jcc.540130812},
	abstract = {The Weighted Histogram Analysis Method (WHAM), an extension of Ferrenberg and Swendsen's Multiple Histogram Technique, has been applied for the first time on complex biomolecular Hamiltonians. The method is presented here as an extension of the Umbrella Sampling method for free-energy and Potential of Mean Force calculations. This algorithm possesses the following advantages over methods that are currently employed: (1) It provides a built-in estimate of sampling errors thereby yielding objective estimates of the optimal location and length of additional simulations needed to achieve a desired level of precision; (2) it yields the “best” value of free energies by taking into account all the simulations so as to minimize the statistical errors; (3) in addition to optimizing the links between simulations, it also allows multiple overlaps of probability distributions for obtaining better estimates of the free-energy differences. By recasting the Ferrenberg–Swendsen Multiple Histogram equations in a form suitable for molecular mechanics type Hamiltonians, we have demonstrated the feasibility and robustness of this method by applying it to a test problem of the generation of the Potential of Mean Force profile of the pseudorotation phase angle of the sugar ring in deoxyadenosine. © 1992 by John Wiley \& Sons, Inc.},
	language = {en},
	number = {8},
	urldate = {2014-10-30},
	journal = {Journal of Computational Chemistry},
	author = {Kumar, Shankar and Rosenberg, John M. and Bouzida, Djamal and Swendsen, Robert H. and Kollman, Peter A.},
	month = oct,
	year = {1992},
	pages = {1011--1021},
	file = {Full Text PDF:/home/gbubnis/Zotero/storage/8ADWPVZT/Kumar et al. - 1992 - THE weighted histogram analysis method for free-en.pdf:application/pdf;Snapshot:/home/gbubnis/Zotero/storage/CESGTBM2/abstract.html:text/html;WHAMfulltext.pdf:/home/gbubnis/Zotero/storage/4W5NH82A/WHAMfulltext.pdf:application/pdf}
}

@article{torrie_nonphysical_1977,
	title = {Nonphysical sampling distributions in {Monte} {Carlo} free-energy estimation: {Umbrella} sampling},
	volume = {23},
	issn = {0021-9991},
	shorttitle = {Nonphysical sampling distributions in {Monte} {Carlo} free-energy estimation},
	url = {http://www.sciencedirect.com/science/article/B6WHY-4DDR2HH-3V/2/0884c987f3a8b098b688e68aa847371f},
	doi = {10.1016/0021-9991(77)90121-8},
	abstract = {The free energy difference between a model system and some reference system can easily be written as an ensemble average, but the conventional Monte Carlo methods of obtaining such averages are inadequate for the free-energy case. That is because the Boltzmann-weighted sampling distribution ordinarily used is extremely inefficient for the purpose. This paper describes the use of arbitrary sampling distributions chosen to facilitate such estimates. The methods have been tested successfully on the Lennard-Jones system over a wide range of temperature and density, including the gas-liquid coexistence region, and are found to be extremely powerful and economical.},
	number = {2},
	urldate = {2010-04-13},
	journal = {Journal of Computational Physics},
	author = {Torrie, G. M. and Valleau, J. P.},
	month = feb,
	year = {1977},
	pages = {187--199},
	file = {ScienceDirect Full Text PDF:/home/gbubnis/Zotero/storage/EF5M7P36/Torrie and Valleau - 1977 - Nonphysical sampling distributions in Monte Carlo .pdf:application/pdf;ScienceDirect Snapshot:/home/gbubnis/Zotero/storage/TVVXEH9V/0021999177901218.html:text/html}
}

@article{kawamoto_free_2014,
	title = {Free energy analysis along the stalk mechanism of membrane fusion},
	volume = {10},
	issn = {1744-6848},
	url = {http://pubs.rsc.org/en/content/articlelanding/2014/sm/c3sm52344f},
	doi = {10.1039/C3SM52344F},
	abstract = {The free energy profile of the stalk model of membrane fusion has been calculated using coarse-grained molecular dynamics simulations. The proposed method guides the lipid configuration using a guiding wall potential to make the transition from two apposed membranes to a stalk and a fusion pore. The free energy profile is obtained with a thermodynamic integration scheme using the mean force working on the guiding wall as a response of the system. We applied the method to two apposed flat bilayers composed of dioleoyl phosphatidylethanolamine/dioleoyl phosphatidylcholine expanding over the simulation box under the periodic boundary conditions. The two transition states are identified as pre-stalk and pre-pore states. The free energy barrier for the latter is confirmed to be in good agreement with that estimated by the pulling method. The present method provides a practical way to calculate the free energy profile along the stalk mechanism.},
	language = {en},
	number = {17},
	urldate = {2014-10-28},
	journal = {Soft Matter},
	author = {Kawamoto, Shuhei and Shinoda, Wataru},
	month = apr,
	year = {2014},
	pages = {3048--3054},
	file = {Full Text PDF:/home/gbubnis/Zotero/storage/8GBQNZ7E/Kawamoto and Shinoda - 2014 - Free energy analysis along the stalk mechanism of .pdf:application/pdf;Full Text PDF:/home/gbubnis/Zotero/storage/BFC3WIGC/Kawamoto and Shinoda - 2014 - Free energy analysis along the stalk mechanism of .pdf:application/pdf;Snapshot:/home/gbubnis/Zotero/storage/C6HB888N/c3sm52344f.html:text/html;Snapshot:/home/gbubnis/Zotero/storage/8K884E52/c3sm52344f.html:text/html}
}

@article{fuhrmans_coarse-grained_2015,
	title = {Coarse-grained simulation of dynamin-mediated fission},
	volume = {11},
	issn = {1744-683X, 1744-6848},
	url = {http://xlink.rsc.org/?DOI=C4SM02533D},
	doi = {10.1039/C4SM02533D},
	language = {en},
	number = {8},
	urldate = {2016-04-12},
	journal = {Soft Matter},
	author = {Fuhrmans, Marc and Müller, Marcus},
	year = {2015},
	pages = {1464--1480}
}

@article{awasthi_simulations_2016,
	title = {Simulations of {Pore} {Formation} in {Lipid} {Membranes}: {Reaction} {Coordinates}, {Convergence}, {Hysteresis}, and {Finite}-{Size} {Effects}},
	volume = {12},
	issn = {1549-9618},
	shorttitle = {Simulations of {Pore} {Formation} in {Lipid} {Membranes}},
	url = {https://doi.org/10.1021/acs.jctc.6b00369},
	doi = {10.1021/acs.jctc.6b00369},
	abstract = {Transmembrane pores play an important role in various biophysical processes such as membrane permeation, membrane fusion, and antimicrobial peptide activity. In principal, all-atom molecular dynamics (MD) simulations provide an accurate model of pore formation in lipid membranes. However, the free energy landscape of transmembrane pore formation remains poorly understood, partly because potential of mean force (PMF) calculations of pore formation strongly depend on the choice of the reaction coordinate. In this study, we used umbrella sampling to compute PMFs for pore formation using three different reaction coordinates, namely, (i) a coordinate that steers the lipids in the lateral direction away from the pore center, (ii) the distance of a single lipid phosphate group from the membrane center, and (iii) the average water density inside a membrane-spanning cylinder. Our results show that while the three reaction coordinates efficiently form pores in membranes, they suffer from strong hysteresis between pore-opening and pore-closing simulations, suggesting that they do not restrain the systems close to the transition state for pore formation. The two reaction coordinates that act via restraining the lipids lead to more pronounced hysteresis compared with the coordinate acting on the water molecules. By comparing PMFs computed from membranes with different numbers of lipids, we observed significant artifacts from the periodic boundary conditions in small simulation systems. Further analysis suggests that the formation and disruption of a continuous hydrogen-bonding network across the membrane corresponds to the transition state for pore formation. Our study provides molecular insights into the critical steps of transmembrane pore formation, and it may guide the development of efficient reaction coordinates for pore formation.},
	number = {7},
	urldate = {2018-08-30},
	journal = {Journal of Chemical Theory and Computation},
	author = {Awasthi, Neha and Hub, Jochen S.},
	month = jul,
	year = {2016},
	pages = {3261--3269},
	file = {ACS Full Text PDF w/ Links:/home/gbubnis/Zotero/storage/7W396EUS/Awasthi and Hub - 2016 - Simulations of Pore Formation in Lipid Membranes .pdf:application/pdf;ACS Full Text Snapshot:/home/gbubnis/Zotero/storage/VEYL74RY/acs.jctc.html:text/html}
}

@article{hub_probing_2017,
	title = {Probing a {Continuous} {Polar} {Defect}: {A} {Reaction} {Coordinate} for {Pore} {Formation} in {Lipid} {Membranes}},
	volume = {13},
	issn = {1549-9618},
	shorttitle = {Probing a {Continuous} {Polar} {Defect}},
	url = {https://doi.org/10.1021/acs.jctc.7b00106},
	doi = {10.1021/acs.jctc.7b00106},
	abstract = {Various biophysical processes involve the formation of aqueous pores over lipid membranes, including processes of membrane fusion, antimicrobial peptide activity, lipid flip-flop, and membrane permeation. Reliable and efficient free-energy calculations of pore formation using molecular dynamics simulations remained challenging due to the lack of good reaction coordinates (RCs) for pore formation. We present a new RC for pore formation that probes the formation and rupture of a continuous polar defect over the membrane. Potential of mean force (PMF) calculations along the new RC rapidly converge and exhibit no hysteresis between pore-opening and pore-closing pathways, in contrast to calculations based on previous RCs. We show that restraints along the new RC may restrain the system tightly to the transition state of pore formation, rationalizing the absence of hysteresis. We observe that the PMF of pore formation in a tension-free membrane of dimyristoylphosphatidylcholine (DMPC) reveals a free-energy barrier for pore nucleation, confirming a long-hypothesized metastable prepore state. We test the influence of the lipid force field, the cutoff distance used for Lennard-Jones interactions, and the lateral membrane size on the free energies of pore formation. In contrast to PMF calculations based on previous RCs, we find that such parameters have a relatively small influence on the free energies of pore nucleation. However, the metastability of the open pore in DMPC may depend on such parameters. The RC has been implemented into an extension of the GROMACS simulation software. The new RC allows for reliable and computationally efficient free-energy calculations of pore formation in lipid membranes.},
	number = {5},
	urldate = {2018-08-30},
	journal = {Journal of Chemical Theory and Computation},
	author = {Hub, Jochen S. and Awasthi, Neha},
	month = may,
	year = {2017},
	pages = {2352--2366},
	file = {ACS Full Text PDF w/ Links:/home/gbubnis/Zotero/storage/LLDMTGJ4/Hub and Awasthi - 2017 - Probing a Continuous Polar Defect A Reaction Coor.pdf:application/pdf;ACS Full Text Snapshot:/home/gbubnis/Zotero/storage/GRMC9TYF/acs.jctc.html:text/html}
}

@article{ting_metastable_2018,
	title = {Metastable {Prepores} in {Tension}-{Free} {Lipid} {Bilayers}},
	volume = {120},
	url = {https://link.aps.org/doi/10.1103/PhysRevLett.120.128103},
	doi = {10.1103/PhysRevLett.120.128103},
	abstract = {The formation and closure of aqueous pores in lipid bilayers is a key step in various biophysical processes. Large pores are well described by classical nucleation theory, but the free-energy landscape of small, biologically relevant pores has remained largely unexplored. The existence of small and metastable “prepores” was hypothesized decades ago from electroporation experiments, but resolving metastable prepores from theoretical models remained challenging. Using two complementary methods—atomistic simulations and self-consistent field theory of a minimal lipid model—we determine the parameters for which metastable prepores occur in lipid membranes. Both methods consistently suggest that pore metastability depends on the relative volume ratio between the lipid head group and lipid tails: lipids with a larger head-group volume fraction (or shorter saturated tails) form metastable prepores, whereas lipids with a smaller head-group volume fraction (or longer unsaturated tails) form unstable prepores.},
	number = {12},
	urldate = {2018-08-30},
	journal = {Physical Review Letters},
	author = {Ting, Christina L. and Awasthi, Neha and Müller, Marcus and Hub, Jochen S.},
	month = mar,
	year = {2018},
	pages = {128103},
	file = {APS Snapshot:/home/gbubnis/Zotero/storage/35JEGX35/PhysRevLett.120.html:text/html}
}

@article{nakao_effects_2018,
	title = {Effects of {Hydrophilic} {Residues} and {Hydrophobic} {Length} on {Flip}-{Flop} {Promotion} by {Transmembrane} {Peptides}},
	volume = {122},
	issn = {1520-6106},
	url = {https://doi.org/10.1021/acs.jpcb.8b00298},
	doi = {10.1021/acs.jpcb.8b00298},
	abstract = {Peptide-induced phospholipid flip-flop (scrambling) was evaluated using transmembrane model peptides in which the central residue was substituted with various amino acid residues (sequence: Ac-GKK(LA)nXW(LA)nLKKA-CONH2). Peptides with a strongly hydrophilic residue (X = Q, N, or H) had higher scramblase activity than that of other peptides, and the activity was also dependent on the length of the peptides. Peptides with a hydrophobic stretch of 17 residues showed high flip-promotion propensity, whereas those of 21 and 25 residues did not, suggesting that membrane thinning under negative mismatch conditions promotes the flipping. Interestingly, a hydrophobic stretch of 19 residues intensively promoted phospholipid scrambling and membrane leakage. The distinctive characteristics of the peptide were ascribed by long-term molecular dynamics simulation to the arrangement of central glutamine and terminal four lysine residues on the same side of the helix. The combination of simulated and experimental data enables understanding of the mechanisms by which transmembrane helices, and ultimately unidentified scramblases in biomembranes, cause lipid scrambling.},
	number = {15},
	urldate = {2018-05-28},
	journal = {The Journal of Physical Chemistry B},
	author = {Nakao, Hiroyuki and Hayashi, Chihiro and Ikeda, Keisuke and Saito, Hiroaki and Nagao, Hidemi and Nakano, Minoru},
	month = apr,
	year = {2018},
	pages = {4318--4324},
	file = {ACS Full Text PDF w/ Links:/home/gbubnis/Zotero/storage/TNLAWBW5/Nakao et al. - 2018 - Effects of Hydrophilic Residues and Hydrophobic Le.pdf:application/pdf;ACS Full Text Snapshot:/home/gbubnis/Zotero/storage/ITNNJ3V8/acs.jpcb.html:text/html}
}

@article{banushkina_optimal_2016,
	title = {Optimal reaction coordinates},
	volume = {6},
	issn = {1759-0884},
	url = {http://onlinelibrary.wiley.com/doi/10.1002/wcms.1276/abstract},
	doi = {10.1002/wcms.1276},
	abstract = {The dynamic behavior of complex systems with many degrees of freedom is often analyzed by projection onto one or a few reaction coordinates. The dynamics is then described in a simple and intuitive way as diffusion on the associated free-energy profile. In order to use such a picture for a quantitative description of the dynamics one needs to select the coordinate in an optimal way so as to minimize non-Markovian effects due to the projection. For equilibrium dynamics between two boundary states (e.g., a reaction), the optimal coordinate is known as the committor or the pfold coordinate in protein folding studies. While the dynamics projected on the committor is not Markovian, many important quantities of the original multidimensional dynamics on an arbitrarily complex landscape can be computed exactly. In this study, we summarize the derivation of this result, discuss different approaches to determine and validate the committor coordinate, and present three illustrative applications: protein folding, the game of chess, and patient recovery dynamics after kidney transplant. WIREs Comput Mol Sci 2016, 6:748–763. doi: 10.1002/wcms.1276 This article is categorized under:


* Structure and Mechanism {\textgreater} Computational Biochemistry and Biophysics

* Theoretical and Physical Chemistry {\textgreater} Reaction Dynamics and Kinetics

* Theoretical and Physical Chemistry {\textgreater} Statistical Mechanics},
	language = {en},
	number = {6},
	urldate = {2018-03-09},
	journal = {Wiley Interdisciplinary Reviews: Computational Molecular Science},
	author = {Banushkina, Polina V. and Krivov, Sergei V.},
	month = nov,
	year = {2016},
	pages = {748--763},
	file = {Full Text PDF:/home/gbubnis/Zotero/storage/CMJ2TEAT/Banushkina and Krivov - 2016 - Optimal reaction coordinates.pdf:application/pdf;Snapshot:/home/gbubnis/Zotero/storage/CBWXIPCH/abstract.html:text/html}
}

@article{elber_calculating_2017,
	title = {Calculating {Iso}-{Committor} {Surfaces} as {Optimal} {Reaction} {Coordinates} with {Milestoning}},
	volume = {19},
	issn = {1099-4300},
	url = {https://www.ncbi.nlm.nih.gov/pmc/articles/PMC5531205/},
	doi = {10.3390/e19050219},
	abstract = {Reaction coordinates are vital tools for qualitative and quantitative analysis of molecular processes. They provide a simple picture of reaction progress and essential input for calculations of free energies and rates. Iso-committor surfaces are considered the optimal reaction coordinate. We present an algorithm to compute efficiently a sequence of isocommittor surfaces. These surfaces are considered an optimal reaction coordinate. The algorithm analyzes Milestoning results to determine the committor function. It requires only the transition probabilities between the milestones, and not transition times. We discuss the following numerical examples: (i) a transition in the Mueller potential; (ii) a conformational change of a solvated peptide; and (iii) cholesterol aggregation in membranes.},
	number = {5},
	urldate = {2018-03-09},
	journal = {Entropy (Basel, Switzerland)},
	author = {Elber, Ron and Bello-Rivas, Juan M. and Ma, Piao and Cardenas, Alfredo E. and Fathizadeh, Arman},
	month = may,
	year = {2017},
	file = {PubMed Central Full Text PDF:/home/gbubnis/Zotero/storage/BP4MV8GP/Elber et al. - 2017 - Calculating Iso-Committor Surfaces as Optimal Reac.pdf:application/pdf}
}

@article{best_reaction_2005,
	title = {Reaction coordinates and rates from transition paths},
	volume = {102},
	copyright = {© 2005},
	issn = {0027-8424, 1091-6490},
	url = {http://www.pnas.org/content/102/19/6732},
	doi = {10.1073/pnas.0408098102},
	abstract = {The molecular mechanism of a reaction in solution is reflected in its transition-state ensemble and transition paths. We use a Bayesian formula relating the equilibrium and transition-path ensembles to identify transition states, rank reaction coordinates, and estimate rate coefficients. We also introduce a variational procedure to optimize reaction coordinates. The theory is illustrated with applications to protein folding and the dipole reorientation of an ordered water chain inside a carbon nanotube. To describe the folding of a simple model of a three-helix bundle protein, we variationally optimize the weights of a projection onto the matrix of native and nonnative amino acid contacts. The resulting one-dimensional reaction coordinate captures the folding transition state, with formation and packing of helix 2 and 3 constituting the bottleneck for folding.},
	language = {en},
	number = {19},
	urldate = {2018-03-09},
	journal = {Proceedings of the National Academy of Sciences},
	author = {Best, Robert B. and Hummer, Gerhard},
	month = may,
	year = {2005},
	keywords = {carbon nanotubes, chemical kinetics, Grotthuss mechanism, protein folding, transition-state theory},
	pages = {6732--6737},
	file = {Full Text PDF:/home/gbubnis/Zotero/storage/WRQFVSF4/Best and Hummer - 2005 - Reaction coordinates and rates from transition pat.pdf:application/pdf;Snapshot:/home/gbubnis/Zotero/storage/CC4Z7UMG/6732.html:text/html}
}

@article{ma_automatic_2005,
	title = {Automatic {Method} for {Identifying} {Reaction} {Coordinates} in {Complex} {Systems}},
	volume = {109},
	issn = {1520-6106},
	url = {https://doi.org/10.1021/jp045546c},
	doi = {10.1021/jp045546c},
	abstract = {To interpret simulations of a complex system to determine the physical mechanism of a dynamical process, it is necessary to identify the small number of coordinates that distinguish the stable states from the transition states. We develop an automatic method for identifying these degrees of freedom from a database of candidate physical variables. In the method neural networks are used to determine the functional dependence of the probability of committing to a stable state (committor) on a set of coordinates, and a genetic algorithm selects the combination of inputs that yields the best fit. The method enables us to obtain the first set of coordinates that is demonstrably sufficient to specify the transition state of the C7eq→ αR isomerization of the alanine dipeptide in the presence of explicit water molecules. It is revealed that the solute−solvent coupling can be described by a solvent-derived electrostatic torque around one of the main-chain bonds, and the collective, long-ranged nature of this interaction accounts for previous failures to characterize this reaction.},
	number = {14},
	urldate = {2018-03-09},
	journal = {The Journal of Physical Chemistry B},
	author = {Ma, Ao and Dinner, Aaron R.},
	month = apr,
	year = {2005},
	pages = {6769--6779},
	file = {ACS Full Text PDF w/ Links:/home/gbubnis/Zotero/storage/4UNJSBE3/Ma and Dinner - 2005 - Automatic Method for Identifying Reaction Coordina.pdf:application/pdf;ACS Full Text Snapshot:/home/gbubnis/Zotero/storage/X6MN5VPH/jp045546c.html:text/html}
}

@article{gurtovenko_molecular_2007,
	title = {Molecular {Mechanism} for {Lipid} {Flip}-{Flops}},
	volume = {111},
	issn = {1520-6106},
	url = {https://doi.org/10.1021/jp077094k},
	doi = {10.1021/jp077094k},
	abstract = {Transmembrane lipid translocation (flip-flop) processes are involved in a variety of properties and functions of cell membranes, such as membrane asymmetry and programmed cell death. Yet, flip-flops are one of the least understood dynamical processes in membranes. In this work, we elucidate the molecular mechanism of pore-mediated transmembrane lipid translocation (flip-flop) acquired from extensive atomistic molecular dynamics simulations. On the basis of 50 successful flip-flop events resolved in atomic detail, we demonstrate that lipid flip-flops may spontaneously occur in protein-free phospholipid membranes under physiological conditions through transient water pores on a time scale of tens of nanoseconds. While the formation of a water pore is induced here by a transmembrane ion density gradient, the particular way by which the pore is formed is irrelevant for the reported flip-flop mechanism:  the appearance of a transient pore (defect) in the membrane inevitably leads to diffusive translocation of lipids through the pore, which is driven by thermal fluctuations. Our findings strongly support the idea that the formation of membrane defects in terms of water pores is the rate-limiting step in the process of transmembrane lipid flip-flop, which, on average, requires several hours. The findings are consistent with available experimental and computational data and provide a view to interpret experimental observations. For example, the simulation results provide a molecular-level explanation in terms of pores for the experimentally observed fact that the exposure of lipid membranes to electric field pulses considerably reduces the time required for lipid flip-flops.},
	number = {48},
	urldate = {2018-03-08},
	journal = {The Journal of Physical Chemistry B},
	author = {Gurtovenko, Andrey A. and Vattulainen, Ilpo},
	month = dec,
	year = {2007},
	pages = {13554--13559},
	file = {ACS Full Text PDF w/ Links:/home/gbubnis/Zotero/storage/66N9DASG/Gurtovenko and Vattulainen - 2007 - Molecular Mechanism for Lipid Flip-Flops.pdf:application/pdf;ACS Full Text Snapshot:/home/gbubnis/Zotero/storage/C8I3J6NH/jp077094k.html:text/html}
}

@article{bockmann_kinetics_2008,
	title = {Kinetics, {Statistics}, and {Energetics} of {Lipid} {Membrane} {Electroporation} {Studied} by {Molecular} {Dynamics} {Simulations}},
	volume = {95},
	issn = {0006-3495},
	url = {https://www.ncbi.nlm.nih.gov/pmc/articles/PMC2483751/},
	doi = {10.1529/biophysj.108.129437},
	abstract = {Membrane electroporation is the method to directly transfer bioactive substances such as drugs and genes into living cells, as well as preceding electrofusion. Although much information on the microscopic mechanism has been obtained both from experiment and simulation, the existence and nature of possible intermediates is still unclear. To elucidate intermediates of electropore formation by direct comparison with measured prepore formation kinetics, we have carried out 49 atomistic electroporation simulations on a palmitoyl-oleoyl-phosphatidylcholine bilayer for electric field strengths between 0.04 and 0.7 V/nm. A statistical theory is developed to facilitate direct comparison of experimental (macroscopic) prepore formation kinetics with the (single event) preporation times derived from the simulations, which also allows us to extract an effective number of lipids involved in each pore formation event. A linear dependency of the activation energy for prepore formation on the applied field is seen, with quantitative agreement between experiment and simulation. The distribution of preporation times suggests a four-state pore formation model. The model involves a first intermediate characterized by a differential tilt of the polar lipid headgroups on both leaflets, and a second intermediate (prepore), where a polar chain across the bilayer is formed by 3–4 lipid headgroups and several water molecules, thereby providing a microscopic explanation for the polarizable volume derived previously from the measured kinetics. An average pore radius of 0.47 ± 0.15 nm is seen, in favorable agreement with conductance measurements and electrooptical experiments of lipid vesicles.},
	number = {4},
	urldate = {2018-03-08},
	journal = {Biophysical Journal},
	author = {Böckmann, Rainer A. and de Groot, Bert L. and Kakorin, Sergej and Neumann, Eberhard and Grubmüller, Helmut},
	month = aug,
	year = {2008},
	pages = {1837--1850},
	file = {PubMed Central Full Text PDF:/home/gbubnis/Zotero/storage/Q43Z4BU4/Böckmann et al. - 2008 - Kinetics, Statistics, and Energetics of Lipid Memb.pdf:application/pdf}
}

@article{melikov_voltage-induced_2001,
	title = {Voltage-induced nonconductive pre-pores and metastable single pores in unmodified planar lipid bilayer},
	volume = {80},
	issn = {0006-3495},
	doi = {10.1016/S0006-3495(01)76153-X},
	abstract = {Electric fields promote pore formation in both biological and model membranes. We clamped unmodified planar bilayers at 150-550 mV to monitor transient single pores for a long period of time. We observed fast transitions between different conductance levels reflecting opening and closing of metastable lipid pores. Although mean lifetime of the pores was 3 +/- 0.8 ms (250 mV), some pores remained open for up to approximately 1 s. The mean amplitude of conductance fluctuations (approximately 500 pS) was independent of voltage and close for bilayers of different area (40,000 and 10 microm(2)), indicating the local nature of the conductive defects. The distribution of pore conductance was rather broad (dispersion of approximately 250 pS). Based on the conductance value and its dependence of the ion size, the radius of the average pore was estimated as approximately 1 nm. Short bursts of conductance spikes (opening and closing of pores) were often separated by periods of background conductance. Within the same burst the conductance between spikes was indistinguishable from the background. The mean time interval between spikes in the burst was much smaller than that between adjacent bursts. These data indicate that opening and closing of lipidic pores proceed through some electrically invisible (silent) pre-pores. Similar pre-pore defects and metastable conductive pores might be involved in remodeling of cell membranes in different biologically relevant processes.},
	language = {eng},
	number = {4},
	journal = {Biophysical Journal},
	author = {Melikov, K. C. and Frolov, V. A. and Shcherbakov, A. and Samsonov, A. V. and Chizmadzhev, Y. A. and Chernomordik, L. V.},
	month = apr,
	year = {2001},
	keywords = {Electrophysiology, Kinetics, Lipid Bilayers, Membranes, Artificial, Normal Distribution, Patch-Clamp Techniques, Time Factors},
	pages = {1829--1836}
}

@article{chodera_use_2007,
	title = {Use of the {Weighted} {Histogram} {Analysis} {Method} for the {Analysis} of {Simulated} and {Parallel} {Tempering} {Simulations}},
	volume = {3},
	issn = {1549-9618},
	url = {http://dx.doi.org/10.1021/ct0502864},
	doi = {10.1021/ct0502864},
	abstract = {The growing adoption of generalized-ensemble algorithms for biomolecular simulation has resulted in a resurgence in the use of the weighted histogram analysis method (WHAM) to make use of all data generated by these simulations. Unfortunately, the original presentation of WHAM by Kumar et al. is not directly applicable to data generated by these methods. WHAM was originally formulated to combine data from independent samplings of the canonical ensemble, whereas many generalized-ensemble algorithms sample from mixtures of canonical ensembles at different temperatures. Sorting configurations generated from a parallel tempering simulation by temperature obscures the temporal correlation in the data and results in an improper treatment of the statistical uncertainties used in constructing the estimate of the density of states. Here we present variants of WHAM, STWHAM and PTWHAM, derived with the same set of assumptions, that can be directly applied to several generalized ensemble algorithms, including simulated tempering, parallel tempering (better known as replica-exchange among temperatures), and replica-exchange simulated tempering. We present methods that explicitly capture the considerable temporal correlation in sequentially generated configurations using autocorrelation analysis. This allows estimation of the statistical uncertainty in WHAM estimates of expectations for the canonical ensemble. We test the method with a one-dimensional model system and then apply it to the estimation of potentials of mean force from parallel tempering simulations of the alanine dipeptide in both implicit and explicit solvent.},
	number = {1},
	urldate = {2016-08-29},
	journal = {Journal of Chemical Theory and Computation},
	author = {Chodera, John D. and Swope, William C. and Pitera, Jed W. and Seok, Chaok and Dill, Ken A.},
	month = jan,
	year = {2007},
	pages = {26--41},
	file = {ACS Full Text Snapshot:/home/gbubnis/Zotero/storage/U6KIFFND/ct0502864.html:text/html;No Job Name - ct0502864:/home/gbubnis/Zotero/storage/9G4TGEN5/ct0502864.pdf:application/pdf}
}

@article{kirsch_membrane_2016,
	title = {Membrane pore formation in atomistic and coarse-grained simulations},
	volume = {1858},
	issn = {0005-2736},
	url = {http://www.sciencedirect.com/science/article/pii/S000527361500437X},
	doi = {10.1016/j.bbamem.2015.12.031},
	abstract = {Biological cells and their organelles are protected by ultra thin membranes. These membranes accomplish a broad variety of important tasks like separating the cell content from the outer environment, they are the site for cell–cell interactions and many enzymatic reactions, and control the in- and efflux of metabolites.

For certain physiological functions e.g. in the fusion of membranes and also in a number of biotechnological applications like gene transfection the membrane integrity needs to be compromised to allow for instance for the exchange of polar molecules across the membrane barrier. Mechanisms enabling the transport of molecules across the membrane involve membrane proteins that form specific pores or act as transporters, but also so-called lipid pores induced by external fields, stress, or peptides.

Recent progress in the simulation field enabled to closely mimic pore formation as supposed to occur in vivo or in vitro. Here, we review different simulation-based approaches in the study of membrane pores with a focus on lipid pore properties such as their size and energetics, poration mechanisms based on the application of external fields, charge imbalances, or surface tension, and on pores that are induced by small molecules, peptides, and lipids. This article is part of a Special Issue entitled: Biosimulations edited by Ilpo Vattulainen and Tomasz Róg.},
	number = {10},
	urldate = {2016-08-11},
	journal = {Biochimica et Biophysica Acta (BBA) - Biomembranes},
	author = {Kirsch, Sonja A. and Böckmann, Rainer A.},
	month = oct,
	year = {2016},
	keywords = {Atomistic simulation, Coarse-grained simulation, Electroporation, Lipid pore, Membrane pore, molecular dynamics simulation},
	pages = {2266--2277},
	file = {ScienceDirect Full Text PDF:/home/gbubnis/Zotero/storage/57C6MTXI/Kirsch and Böckmann - 2016 - Membrane pore formation in atomistic and coarse-gr.pdf:application/pdf;ScienceDirect Snapshot:/home/gbubnis/Zotero/storage/3EEBN2ZK/S000527361500437X.html:text/html}
}

@article{zhelev_tension-stabilized_1993,
	title = {Tension-stabilized pores in giant vesicles: determination of pore size and pore line tension},
	volume = {1147},
	issn = {0005-2736},
	shorttitle = {Tension-stabilized pores in giant vesicles},
	url = {http://www.sciencedirect.com/science/article/pii/000527369390319U},
	doi = {10.1016/0005-2736(93)90319-U},
	abstract = {We present the first observations of giant, long-existing, stabilized pores in vesicle membranes. Using a new experimental technique for studying the electropermeabilization of lipid membranes, giant liposomes (from 25 to 56 μm in diameter) were subjected to single, square, electric pulses (duration 150 μs and electric field strength from 63 to 126 kV/m). The liposomes were held by a micropipet and small membrane tensions were created by controlling the pipet suction pressure. The liposomes were loaded with media having different refractive index from the outside solution, and, under these conditions, the formation of pores in the pressurized liposome could be visualized by the jet of inside solution that flowed out from the membrane pore. By adjusting the membrane tension, pores were kept open, and pore lifetimes could be varied from tenths of a second to several seconds. The pore size was determined from the volumetric flow in the pore region and the measured pressure differences across the bilayer. It was clear from the experiments that only one pore remained opened after the pulse. The estimated pore radii were on the order of one micrometer. The pores were in a quasi-stationary state and when they closed they did so spontaneously in a quick process (in milliseconds). The isotropic membrane tension was determined for the same measurements and from determinations of both pore size and dynamic membrane tension the pore line tension was found. The line tension of the pore region was determined for two lipid compositions, stearoyloleoylphosphatidylcholine and stearoyloleoylphosphatidylcholine with 50 mol\% cholesterol, and the obtained values for single bilayers were (0.92 ± 0.07)·10−11 N and (3.05 ± 0.12)·10−11 N, respectively.},
	number = {1},
	urldate = {2016-05-17},
	journal = {Biochimica et Biophysica Acta (BBA) - Biomembranes},
	author = {Zhelev, Doncho V. and Needham, David},
	month = apr,
	year = {1993},
	keywords = {Electroporation, Line tension, Liposome, Pore size},
	pages = {89--104},
	file = {1-s2.0-000527369390319U-main.pdf:/home/gbubnis/Zotero/storage/ZGS6XPE7/1-s2.0-000527369390319U-main.pdf:application/pdf;ScienceDirect Snapshot:/home/gbubnis/Zotero/storage/4PBN9XHU/000527369390319U.html:text/html;ScienceDirect Snapshot:/home/gbubnis/Zotero/storage/QXAKE4R3/000527369390319U.html:text/html;Zotero:/home/gbubnis/Zotero/storage/J6GA2HHF/tab.xul:application/vnd.mozilla.xul+xml}
}

@article{leontiadou_molecular_2004,
	title = {Molecular {Dynamics} {Simulations} of {Hydrophilic} {Pores} in {Lipid} {Bilayers}},
	volume = {86},
	issn = {0006-3495},
	url = {http://www.sciencedirect.com/science/article/pii/S0006349504742757},
	doi = {10.1016/S0006-3495(04)74275-7},
	abstract = {Hydrophilic pores are formed in peptide free lipid bilayers under mechanical stress. It has been proposed that the transport of ionic species across such membranes is largely determined by the existence of such meta-stable hydrophilic pores. To study the properties of these structures and understand the mechanism by which pore expansion leads to membrane rupture, a series of molecular dynamics simulations of a dipalmitoylphosphatidylcholine (DPPC) bilayer have been conducted. The system was simulated in two different states; first, as a bilayer containing a meta-stable pore and second, as an equilibrated bilayer without a pore. Surface tension in both cases was applied to study the formation and stability of hydrophilic pores inside the bilayers. It is observed that below a critical threshold tension of ∼38 mN/m the pores are stabilized. The minimum radius at which a pore can be stabilized is 0.7 nm. Based on the critical threshold tension the line tension of the bilayer was estimated to be ∼3 × 10−11 N, in good agreement with experimental measurements. The flux of water molecules through these stabilized pores was analyzed, and the structure and size of the pores characterized. When the lateral pressure exceeds the threshold tension, the pores become unstable and start to expand causing the rupture of the membrane. In the simulations the mechanical threshold tension necessary to cause rupture of the membrane on a nanosecond timescale is much higher in the case of the equilibrated bilayers, as compared with membranes containing preexisting pores.},
	number = {4},
	urldate = {2016-05-17},
	journal = {Biophysical Journal},
	author = {Leontiadou, Hari and Mark, Alan E. and Marrink, Siewert J.},
	month = apr,
	year = {2004},
	pages = {2156--2164},
	file = {ScienceDirect Full Text PDF:/home/gbubnis/Zotero/storage/GQ3MIT9G/Leontiadou et al. - 2004 - Molecular Dynamics Simulations of Hydrophilic Pore.pdf:application/pdf;ScienceDirect Snapshot:/home/gbubnis/Zotero/storage/DPZKDSRW/S0006349504742757.html:text/html}
}

@article{fromherz_lipid-vesicle_1983,
	title = {Lipid-vesicle structure: {Size} control by edge-active agents},
	volume = {94},
	issn = {0009-2614},
	shorttitle = {Lipid-vesicle structure},
	url = {http://www.sciencedirect.com/science/article/pii/0009261483870833},
	doi = {10.1016/0009-2614(83)87083-3},
	abstract = {Fragments of planar lipid bilayers bifurcate into closed vesicles near a critical vesiculation index as determined by their size and by their edge tension. Gradual modulation by edge-active agents — as described in analogy to the Gibbs adsorption equation — leads to the formation of monodisperse vesicles of controlled size.},
	number = {3},
	urldate = {2016-05-12},
	journal = {Chemical Physics Letters},
	author = {Fromherz, Peter},
	month = jan,
	year = {1983},
	pages = {259--266},
	file = {ScienceDirect Snapshot:/home/gbubnis/Zotero/storage/J34WA679/0009261483870833.html:text/html}
}

@article{warner_evolution_2012,
	title = {Evolution of the {Hemifused} {Intermediate} on the {Pathway} to {Membrane} {Fusion}},
	volume = {103},
	issn = {0006-3495},
	url = {http://www.sciencedirect.com/science/article/pii/S0006349512007308},
	doi = {10.1016/j.bpj.2012.06.041},
	abstract = {The pathway to membrane fusion in synthetic and biological systems is thought to pass through hemifusion, in which the outer leaflets are fused while the inner leaflets engage in a hemifusion diaphragm (HD). Fusion has been proposed to be completed by lysis of the expanded HD that matures from a localized stalklike initial connection. However, the process that establishes the expanded HD is poorly understood. Here we mathematically modeled hemifusion of synthetic vesicles, where hemifusion and fusion are most commonly driven by calcium and membrane tension. The model shows that evolution of the hemifused state is driven by these agents and resisted by interleaflet frictional and tensile stresses. Predicted HD growth rates depend on tension and salt concentration, and agree quantitatively with experimental measurements. For typical conditions, we predict that HDs expand at ∼30 μm2/s, reaching a final equilibrium area ∼7\% of the vesicle area. Key model outputs are the evolving HD tension and area during the growth transient, properties that may determine whether HD lysis occurs. Applying the model to numerous published experimental studies that reported fusion, our results are consistent with a final fusion step in which the HD ruptures due to super-lysis HD membrane tensions.},
	number = {4},
	urldate = {2016-05-12},
	journal = {Biophysical Journal},
	author = {Warner, Jason M. and O’Shaughnessy, Ben},
	month = aug,
	year = {2012},
	pages = {689--701},
	file = {ScienceDirect Full Text PDF:/home/gbubnis/Zotero/storage/AAUHDFT5/Warner and O’Shaughnessy - 2012 - Evolution of the Hemifused Intermediate on the Pat.pdf:application/pdf;ScienceDirect Snapshot:/home/gbubnis/Zotero/storage/B37WDWCV/S0006349512007308.html:text/html}
}

@article{evans_dynamic_2003,
	title = {Dynamic {Tension} {Spectroscopy} and {Strength} of {Biomembranes}},
	volume = {85},
	issn = {0006-3495},
	url = {http://www.sciencedirect.com/science/article/pii/S000634950374658X},
	doi = {10.1016/S0006-3495(03)74658-X},
	abstract = {Rupturing fluid membrane vesicles with a steady ramp of micropipette suction produces a distribution of breakage tensions governed by the kinetic process of membrane failure. When plotted as a function of log(tension loading rate), the locations of distribution peaks define a dynamic tension spectrum with distinct regimes that reflect passage of prominent energy barriers along the kinetic pathway. Using tests on five types of giant phosphatidylcholine lipid vesicles over loading rates(tension/time) from 0.01–100 mN/m/s, we show that the kinetic process of membrane breakage can be modeled by a causal sequence of two thermally-activated transitions. At fast loading rates, a steep linear regime appears in each spectrum which implies that membrane failure starts with nucleation of a rare precursor defect. The slope and projected intercept of this regime are set by defect size and frequency of spontaneous formation, respectively. But at slow loading rates, each spectrum crosses over to a shallow-curved regime where rupture tension changes weakly with rate. This regime is predicted by the classical cavitation theory for opening an unstable hole in a two-dimensional film within the lifetime of the defect state. Under slow loading, membrane edge energy and the frequency scale for thermal fluctuations in hole size are the principal factors that govern the level of tension at failure. To critically test the model and obtain the parameters governing the rates of transition under stress, distributions of rupture tension were computed and matched to the measured histograms through solution of the kinetic master (Markov) equations for defect formation and annihilation or evolution to an unstable hole under a ramp of tension. As key predictors of membrane strength, the results for spontaneous frequencies of defect formation and hole edge energies were found to correlate with membrane thicknesses and elastic bending moduli, respectively.},
	number = {4},
	urldate = {2016-05-12},
	journal = {Biophysical Journal},
	author = {Evans, Evan and Heinrich, Volkmar and Ludwig, Florian and Rawicz, Wieslawa},
	month = oct,
	year = {2003},
	pages = {2342--2350},
	file = {ScienceDirect Full Text PDF:/home/gbubnis/Zotero/storage/VAH5AWUA/Evans et al. - 2003 - Dynamic Tension Spectroscopy and Strength of Biome.pdf:application/pdf;ScienceDirect Snapshot:/home/gbubnis/Zotero/storage/WXKTDQ3A/S000634950374658X.html:text/html}
}

@article{risselada_free_2014,
	title = {Free {Energy} {Landscape} of {Rim}-{Pore} {Expansion} in {Membrane} {Fusion}},
	volume = {107},
	issn = {0006-3495},
	url = {http://www.ncbi.nlm.nih.gov/pmc/articles/PMC4241460/},
	doi = {10.1016/j.bpj.2014.08.022},
	abstract = {The productive fusion pore in membrane fusion is generally thought to be toroidally shaped. Theoretical studies and recent experiments suggest that its formation, in some scenarios, may be preceded by an initial pore formed near the rim of the extended hemifusion diaphragm (HD), a rim-pore. This rim-pore is characterized by a nontoroidal shape that changes with size. To determine this shape as well as the free energy along the pathway of rim-pore expansion, we derived a simple analytical free energy model. We argue that dilation of HD material via expansion of a rim-pore is favored over a regular, circular pore. Further, the expanding rim-pore faces a free energy barrier that linearly increases with HD size. In contrast, the tension required to expand the rim-pore decreases with HD size. Pore flickering, followed by sudden opening, occurs when the tension in the HD competes with the line energy of the rim-pore, and the rim-pore reaches its equilibrium size before reaching the critical pore size. The experimental observation of flickering and closing fusion pores (kiss-and-run) is very well explained by the observed behavior of rim-pores. Finally, the free energy landscape of rim-pore expansion/HD dilation may very well explain why some cellular fusion reactions, in their attempt to minimize energetic costs, progress via alternative formation and dilation of microscopic hemifusion intermediates.},
	number = {10},
	urldate = {2016-05-12},
	journal = {Biophysical Journal},
	author = {Risselada, Herre Jelger and Smirnova, Yuliya and Grubmüller, Helmut},
	month = nov,
	year = {2014},
	pages = {2287--2295},
	file = {PubMed Central Full Text PDF:/home/gbubnis/Zotero/storage/8ZAZDTZH/Risselada et al. - 2014 - Free Energy Landscape of Rim-Pore Expansion in Mem.pdf:application/pdf}
}

@article{akimov_pore_2017,
	title = {Pore formation in lipid membrane {I}: {Continuous} reversible trajectory from intact bilayer through hydrophobic defect to transversal pore},
	volume = {7},
	copyright = {2017 The Author(s)},
	issn = {2045-2322},
	shorttitle = {Pore formation in lipid membrane {I}},
	url = {https://www.nature.com/articles/s41598-017-12127-7},
	doi = {10.1038/s41598-017-12127-7},
	abstract = {Lipid membranes serve as effective barriers allowing cells to maintain internal composition differing from that of extracellular medium. Membrane permeation, both natural and artificial, can take place via appearance of transversal pores. The rearrangements of lipids leading to pore formation in the intact membrane are not yet understood in details. We applied continuum elasticity theory to obtain continuous trajectory of pore formation and closure, and analyzed molecular dynamics trajectories of pre-formed pore reseal. We hypothesized that a transversal pore is preceded by a hydrophobic defect: intermediate structure spanning through the membrane, the side walls of which are partially aligned by lipid tails. This prediction was confirmed by our molecular dynamics simulations. Conversion of the hydrophobic defect into the hydrophilic pore required surmounting some energy barrier. A metastable state was found for the hydrophilic pore at the radius of a few nanometers. The dependence of the energy on radius was approximately quadratic for hydrophobic defect and small hydrophilic pore, while for large radii it depended on the radius linearly. The pore energy related to its perimeter, line tension, thus depends of the pore radius. Calculated values of the line tension for large pores were in quantitative agreement with available experimental data.},
	language = {En},
	number = {1},
	urldate = {2019-05-01},
	journal = {Scientific Reports},
	author = {Akimov, Sergey A. and Volynsky, Pavel E. and Galimzyanov, Timur R. and Kuzmin, Peter I. and Pavlov, Konstantin V. and Batishchev, Oleg V.},
	month = sep,
	year = {2017},
	pages = {12152},
	file = {Full Text PDF:/home/gbubnis/Zotero/storage/PRM6XRLM/Akimov et al. - 2017 - Pore formation in lipid membrane I Continuous rev.pdf:application/pdf;Snapshot:/home/gbubnis/Zotero/storage/A23KVH6B/s41598-017-12127-7.html:text/html}
}

@article{quaytman_reaction_2007,
	title = {Reaction coordinate of an enzymatic reaction revealed by transition path sampling},
	volume = {104},
	copyright = {© 2007 by The National Academy of Sciences of the USA},
	issn = {0027-8424, 1091-6490},
	url = {https://www.pnas.org/content/104/30/12253},
	doi = {10.1073/pnas.0704304104},
	abstract = {The transition path sampling method previously applied in our group to the reaction catalyzed by lactate dehydrogenase was used to generate a transition path ensemble for this reaction. Based on analysis of the reactive trajectories generated, important residues behind the active site were implicated in a compressional motion that brought the donor–acceptor atoms of the hydride closer together. In addition, residues behind the active site were implicated in a relaxational motion, locking the substrate in product formation. Although this suggested that the compression-relaxation motions of these residues were important to catalysis, it remained unproven. In this work, we used committor distribution analysis to show that these motions are integral components of the reaction coordinate.},
	language = {en},
	number = {30},
	urldate = {2019-05-01},
	journal = {Proceedings of the National Academy of Sciences},
	author = {Quaytman, Sara L. and Schwartz, Steven D.},
	month = jul,
	year = {2007},
	keywords = {catalysis, protein dynamics},
	pages = {12253--12258},
	file = {Full Text PDF:/home/gbubnis/Zotero/storage/4MCQSQBB/Quaytman and Schwartz - 2007 - Reaction coordinate of an enzymatic reaction revea.pdf:application/pdf;Snapshot:/home/gbubnis/Zotero/storage/4XRNHFTM/12253.html:text/html}
}

@article{chodera_splitting_2011,
	title = {Splitting probabilities as a test of reaction coordinate choice in single-molecule experiments},
	volume = {107},
	issn = {1079-7114},
	doi = {10.1103/PhysRevLett.107.098102},
	abstract = {To explain the observed dynamics in equilibrium single-molecule measurements of biomolecules, the experimental observable is often chosen as a putative reaction coordinate along which kinetic behavior is presumed to be governed by diffusive dynamics. Here, we invoke the splitting probability as a test of the suitability of such a proposed reaction coordinate. Comparison of the observed splitting probability with that computed from the kinetic model provides a simple test to reject poor reaction coordinates. We demonstrate this test for a force spectroscopy measurement of a DNA hairpin.},
	language = {eng},
	number = {9},
	journal = {Physical Review Letters},
	author = {Chodera, John D. and Pande, Vijay S.},
	month = aug,
	year = {2011},
	keywords = {DNA, Kinetics, Macromolecular Substances, Models, Chemical, Nucleic Acid Conformation},
	pages = {098102},
	file = {Accepted Version:/home/gbubnis/Zotero/storage/JMCDH989/Chodera and Pande - 2011 - Splitting probabilities as a test of reaction coor.pdf:application/pdf}
}

@article{tan_theory_2012,
	title = {Theory of binless multi-state free energy estimation with applications to protein-ligand binding},
	volume = {136},
	issn = {1089-7690},
	doi = {10.1063/1.3701175},
	abstract = {The weighted histogram analysis method (WHAM) is routinely used for computing free energies and expectations from multiple ensembles. Existing derivations of WHAM require observations to be discretized into a finite number of bins. Yet, WHAM formulas seem to hold even if the bin sizes are made arbitrarily small. The purpose of this article is to demonstrate both the validity and value of the multi-state Bennet acceptance ratio (MBAR) method seen as a binless extension of WHAM. We discuss two statistical arguments to derive the MBAR equations, in parallel to the self-consistency and maximum likelihood derivations already known for WHAM. We show that the binless method, like WHAM, can be used not only to estimate free energies and equilibrium expectations, but also to estimate equilibrium distributions. We also provide a number of useful results from the statistical literature, including the determination of MBAR estimators by minimization of a convex function. This leads to an approach to the computation of MBAR free energies by optimization algorithms, which can be more effective than existing algorithms. The advantages of MBAR are illustrated numerically for the calculation of absolute protein-ligand binding free energies by alchemical transformations with and without soft-core potentials. We show that binless statistical analysis can accurately treat sparsely distributed interaction energy samples as obtained from unmodified interaction potentials that cannot be properly analyzed using standard binning methods. This suggests that binless multi-state analysis of binding free energy simulations with unmodified potentials offers a straightforward alternative to the use of soft-core potentials for these alchemical transformations.},
	language = {eng},
	number = {14},
	journal = {The Journal of Chemical Physics},
	author = {Tan, Zhiqiang and Gallicchio, Emilio and Lapelosa, Mauro and Levy, Ronald M.},
	month = apr,
	year = {2012},
	keywords = {Ligands, Likelihood Functions, Models, Molecular, Protein Binding, Proteins, Thermodynamics},
	pages = {144102},
	file = {Full Text:/home/gbubnis/Zotero/storage/YKGEFARB/Tan et al. - 2012 - Theory of binless multi-state free energy estimati.pdf:application/pdf}
}

@article{peters_obtaining_2006,
	title = {Obtaining reaction coordinates by likelihood maximization},
	volume = {125},
	issn = {0021-9606},
	url = {https://aip.scitation.org/doi/abs/10.1063/1.2234477},
	doi = {10.1063/1.2234477},
	number = {5},
	urldate = {2019-05-01},
	journal = {The Journal of Chemical Physics},
	author = {Peters, Baron and Trout, Bernhardt L.},
	month = aug,
	year = {2006},
	pages = {054108},
	file = {Snapshot:/home/gbubnis/Zotero/storage/T79F8A8M/1.html:text/html}
}

@article{li_recent_2014,
	title = {Recent developments in methods for identifying reaction coordinates},
	volume = {40},
	issn = {0892-7022},
	doi = {10.1080/08927022.2014.907898},
	abstract = {In the study of rare events in complex systems with many degrees of freedom, a key element is to identify the reaction coordinates of a given process. Over recent years, a number of methods and protocols have been developed to extract the reaction coordinates based on limited information from molecular dynamics simulations. In this review, we provide a brief survey over a number of major methods developed in the past decade, some of which are discussed in greater detail, to provide an overview of the problems that are partially solved and challenges that still remain. A particular emphasis has been placed on methods for identifying reaction coordinates that are related to the committor.},
	language = {eng},
	number = {10-11},
	journal = {Molecular Simulation},
	author = {Li, Wenjin and Ma, Ao},
	year = {2014},
	keywords = {committor, molecular dynamics, rare events, reaction coordinate},
	pages = {784--793},
	file = {Accepted Version:/home/gbubnis/Zotero/storage/UBBMBCU9/Li and Ma - 2014 - Recent developments in methods for identifying rea.pdf:application/pdf}
}

@article{kopelevich_one-dimensional_2013,
	title = {One-dimensional potential of mean force underestimates activation barrier for transport across flexible lipid membranes},
	volume = {139},
	issn = {0021-9606},
	url = {https://aip.scitation.org/doi/abs/10.1063/1.4823500},
	doi = {10.1063/1.4823500},
	number = {13},
	urldate = {2019-05-01},
	journal = {The Journal of Chemical Physics},
	author = {Kopelevich, Dmitry I.},
	month = oct,
	year = {2013},
	pages = {134906},
	file = {Snapshot:/home/gbubnis/Zotero/storage/5KDQ8Q7V/1.html:text/html}
}

@article{chodera_splitting_2011-1,
	title = {Splitting {Probabilities} as a {Test} of {Reaction} {Coordinate} {Choice} in {Single}-{Molecule} {Experiments}},
	volume = {107},
	url = {https://link.aps.org/doi/10.1103/PhysRevLett.107.098102},
	doi = {10.1103/PhysRevLett.107.098102},
	abstract = {To explain the observed dynamics in equilibrium single-molecule measurements of biomolecules, the experimental observable is often chosen as a putative reaction coordinate along which kinetic behavior is presumed to be governed by diffusive dynamics. Here, we invoke the splitting probability as a test of the suitability of such a proposed reaction coordinate. Comparison of the observed splitting probability with that computed from the kinetic model provides a simple test to reject poor reaction coordinates. We demonstrate this test for a force spectroscopy measurement of a DNA hairpin.},
	number = {9},
	urldate = {2019-05-09},
	journal = {Physical Review Letters},
	author = {Chodera, John D. and Pande, Vijay S.},
	month = aug,
	year = {2011},
	pages = {098102},
	file = {APS Snapshot:/home/gbubnis/Zotero/storage/FNQRI8GQ/PhysRevLett.107.html:text/html;Full Text PDF:/home/gbubnis/Zotero/storage/YGISJKHF/Chodera and Pande - 2011 - Splitting Probabilities as a Test of Reaction Coor.pdf:application/pdf}
}

@article{abidor_electric_1979,
	title = {Electric breakdown of bilayer lipid membranes: {I}. {The} main experimental facts and their qualitative discussion},
	volume = {104},
	issn = {0022-0728},
	shorttitle = {Electric breakdown of bilayer lipid membranes},
	url = {http://www.sciencedirect.com/science/article/pii/S0022072879810062},
	doi = {10.1016/S0022-0728(79)81006-2},
	abstract = {The process of the electric breakdown of bilayer lipid membranes is studied experimentally. The mean lifetime of the membranes is measured as a function of the membrane potential difference. The distribution functions for the membrane lifetime are obtained for various values of the applied voltage. A qualitative interpretation of the breakdown phenomenon is proposed, which is based upon the concept of structural defects of the type of throughgoing pores, whose development is favoured by the electric field. These pores are analogous to the nuclei in the theory of new phase formation. The elucidation of the mechanism of BLM breakdown in the electricfield may provide an important tool for studying the electric breakdown of biological membranes and possibly the nature of background conductivity of membranes.},
	language = {en},
	urldate = {2020-05-18},
	journal = {Journal of Electroanalytical Chemistry and Interfacial Electrochemistry},
	author = {Abidor, I. G. and Arakelyan, V. B. and Chernomordik, L. V. and Chizmadzhev, Yu. A. and Pastushenko, V. F. and Tarasevich, M. P.},
	month = jan,
	year = {1979},
	pages = {37--52},
	file = {ScienceDirect Snapshot:/home/gbubnis/Zotero/storage/WHBG7I3V/S0022072879810062.html:text/html}
}

@article{onsager_initial_1938,
	title = {Initial {Recombination} of {Ions}},
	volume = {54},
	url = {https://link.aps.org/doi/10.1103/PhysRev.54.554},
	doi = {10.1103/PhysRev.54.554},
	abstract = {The probability that a pair of ions of given initial separation will recombine with each other is computed from the laws of Brownian motion, which is the proper procedure whenever the Langevin factor equals unity, as in gases at high pressures. In the absence of forces other than the Coulomb attraction, the probability of escape equals the reciprocal of the Boltzmann factor. This result includes the correlation between temperature and pressure coefficients of the ionization by light particles previously predicted by Compton, Bennett and Stearns, if one allows their basic hypothesis about the laws which govern the initial separation of the ions. The effect of an electric field is to increase the fraction of escaping ions by a factor which in the incipient stage of the effect is proportional to the field intensity and independent of the initial distance, although it depends on the orientation of an ion pair. The predicted increase of the ionization current is a little more than one percent for every 100 volts/cm, which accounts for the observed effects of fields exceeding 1500 volts/cm. A reasonable amount of columnar recombination would help to explain the proportionately greater effects of weak fields. The inferred initial separations of the ions are apparently compatible with present knowledge of electron scattering and attachment.},
	number = {8},
	urldate = {2020-05-18},
	journal = {Physical Review},
	author = {Onsager, L.},
	month = oct,
	year = {1938},
	pages = {554--557},
	file = {APS Snapshot:/home/gbubnis/Zotero/storage/9B7W5EGD/PhysRev.54.html:text/html}
}

@article{peters_reaction_2016,
	title = {Reaction {Coordinates} and {Mechanistic} {Hypothesis} {Tests}},
	volume = {67},
	issn = {1545-1593},
	doi = {10.1146/annurev-physchem-040215-112215},
	abstract = {Reaction coordinates are integral to several classic rate theories that can (a) predict kinetic trends across conditions and homologous reactions, (b) extract activation parameters with a clear physical interpretation from experimental rates, and (c) enable efficient calculations of free energy barriers and rates. New trajectory-based rare events methods can provide rates directly from dynamical trajectories without a reaction coordinate. Trajectory-based frameworks can also generate ideal (but abstract) reaction coordinates such as committors and eigenfunctions of the master equation. However, rates and mechanistic insights obtained from trajectory-based methods and abstract coordinates are not readily generalized across simulation conditions or reaction families. We discuss methods for identifying physically meaningful reaction coordinates, including committor analysis, variational transition state theory, Kramers-Langer-Berezhkovskii-Szabo theory, and statistical inference methods that can use path sampling data to screen, mix, and optimize thousands of trial coordinates. Special focus is given to likelihood maximization and inertial likelihood maximization approaches.},
	language = {eng},
	journal = {Annual Review of Physical Chemistry},
	author = {Peters, Baron},
	year = {2016},
	keywords = {collective variable, committor, dimensionality reduction, likelihood maximization, transition path sampling, transmission coefficient},
	pages = {669--690}
}

@article{peters_using_2006,
	title = {Using the histogram test to quantify reaction coordinate error},
	volume = {125},
	issn = {0021-9606},
	url = {https://aip.scitation.org/doi/abs/10.1063/1.2409924},
	doi = {10.1063/1.2409924},
	number = {24},
	urldate = {2020-05-18},
	journal = {The Journal of Chemical Physics},
	author = {Peters, Baron},
	month = dec,
	year = {2006},
	pages = {241101},
	file = {Snapshot:/home/gbubnis/Zotero/storage/VTAM36EB/1.html:text/html}
}

@article{du_transition_1998,
	title = {On the transition coordinate for protein folding},
	volume = {108},
	issn = {0021-9606},
	url = {https://aip.scitation.org/doi/10.1063/1.475393},
	doi = {10.1063/1.475393},
	number = {1},
	urldate = {2020-05-19},
	journal = {The Journal of Chemical Physics},
	author = {Du, Rose and Pande, Vijay S. and Grosberg, Alexander Yu. and Tanaka, Toyoichi and Shakhnovich, Eugene S.},
	month = jan,
	year = {1998},
	pages = {334--350},
	file = {Snapshot:/home/gbubnis/Zotero/storage/IYMSIIAU/1.html:text/html}
}

@article{grafmuller_free_2014,
	title = {The free energy of nanopores in tense membranes},
	volume = {16},
	issn = {1463-9084},
	url = {https://pubs.rsc.org/en/content/articlelanding/2014/cp/c3cp54685c},
	doi = {10.1039/C3CP54685C},
	abstract = {Membrane nanopores are central players for a range of important cellular membrane remodeling processes as well as membrane rupture. Understanding pore formation in tense membranes requires comprehension of the molecular mechanism of pore formation and the associated free energy change as a function of the membrane tension. Here we propose a scheme to calculate the free energy change associated with the formation of a nanometer sized pore in molecular dynamics simulations as a function of membrane tension, which requires the calculation of only one computationally expensive potential of mean force. We show that membrane elastic theory can be used to estimate the pore formation free energy at different tension values from the free energy change in a relaxed membrane and the area expansion curves of the membranes. We have computed the pore formation free energy for a dipalmitoyl-phosphatidylcholine (DPPC) membrane at two different lateral pressure values, 1 bar and −40 bar, by calculating the potential of mean force acting on the head group of a single lipid molecule. Unrestrained simulations of the closing process confirm that the intermediate states along this reaction coordinate are reasonable and show that hydrophilic indentations spanning half the bilayer connected by a hydrophobic pore segment represent the corresponding high energy transition state. A comparison of the stability of simulated membranes to experiment at high loading rates show that, contrary to expectation, pores form too easily in small simulated membrane patches. This discrepancy originates from a combination of the absence of ions in the simulations and the small membrane size.},
	language = {en},
	number = {23},
	urldate = {2020-05-19},
	journal = {Physical Chemistry Chemical Physics},
	author = {Grafmüller, Andrea and Knecht, Volker},
	month = may,
	year = {2014},
	pages = {11270--11278},
	file = {Full Text PDF:/home/gbubnis/Zotero/storage/NZYMR3ZL/Grafmüller and Knecht - 2014 - The free energy of nanopores in tense membranes.pdf:application/pdf;Snapshot:/home/gbubnis/Zotero/storage/AR4FSRG7/c3cp54685c.html:text/html}
}

@article{wohlert_free_2006,
	title = {Free energy of a trans-membrane pore calculated from atomistic molecular dynamics simulations},
	volume = {124},
	issn = {0021-9606},
	url = {https://aip.scitation.org/doi/10.1063/1.2171965},
	doi = {10.1063/1.2171965},
	number = {15},
	urldate = {2020-05-19},
	journal = {The Journal of Chemical Physics},
	author = {Wohlert, J. and den Otter, W. K. and Edholm, O. and Briels, W. J.},
	month = apr,
	year = {2006},
	pages = {154905},
	file = {Snapshot:/home/gbubnis/Zotero/storage/FS4DSIZB/1.html:text/html}
}

@article{tieleman_lipids_2006,
	title = {Lipids {Out} of {Equilibrium}:  {Energetics} of {Desorption} and {Pore} {Mediated} {Flip}-{Flop}},
	volume = {128},
	issn = {0002-7863},
	shorttitle = {Lipids {Out} of {Equilibrium}},
	url = {https://doi.org/10.1021/ja0624321},
	doi = {10.1021/ja0624321},
	abstract = {The potential of mean force (PMF) of a phospholipid in a bilayer is a key thermodynamic property that describes the energetic cost of localized lipid defects. We have calculated the PMF by umbrella sampling using molecular dynamics simulations. The profile has a deep minimum at the equilibrium position in the bilayer and steeply rises for displacements both deeper into the bilayer and moving away from the bilayer. As the lipid loses contact with the bilayer, the profile abruptly flattens without a significant barrier. The calculated free energy difference of 80 kJ/mol between the minimum of the PMF and the value in water agrees well with the free energy difference calculated from the experimentally measured critical micelle concentration. Significant water/lipid defects form when a lipid is forced into the bilayer interior, in the form of a small water pore that spans the membrane. The energy required to form such a water pore is also found to be 80 kJ/mol. On the basis of this energy, we estimate the lipid flip-flop rate and permeability rate of sodium ions. The resulting rates are in good agreement with experimental measurements, suggesting lipid flip-flop and basal permeability of ions are pore mediated.},
	number = {38},
	urldate = {2020-05-19},
	journal = {Journal of the American Chemical Society},
	author = {Tieleman, D. Peter and Marrink, Siewert-Jan},
	month = sep,
	year = {2006},
	pages = {12462--12467},
	file = {Full Text:/home/gbubnis/Zotero/storage/7ITHQM3V/Tieleman and Marrink - 2006 - Lipids Out of Equilibrium  Energetics of Desorpti.pdf:application/pdf}
}

@article{litster_stability_1975,
	title = {Stability of lipid bilayers and red blood cell membranes},
	volume = {53},
	issn = {0375-9601},
	url = {http://www.sciencedirect.com/science/article/pii/0375960175904028},
	doi = {10.1016/0375-9601(75)90402-8},
	abstract = {It is suggested that Helfrich's concept of edge energy of a lipid bilayer explains the stability of black membranes and the ability of erythrocytes to reseal following hemolysis.},
	language = {en},
	number = {3},
	urldate = {2020-05-19},
	journal = {Physics Letters A},
	author = {Litster, J. D.},
	month = jun,
	year = {1975},
	pages = {193--194},
	file = {ScienceDirect Snapshot:/home/gbubnis/Zotero/storage/BCHKHHQ7/0375960175904028.html:text/html}
}

@article{melikov_voltage-induced_2001-1,
	title = {Voltage-{Induced} {Nonconductive} {Pre}-{Pores} and {Metastable} {Single} {Pores} in {Unmodified} {Planar} {Lipid} {Bilayer}},
	volume = {80},
	issn = {0006-3495},
	url = {https://www.cell.com/biophysj/abstract/S0006-3495(01)76153-X},
	doi = {10.1016/S0006-3495(01)76153-X},
	language = {English},
	number = {4},
	urldate = {2020-05-19},
	journal = {Biophysical Journal},
	author = {Melikov, Kamran C. and Frolov, Vadim A. and Shcherbakov, Arseniy and Samsonov, Andrey V. and Chizmadzhev, Yury A. and Chernomordik, Leonid V.},
	month = apr,
	year = {2001},
	pages = {1829--1836},
	file = {Full Text PDF:/home/gbubnis/Zotero/storage/5CXE4T3G/Melikov et al. - 2001 - Voltage-Induced Nonconductive Pre-Pores and Metast.pdf:application/pdf;Snapshot:/home/gbubnis/Zotero/storage/VWM32ZQW/S0006-3495(01)76153-X.html:text/html}
}

@article{essmann_smooth_1995,
	title = {A smooth particle mesh {Ewald} method},
	volume = {103},
	issn = {0021-9606},
	url = {https://aip.scitation.org/doi/abs/10.1063/1.470117},
	doi = {10.1063/1.470117},
	number = {19},
	urldate = {2020-05-25},
	journal = {The Journal of Chemical Physics},
	author = {Essmann, Ulrich and Perera, Lalith and Berkowitz, Max L. and Darden, Tom and Lee, Hsing and Pedersen, Lee G.},
	month = nov,
	year = {1995},
	pages = {8577--8593},
	file = {Snapshot:/home/gbubnis/Zotero/storage/B3UKCNAF/1.html:text/html;Submitted Version:/home/gbubnis/Zotero/storage/HWKBR2B8/Essmann et al. - 1995 - A smooth particle mesh Ewald method.pdf:application/pdf}
}

@article{berger_molecular_1997,
	title = {Molecular dynamics simulations of a fluid bilayer of dipalmitoylphosphatidylcholine at full hydration, constant pressure, and constant temperature.},
	volume = {72},
	issn = {0006-3495},
	url = {https://www.ncbi.nlm.nih.gov/pmc/articles/PMC1184396/},
	abstract = {Molecular dynamics simulations of 500 ps were performed on a system consisting of a bilayer of 64 molecules of the lipid dipalmitoylphosphatidylcholine and 23 water molecules per lipid at an isotropic pressure of 1 atm and 50 degrees C. Special attention was devoted to reproduce the correct density of the lipid, because this quantity is known experimentally with a precision better than 1\%. For this purpose, the Lennard-Jones parameters of the hydrocarbon chains were adjusted by simulating a system consisting of 128 pentadecane molecules and varying the Lennard-Jones parameters until the experimental density and heat of vaporization were obtained. With these parameters the lipid density resulted in perfect agreement with the experimental density. The orientational order parameter of the hydrocarbon chains agreed perfectly well with the experimental values, which, because of its correlation with the area per lipid, makes it possible to give a proper estimate of the area per lipid of 0.61 +/- 0.01 nm2.},
	number = {5},
	urldate = {2020-05-25},
	journal = {Biophysical Journal},
	author = {Berger, O and Edholm, O and Jähnig, F},
	month = may,
	year = {1997},
	pages = {2002--2013},
	file = {PubMed Central Full Text PDF:/home/gbubnis/Zotero/storage/68CP4NEY/Berger et al. - 1997 - Molecular dynamics simulations of a fluid bilayer .pdf:application/pdf}
}

@incollection{berendsen_interaction_1981,
	address = {Dordrecht},
	series = {The {Jerusalem} {Symposia} on {Quantum} {Chemistry} and {Biochemistry}},
	title = {Interaction {Models} for {Water} in {Relation} to {Protein} {Hydration}},
	isbn = {978-94-015-7658-1},
	url = {https://doi.org/10.1007/978-94-015-7658-1_21},
	abstract = {For molecular dynamics simulations of hydrated proteins a simple yet reliable model for the intermolecular potential for water is required. Such a model must be an effective pair potential valid for liquid densities that takes average many-body interactions into account. We have developed a three-point charge model (on hydrogen and oxygen positions) with a Lennard-Jones 6–12 potential on the oxygen positions only. Parameters for the model were determined from 12 molecular dynamics runs covering the two-dimensional parameter space of charge and oxygen repulsion. Both potential energy and pressure were required to coincide with experimental values. The model has very satisfactory properties, is easily incorporated into protein-water potentials, and requires only 0.25 sec computertime per dynamics step (for 216 molecules) on a CRAY-1 computer.},
	language = {en},
	urldate = {2020-05-25},
	booktitle = {Intermolecular {Forces}: {Proceedings} of the {Fourteenth} {Jerusalem} {Symposium} on {Quantum} {Chemistry} and {Biochemistry} {Held} in {Jerusalem}, {Israel}, {April} 13–16, 1981},
	publisher = {Springer Netherlands},
	author = {Berendsen, H. J. C. and Postma, J. P. M. and van Gunsteren, W. F. and Hermans, J.},
	editor = {Pullman, Bernard},
	year = {1981},
	keywords = {Effective Pair Potential, Mole Cular Cage, Oxygen Position, Protein Hydration, Spectral CORREl Density},
	pages = {331--342}
}

@article{abraham_gromacs_2015,
	title = {{GROMACS}: {High} performance molecular simulations through multi-level parallelism from laptops to supercomputers},
	volume = {1-2},
	issn = {2352-7110},
	shorttitle = {{GROMACS}},
	url = {http://www.sciencedirect.com/science/article/pii/S2352711015000059},
	doi = {10.1016/j.softx.2015.06.001},
	abstract = {GROMACS is one of the most widely used open-source and free software codes in chemistry, used primarily for dynamical simulations of biomolecules. It provides a rich set of calculation types, preparation and analysis tools. Several advanced techniques for free-energy calculations are supported. In version 5, it reaches new performance heights, through several new and enhanced parallelization algorithms. These work on every level; SIMD registers inside cores, multithreading, heterogeneous CPU–GPU acceleration, state-of-the-art 3D domain decomposition, and ensemble-level parallelization through built-in replica exchange and the separate Copernicus framework. The latest best-in-class compressed trajectory storage format is supported.},
	language = {en},
	urldate = {2020-05-25},
	journal = {SoftwareX},
	author = {Abraham, Mark James and Murtola, Teemu and Schulz, Roland and Páll, Szilárd and Smith, Jeremy C. and Hess, Berk and Lindahl, Erik},
	month = sep,
	year = {2015},
	keywords = {Molecular dynamics, SIMD, GPU, Free energy},
	pages = {19--25},
	file = {ScienceDirect Full Text PDF:/home/gbubnis/Zotero/storage/8DMNMDW3/Abraham et al. - 2015 - GROMACS High performance molecular simulations th.pdf:application/pdf;ScienceDirect Snapshot:/home/gbubnis/Zotero/storage/H84BEH6K/S2352711015000059.html:text/html}
}

@article{berendsen_molecular_1984,
	title = {Molecular dynamics with coupling to an external bath},
	volume = {81},
	issn = {0021-9606},
	url = {https://aip.scitation.org/doi/10.1063/1.448118},
	doi = {10.1063/1.448118},
	number = {8},
	urldate = {2020-05-25},
	journal = {The Journal of Chemical Physics},
	author = {Berendsen, H. J. C. and Postma, J. P. M. and van Gunsteren, W. F. and DiNola, A. and Haak, J. R.},
	month = oct,
	year = {1984},
	pages = {3684--3690},
	file = {Snapshot:/home/gbubnis/Zotero/storage/L9AWH4BG/1.html:text/html;Submitted Version:/home/gbubnis/Zotero/storage/NABNUNX8/Berendsen et al. - 1984 - Molecular dynamics with coupling to an external ba.pdf:application/pdf}
}

@article{goga_efficient_2012,
	title = {Efficient {Algorithms} for {Langevin} and {DPD} {Dynamics}},
	volume = {8},
	issn = {1549-9618},
	url = {https://doi.org/10.1021/ct3000876},
	doi = {10.1021/ct3000876},
	abstract = {In this article, we present several algorithms for stochastic dynamics, including Langevin dynamics and different variants of Dissipative Particle Dynamics (DPD), applicable to systems with or without constraints. The algorithms are based on the impulsive application of friction and noise, thus avoiding the computational complexity of algorithms that apply continuous friction and noise. Simulation results on thermostat strength and diffusion properties for ideal gas, coarse-grained (MARTINI) water, and constrained atomic (SPC/E) water systems are discussed. We show that the measured thermal relaxation rates agree well with theoretical predictions. The influence of various parameters on the diffusion coefficient is discussed.},
	number = {10},
	urldate = {2020-05-25},
	journal = {Journal of Chemical Theory and Computation},
	author = {Goga, N. and Rzepiela, A. J. and de Vries, A. H. and Marrink, S. J. and Berendsen, H. J. C.},
	month = oct,
	year = {2012},
	pages = {3637--3649},
	file = {ACS Full Text Snapshot:/home/gbubnis/Zotero/storage/V5MX7MJ5/ct3000876.html:text/html;Full Text PDF:/home/gbubnis/Zotero/storage/FKTI2FMA/Goga et al. - 2012 - Efficient Algorithms for Langevin and DPD Dynamics.pdf:application/pdf}
}

@article{miyamoto_settle_1992,
	title = {Settle: {An} analytical version of the {SHAKE} and {RATTLE} algorithm for rigid water models},
	volume = {13},
	copyright = {Copyright © 1992 John Wiley \& Sons, Inc.},
	issn = {1096-987X},
	shorttitle = {Settle},
	url = {https://onlinelibrary.wiley.com/doi/abs/10.1002/jcc.540130805},
	doi = {10.1002/jcc.540130805},
	abstract = {An analytical algorithm, called SETTLE, for resetting the positions and velocities to satisfy the holonomic constraints on the rigid water model is presented. This method is still based on the Cartesian coordinate system and can be used in place of SHAKE and RATTLE. We implemented this algorithm in the SPASMS package of molecular mechanics and dynamics. Several series of molecular dynamics simulations were carried out to examine the performance of the new algorithm in comparison with the original RATTLE method. It was found that SETTLE is of higher accuracy and is faster than RATTLE with reasonable tolerances by three to nine times on a scalar machine. Furthermore, the performance improvement ranged from factors of 26 to 98 on a vector machine since the method presented is not iterative. © 1992 by John Wiley \& Sons, Inc.},
	language = {en},
	number = {8},
	urldate = {2020-05-25},
	journal = {Journal of Computational Chemistry},
	author = {Miyamoto, Shuichi and Kollman, Peter A.},
	year = {1992},
	pages = {952--962},
	file = {Snapshot:/home/gbubnis/Zotero/storage/XRHXZFW6/jcc.html:text/html}
}

@article{hess_lincs_1997,
	title = {{LINCS}: {A} linear constraint solver for molecular simulations},
	volume = {18},
	copyright = {Copyright © 1997 John Wiley \& Sons, Inc.},
	issn = {1096-987X},
	shorttitle = {{LINCS}},
	url = {https://onlinelibrary.wiley.com/doi/abs/10.1002/%28SICI%291096-987X%28199709%2918%3A12%3C1463%3A%3AAID-JCC4%3E3.0.CO%3B2-H},
	doi = {10.1002/(SICI)1096-987X(199709)18:12<1463::AID-JCC4>3.0.CO;2-H},
	abstract = {In this article, we present a new LINear Constraint Solver (LINCS) for molecular simulations with bond constraints. The algorithm is inherently stable, as the constraints themselves are reset instead of derivatives of the constraints, thereby eliminating drift. Although the derivation of the algorithm is presented in terms of matrices, no matrix matrix multiplications are needed and only the nonzero matrix elements have to be stored, making the method useful for very large molecules. At the same accuracy, the LINCS algorithm is three to four times faster than the SHAKE algorithm. Parallelization of the algorithm is straightforward. © 1997 John Wiley \& Sons, Inc. J Comput Chem 18: 1463–1472, 1997},
	language = {en},
	number = {12},
	urldate = {2020-05-25},
	journal = {Journal of Computational Chemistry},
	author = {Hess, Berk and Bekker, Henk and Berendsen, Herman J. C. and Fraaije, Johannes G. E. M.},
	year = {1997},
	keywords = {molecular dynamics, constraints, Langevin dynamics, SHAKE},
	pages = {1463--1472},
	file = {Snapshot:/home/gbubnis/Zotero/storage/KEUH7GRF/(SICI)1096-987X(199709)18121463AID-JCC43.0.html:text/html}
}

@article{knight_memgen_2015,
	title = {{MemGen}: a general web server for the setup of lipid membrane simulation systems},
	volume = {31},
	issn = {1367-4803},
	shorttitle = {{MemGen}},
	url = {https://academic.oup.com/bioinformatics/article/31/17/2897/183942},
	doi = {10.1093/bioinformatics/btv292},
	abstract = {Abstract.  Motivation: Molecular dynamics simulations provide atomic insight into the physicochemical characteristics of lipid membranes and hence, a wide range},
	language = {en},
	number = {17},
	urldate = {2020-05-25},
	journal = {Bioinformatics},
	author = {Knight, Christopher J. and Hub, Jochen S.},
	month = sep,
	year = {2015},
	pages = {2897--2899},
	file = {Full Text PDF:/home/gbubnis/Zotero/storage/DGG8XYWF/Knight and Hub - 2015 - MemGen a general web server for the setup of lipi.pdf:application/pdf;Snapshot:/home/gbubnis/Zotero/storage/SRLSB8HA/183942.html:text/html}
}

@article{marti_molecular_2004,
	title = {A molecular dynamics transition path sampling study of model lipid bilayer membranes in aqueous environments},
	volume = {16},
	issn = {0953-8984},
	url = {https://doi.org/10.1088%2F0953-8984%2F16%2F32%2F004},
	doi = {10.1088/0953-8984/16/32/004},
	abstract = {The transversal motion of lipids across biological membranes or flip–flop in aqueous media is of fundamental importance in understanding chemical reactions produced in cells. It is also an example of a rare event in a biological system. Such a process has been investigated by molecular dynamics transition path sampling, which is a powerful tool designed to deal with rare events in a wide range of systems. The study covers a wide range of temperature conditions and includes the analysis of flip–flop free energies together with a detailed study of vibrational densities of states associated with flip–flop motions.},
	language = {en},
	number = {32},
	urldate = {2020-05-26},
	journal = {Journal of Physics: Condensed Matter},
	author = {Martí, J.},
	month = aug,
	year = {2004},
	pages = {5669--5678}
}

@article{alabi_perspectives_2013,
	title = {Perspectives on {Kiss}-and-{Run}: {Role} in {Exocytosis}, {Endocytosis}, and {Neurotransmission}},
	volume = {75},
	issn = {0066-4278},
	shorttitle = {Perspectives on {Kiss}-and-{Run}},
	url = {https://www.annualreviews.org/doi/10.1146/annurev-physiol-020911-153305},
	doi = {10.1146/annurev-physiol-020911-153305},
	abstract = {Regulated exocytosis and endocytosis are critical to the function of many intercellular networks, particularly the complex neural circuits underlying mammalian behavior. Kiss-and-run (KR) is an unconventional fusion between secretory vesicles and a target membrane that releases intravesicular content through a transient, nanometer-sized fusion pore. The fusing vesicle retains its gross shape, precluding full integration into the planar membrane, and enough molecular components for rapid retrieval, reacidification, and reuse. KR makes judicious use of finite presynaptic resources, and mounting evidence suggests that it influences synaptic information transfer. Here we detail emerging perspectives on KR and its role in neurotransmission. We additionally formulate a restraining force hypothesis as a plausible mechanistic basis for KR and its physiological modulation in small nerve terminals. Clarification of the mechanism and function of KR has bearing on understanding the kinetic transitions underlying SNARE-mediated fusion, interactions between vesicles and their local environment, and the influence of release dynamics on neural information processing.},
	number = {1},
	urldate = {2020-05-27},
	journal = {Annual Review of Physiology},
	author = {Alabi, AbdulRasheed A. and Tsien, Richard W.},
	month = feb,
	year = {2013},
	pages = {393--422},
	file = {Full Text PDF:/home/gbubnis/Zotero/storage/LZVVSI77/Alabi and Tsien - 2013 - Perspectives on Kiss-and-Run Role in Exocytosis, .pdf:application/pdf;Snapshot:/home/gbubnis/Zotero/storage/BB3HEGR9/annurev-physiol-020911-153305.html:text/html}
}

@article{smirnova_free-energy_2015,
	title = {Free-energy calculation methods for collective phenomena in membranes},
	volume = {48},
	issn = {0022-3727},
	url = {https://doi.org/10.1088%2F0022-3727%2F48%2F34%2F343001},
	doi = {10.1088/0022-3727/48/34/343001},
	abstract = {Collective phenomena in membranes are those which involve the co-operative reorganization of many molecules. Examples of these are membrane fusion, pore formation, bending, adhesion or fission. The time and length scales, on which these processes occur, pose a challenge for atomistic simulations. Therefore, in order to solve the length scale problem it is popular to introduce a coarse-grained representation. To facilitate sampling of the relevant states additional computational techniques, which encourage the system to explore the free-energy landscape far from equilibrium and visit transition states, are needed. These computational techniques provide insights about the free-energy changes involved in collective transformations of membranes, yielding information about the rate limiting states, the transformation mechanism and the influence of architectural, compositional and interaction parameters. A common approach is to identify an order parameter (or reaction coordinate), which characterizes the pathway of membrane reorganization. However, no general strategy exists to define such an order parameter that can properly describe cooperative reorganizations in membranes. Recently developed methods can overcome this problem of the order-parameter choice and allow us to study collective phenomena in membranes. We will discuss such methods as thermodynamic integration, umbrella sampling, and the string method and results provided by their applications to particle-based simulations, particularly focusing on membrane fusion and pore formation.},
	language = {en},
	number = {34},
	urldate = {2020-05-27},
	journal = {Journal of Physics D: Applied Physics},
	author = {Smirnova, Yuliya G. and Fuhrmans, Marc and Vidal, Israel A. Barragan and Müller, Marcus},
	month = jul,
	year = {2015},
	pages = {343001}
}

@article{bruckner_flip-flop-induced_2009,
	title = {Flip-flop-induced relaxation of bending energy: implications for membrane remodeling},
	volume = {97},
	issn = {1542-0086},
	shorttitle = {Flip-flop-induced relaxation of bending energy},
	doi = {10.1016/j.bpj.2009.09.025},
	abstract = {Cellular and organellar membranes are dynamic materials that underlie many aspects of cell biology. Biological membranes have long been thought of as elastic materials with respect to bending deformations. A wealth of theory and experimentation on pure phospholipid membranes provides abundant support for this idea. However, biological membranes are not composed solely of phospholipids--they also incorporate a variety of amphiphilic molecules that undergo rapid transbilayer flip-flop. Here we describe several experimental systems that demonstrate deformation-induced molecular flip-flop. First we use a fluorescence assay to track osmotically controlled membrane deformation in single component fatty acid vesicles, and show that the relaxation of the induced bending stress is mediated by fatty acid flip-flop. We then look at two-component phospholipid/cholesterol composite vesicles. We use NMR to show that the steady-state rate of interleaflet diffusion of cholesterol is fast relative to biological membrane remodeling. We then use a Förster resonance energy transfer assay to detect the transbilayer movement of cholesterol upon deformation. We suggest that our results can be interpreted by modifying the area difference elasticity model to account for the time-dependent relaxation of bending energy. Our findings suggest that rapid interleaflet diffusion of cholesterol may play a role in membrane remodeling in vivo. We suggest that the molecular characteristics of sterols make them evolutionarily preferred mediators of stress relaxation, and that the universal presence of sterols in the membranes of eukaryotes, even at low concentrations, reflects the importance of membrane remodeling in eukaryotic cells.},
	language = {eng},
	number = {12},
	journal = {Biophysical Journal},
	author = {Bruckner, R. J. and Mansy, S. S. and Ricardo, A. and Mahadevan, L. and Szostak, J. W.},
	month = dec,
	year = {2009},
	keywords = {2-Naphthylamine, Biomechanical Phenomena, Cell Membrane, Cholesterol, Diffusion, Elasticity, Endocytosis, Eukaryota, Evolution, Molecular, Fatty Acids, Fluorescence Resonance Energy Transfer, Fluorescent Dyes, Kinetics, Laurates, Models, Biological, Movement, Osmotic Pressure, Stress, Mechanical, Thermodynamics},
	pages = {3113--3122},
	file = {Full Text:/home/gbubnis/Zotero/storage/AR7HCQRP/Bruckner et al. - 2009 - Flip-flop-induced relaxation of bending energy im.pdf:application/pdf}
}

@article{fathizadeh_ion_2019,
	title = {Ion {Permeation} through a {Phospholipid} {Membrane}: {Transition} {State}, {Path} {Splitting}, and {Calculation} of {Permeability}},
	volume = {15},
	issn = {1549-9618},
	shorttitle = {Ion {Permeation} through a {Phospholipid} {Membrane}},
	url = {https://doi.org/10.1021/acs.jctc.8b00882},
	doi = {10.1021/acs.jctc.8b00882},
	abstract = {We investigate the thermodynamics and kinetics of the permeation of a potassium ion through a phospholipid membrane. We illustrate that the conventional reaction coordinate (the position of the ion along the normal to the membrane plane) is insufficient to capture essential elements of the process. It is necessary to add coarse variables that measure membrane distortion. New coarse variables are suggested, and a two-dimensional coarse-space is proposed to describe the permeation. We illustrate path splitting and two transition states of comparable barrier heights. The alternative pathways differ by the extent of water solvation of the ion-phosphate pairs. The permeation process cannot be described by a local one-dimensional reaction coordinate, and a network formulation is more appropriate. We use Milestoning with Voronoi tessellation in two dimensions to quantify the equilibrium and rate of the permeation of the positively charged ion. The permeation coefficient is computed and compared favorably to experiment.},
	number = {1},
	urldate = {2020-05-27},
	journal = {Journal of Chemical Theory and Computation},
	author = {Fathizadeh, Arman and Elber, Ron},
	month = jan,
	year = {2019},
	pages = {720--730}
}

@article{enkavi_multiscale_2019,
	title = {Multiscale {Simulations} of {Biological} {Membranes}: {The} {Challenge} {To} {Understand} {Biological} {Phenomena} in a {Living} {Substance}},
	volume = {119},
	issn = {0009-2665},
	shorttitle = {Multiscale {Simulations} of {Biological} {Membranes}},
	url = {https://doi.org/10.1021/acs.chemrev.8b00538},
	doi = {10.1021/acs.chemrev.8b00538},
	abstract = {Biological membranes are tricky to investigate. They are complex in terms of molecular composition and structure, functional over a wide range of time scales, and characterized by nonequilibrium conditions. Because of all of these features, simulations are a great technique to study biomembrane behavior. A significant part of the functional processes in biological membranes takes place at the molecular level; thus computer simulations are the method of choice to explore how their properties emerge from specific molecular features and how the interplay among the numerous molecules gives rise to function over spatial and time scales larger than the molecular ones. In this review, we focus on this broad theme. We discuss the current state-of-the-art of biomembrane simulations that, until now, have largely focused on a rather narrow picture of the complexity of the membranes. Given this, we also discuss the challenges that we should unravel in the foreseeable future. Numerous features such as the actin-cytoskeleton network, the glycocalyx network, and nonequilibrium transport under ATP-driven conditions have so far received very little attention; however, the potential of simulations to solve them would be exceptionally high. A major milestone for this research would be that one day we could say that computer simulations genuinely research biological membranes, not just lipid bilayers.},
	number = {9},
	urldate = {2020-05-27},
	journal = {Chemical Reviews},
	author = {Enkavi, Giray and Javanainen, Matti and Kulig, Waldemar and Róg, Tomasz and Vattulainen, Ilpo},
	month = may,
	year = {2019},
	pages = {5607--5774},
	file = {ACS Full Text Snapshot:/home/gbubnis/Zotero/storage/9ZFMWFJW/acs.chemrev.html:text/html;Full Text PDF:/home/gbubnis/Zotero/storage/TP6KV8BZ/Enkavi et al. - 2019 - Multiscale Simulations of Biological Membranes Th.pdf:application/pdf}
}

@article{van_meer_membrane_2008,
	title = {Membrane lipids: where they are and how they behave},
	volume = {9},
	copyright = {2008 Nature Publishing Group},
	issn = {1471-0080},
	shorttitle = {Membrane lipids},
	url = {https://www.nature.com/articles/nrm2330},
	doi = {10.1038/nrm2330},
	abstract = {Lipids function as essential structural components of membranes, as signalling molecules, as chemical identifiers of specific membranes and as energy storage molecules. The synthesis of lipids is non-uniformly distributed among a few cellular membranes, which requires most organelles to depend on lipid transport processes to achieve their full complement of lipids.Organelles differ both quantitatively and qualitatively in their lipid content. In many organelles, the lipid composition of separate leaflets of the membrane bilayer is significantly different, which produces asymmetry across the bilayer — a situation that is maintained by ATP-dependent flippases.Lipid transport between and within organelles is poorly understood, but a growing number of genes that are involved in these processes have been unambiguously identified, and their mechanisms of action are under active investigation. There is clear evidence that several lipids are transported between organelles by non-vesicular mechanisms that involve zones of apposition between donor and acceptor compartments, and macromolecular assemblies that involve multiple lipids and proteins.Several dozen lipids participate in intra- and intercellular signalling processes. In most instances, the levels of signalling molecules are exceedingly low compared with the complement of structural lipids that is present in membranes.Lipids adopt defined phases depending on their molecular structure and the physical conditions. In lipid mixtures, two fluid phases can coexist with different physical properties: liquid-disordered and liquid-ordered.Liquid-ordered assemblies in biomembranes, known as lipid rafts, are small and transient but can coalesce and become stabilized during signalling and vesicle budding. How proteins contribute to phase separation and preferentially distribute into one of the two different phases (or at their interface) is presently unclear.},
	language = {en},
	number = {2},
	urldate = {2020-05-27},
	journal = {Nature Reviews Molecular Cell Biology},
	author = {van Meer, Gerrit and Voelker, Dennis R. and Feigenson, Gerald W.},
	month = feb,
	year = {2008},
	pages = {112--124},
	file = {Full Text:/home/gbubnis/Zotero/storage/ABW6DVDZ/van Meer et al. - 2008 - Membrane lipids where they are and how they behav.pdf:application/pdf;Snapshot:/home/gbubnis/Zotero/storage/F2J7Z98M/nrm2330.html:text/html}
}

@article{burger_greasing_2000,
	title = {Greasing {Membrane} {Fusion} and {Fission} {Machineries}},
	volume = {1},
	issn = {1600-0854},
	url = {https://onlinelibrary.wiley.com/doi/abs/10.1034/j.1600-0854.2000.010804.x},
	doi = {10.1034/j.1600-0854.2000.010804.x},
	abstract = {Biological membrane fusion is a local-point event, extremely fast, and under strict control. Proteins are responsible for the mutual recognition of the fusion partners and for the initiation of biomembrane fusion, and thus determine where and when fusion occurs. However, the central event during membrane fusion is the merger of two membranes, which requires a transient reorganization of membrane lipids into highly curved fusion intermediates. This review focuses on the potential role of lipids in the generation of membrane curvature, and thus in the regulation of membrane fusion and fission.},
	language = {en},
	number = {8},
	urldate = {2020-05-27},
	journal = {Traffic},
	author = {Burger, Koert N. J.},
	year = {2000},
	keywords = {Fission, fusion, lipid polymorphism, lysophosphatidic acid, membrane curvature, phosphatidic acid, type-II lipids},
	pages = {605--613},
	file = {Full Text PDF:/home/gbubnis/Zotero/storage/LJDDWUD2/Burger - 2000 - Greasing Membrane Fusion and Fission Machineries.pdf:application/pdf;Snapshot:/home/gbubnis/Zotero/storage/KSEKGDSI/j.1600-0854.2000.010804.html:text/html}
}

@article{deamer_permeability_1986,
	series = {Special {Issue}: {Liposomes}},
	title = {Permeability of lipid bilayers to water and ionic solutes},
	volume = {40},
	issn = {0009-3084},
	url = {http://www.sciencedirect.com/science/article/pii/0009308486900691},
	doi = {10.1016/0009-3084(86)90069-1},
	abstract = {The lipid bilayer moiety of biological membranes is considered to be the primary barrier to free diffusion of water and solutes. This conclusion arises from observations of lipid bilayer model membrane systems, which are generally less permeable than biological membranes. However, the nature of the permeability barrier remains unclear, particularly with respect to ionic solutes. For instance, anion permeability is significantly greater than cation permeability, and permeability to proton-hydroxide is orders of magnitude greater than other monovalent inorganic ions. In this review, we first consider bilayer permeability to water and discuss proposed permeation mechanisms which involve transient defects arising from thermal fluctuations. We next consider whether such defects can account for ion permeation, including proton-hydroxide flux. We conclude that at least two varieties of transient defects are required to explain permeation of water and ionic solutes.},
	language = {en},
	number = {2},
	urldate = {2020-05-27},
	journal = {Chemistry and Physics of Lipids},
	author = {Deamer, David W. and Bramhall, John},
	month = jun,
	year = {1986},
	keywords = {gramicidin A, ion flux, liposomes, permeability, planar lipid membranes, proton flux, water flux},
	pages = {167--188},
	file = {ScienceDirect Snapshot:/home/gbubnis/Zotero/storage/EIZL6T97/0009308486900691.html:text/html}
}

@article{jahn_membrane_1999,
	title = {Membrane {Fusion} and {Exocytosis}},
	volume = {68},
	issn = {0066-4154},
	url = {https://www.annualreviews.org/doi/10.1146/annurev.biochem.68.1.863},
	doi = {10.1146/annurev.biochem.68.1.863},
	abstract = {Membrane fusion involves the merger of two phospholipid bilayers in an aqueous environment. In artificial lipid bilayers, fusion proceeds by means of defined transition states, including hourglass-shaped intermediates in which the proximal leaflets of the fusing membranes are merged whereas the distal leaflets are separate (fusion stalk), followed by the reversible opening of small aqueous fusion pores. Fusion of biological membranes requires the action of specific fusion proteins. Best understood are the viral fusion proteins that are responsible for merging the viral with the host cell membrane during infection. These proteins undergo spontaneous and dramatic conformational changes upon activation. In the case of the paradigmatic fusion proteins of the influenza virus and of the human immunodeficiency virus, an amphiphilic fusion peptide is inserted into the target membrane. The protein then reorients itself, thus forcing the fusing membranes together and inducing lipid mixing. Fusion of intracellular membranes in eukaryotic cells involves several protein families including SNAREs, Rab proteins, and Sec1/Munc-18 related proteins (SM-proteins). SNAREs form a novel superfamily of small and mostly membrane-anchored proteins that share a common motif of about 60 amino acids (SNARE motif). SNAREs reversibly assemble into tightly packed helical bundles, the core complexes. Assembly is thought to pull the fusing membranes closely together, thus inducing fusion. SM-proteins comprise a family of soluble proteins that bind to certain types of SNAREs and prevent the formation of core complexes. Rab proteins are GTPases that undergo highly regulated GTP-GDP cycles. In their GTP form, they interact with specific proteins, the effector proteins. Recent evidence suggests that Rab proteins function in the initial membrane contact connecting the fusing membranes but are not involved in the fusion reaction itself.},
	number = {1},
	urldate = {2020-05-27},
	journal = {Annual Review of Biochemistry},
	author = {Jahn, Reinhard and Südhof, Thomas C.},
	month = jun,
	year = {1999},
	pages = {863--911},
	file = {Full Text PDF:/home/gbubnis/Zotero/storage/U4NFA5IZ/Jahn and Südhof - 1999 - Membrane Fusion and Exocytosis.pdf:application/pdf;Snapshot:/home/gbubnis/Zotero/storage/9LCUI9MD/annurev.biochem.68.1.html:text/html}
}

@article{reddy_antimicrobial_2004,
	title = {Antimicrobial peptides: premises and promises},
	volume = {24},
	issn = {0924-8579},
	shorttitle = {Antimicrobial peptides},
	url = {http://www.sciencedirect.com/science/article/pii/S092485790400322X},
	doi = {10.1016/j.ijantimicag.2004.09.005},
	abstract = {Antimicrobial peptides (AMPs) are an important component of the natural defences of most living organisms against invading pathogens. These are relatively small ({\textless}10kDa), cationic and amphipathic peptides of variable length, sequence and structure. During the past two decades several AMPs have been isolated from a wide variety of animals, both vertebrates and invertebrates, and plants as well as from bacteria and fungi. Most of these peptides are obtained from different sources like macrophages, neutrophils, epithelial cells, haemocytes, fat body, reproductive tract, etc. These peptides exhibit broad-spectrum activity against a wide range of microorganisms including Gram-positive and Gram-negative bacteria, protozoa, yeast, fungi and viruses. A few peptides have also been found to be cytotoxic to sperm and tumour cells. AMPs are classified based on the three dimensional structural studies carried out with the help of NMR. The peptides are broadly classified into five major groups namely (a) peptides that form α-helical structures, (b) peptides rich in cysteine residues, (c) peptides that form β-sheet, (d) peptides rich in regular amino acids namely histatin, arginine and proline and (e) peptides composed of rare and modified amino acids. Most of these peptides are believed to act by disrupting the plasma membrane leading to the lysis of the cell. AMPs have been found to be excellent candidates for developing novel antimicrobial agents and a few of these peptides show antimicrobial activity against pathogens causing sexually transmitted infection (STI), including HIV/HSV. Peptides, namely magainin and nisin have been shown to demonstrate contraceptive properties in vitro and in vivo. A few peptides have already entered clinical trials for the treatment of impetigo, diabetic foot ulcers and gastric helicobacter infections. In this review, we discuss the source, structures and mode of action with special reference to therapeutic considerations of various AMPs.},
	language = {en},
	number = {6},
	urldate = {2020-05-27},
	journal = {International Journal of Antimicrobial Agents},
	author = {Reddy, K. V. R. and Yedery, R. D. and Aranha, C.},
	month = dec,
	year = {2004},
	keywords = {Antimicrobial peptides, Microbicides, Sexually transmitted infections},
	pages = {536--547},
	file = {ScienceDirect Snapshot:/home/gbubnis/Zotero/storage/XMDL9S8L/S092485790400322X.html:text/html}
}

@article{shai_mode_2002,
	title = {Mode of action of membrane active antimicrobial peptides},
	volume = {66},
	copyright = {Copyright © 2002 Wiley Periodicals, Inc.},
	issn = {1097-0282},
	url = {https://onlinelibrary.wiley.com/doi/abs/10.1002/bip.10260},
	doi = {10.1002/bip.10260},
	abstract = {Water-membrane soluble protein and peptide toxins are used in the defense and offense systems of all organisms, including plants and humans. A major group includes antimicrobial peptides, which serve as a nonspecific defense system that complements the highly specific cell-mediated immune response. The increasing resistance of bacteria to conventional antibiotics stimulated the isolation and characterization of many antimicrobial peptides for potential use as new target antibiotics. The finding of thousands of antimicrobial peptides with variable lengths and sequences, all of which are active at similar concentrations, suggests a general mechanism for killing bacteria rather than a specific mechanism that requires preferred active structures. Such a mechanism is in agreement with the “carpet model” that does not require any specific structure or sequence. It seems that when there is an appropriate balance between hydrophobicity and a net positive charge the peptides are active on bacteria. However, selective activity depends also on other parameters, such as the volume of the molecule, its structure, and its oligomeric state in solution and membranes. Further, although many studies support that bacterial membrane damage is a lethal event for bacteria, other studies point to a multihit mechanism in which the peptide binds to several targets in the cytoplasmic region of the bacteria.},
	language = {en},
	number = {4},
	urldate = {2020-05-27},
	journal = {Peptide Science},
	author = {Shai, Yechiel},
	year = {2002},
	keywords = {antibacterial peptides, antimicrobial peptides, diasteromeric cationic peptides, lytic peptides, peptide–lipid interaction},
	pages = {236--248},
	file = {Snapshot:/home/gbubnis/Zotero/storage/5DP5R69T/bip.html:text/html}
}

@article{neumann_gene_1982,
	title = {Gene transfer into mouse lyoma cells by electroporation in high electric fields},
	volume = {1},
	issn = {0261-4189},
	abstract = {Electric impulses (8 kV/cm, 5 microseconds) were found to increase greatly the uptake of DNA into cells. When linear or circular plasmid DNA containing the herpes simplex thymidine kinase (TK) gene is added to a suspension of mouse L cells deficient in the TK gene and the cells are then exposed to electric fields, stable transformants are formed that survive in the HAT selection medium. At 20 degrees C after the application of three successive electric impulses followed by 10 min to allow DNA entry there result 95 (+/- 3) transformants per 10(6) cells and per 1.2 micrograms DNA. Compared with biochemical techniques, the electric field method of gene transfer is very simple, easily applicable, and very efficient. Because the mechanism of DNA transport through cell membranes is not known, a simple physical model for the enhanced DNA penetration into cells in high electric fields is proposed. According to this ' electroporation model' the interaction of the external electric field with the lipid dipoles of a pore configuration induces and stabilizes the permeation sites and thus enhances cross membrane transport.},
	language = {eng},
	number = {7},
	journal = {The EMBO journal},
	author = {Neumann, E. and Schaefer-Ridder, M. and Wang, Y. and Hofschneider, P. H.},
	year = {1982},
	keywords = {Animals, Cloning, Molecular, Electric Stimulation, Genes, Genes, Viral, Kinetics, L Cells, Mice, Plasmids, Simplexvirus, Thermodynamics, Thymidine Kinase},
	pages = {841--845}
}

@article{nishizawa_molecular_2013,
	title = {Molecular dynamics simulation analysis of membrane defects and pore propensity of hemifusion diaphragms},
	volume = {104},
	issn = {1542-0086},
	doi = {10.1016/j.bpj.2013.01.022},
	abstract = {Membrane fusion often exhibits slow dynamics in electrophysiological experiments, involving prespike foot and fusion pore-flickering, but the structural basis of such phenomena remains unclear. Hemifusion intermediates have been implicated in the early phase of membrane fusion. To elucidate the dynamics of formation of membrane defects and pores within the hemifusion diaphragm (HD), atomistic and coarse-grained models of hemifusion intermediates were constructed using dipalmitoylphosphatidylcholine or dioleoylphosphatidylcholine membranes. The work necessary to displace a lipid molecule to the hydrophobic core of the bilayer was measured. For a lipid within the HD with radius of 4 nm, the work was ∼80 kJ/mol, similar to that in a planar bilayer. The work was much less (∼40 kJ/mol) when the HD was surrounded by a steep stalk, i.e., stalk wings forming a large angle at the junction of three bilayers. In the latter case, the lipid displacement engendered formation of a pore contacting the HD rim. The work was similarly small (40 kJ/mol) for a small HD of 1.5 nm radius, where a pore formed and grew rapidly, quickly generating a toroidal structure ({\textless}40 ns). Combining the steep stalk and the small HD decreased the work further, although quantitative analysis was difficult because the latter system was not in a stable equilibrium state. Results suggest that fine tuning of fusion dynamics requires strict control of the HD size and the angle between the expanded stalk and HD. In additional free simulations, the steep stalk facilitated widening of a preformed pore contacting the HD rim.},
	language = {eng},
	number = {5},
	journal = {Biophysical Journal},
	author = {Nishizawa, Manami and Nishizawa, Kazuhisa},
	month = mar,
	year = {2013},
	keywords = {Cell Membrane, Lipid Bilayers, Membrane Fusion, Molecular Dynamics Simulation},
	pages = {1038--1048},
	file = {Full Text:/home/gbubnis/Zotero/storage/YILUGE9X/Nishizawa and Nishizawa - 2013 - Molecular dynamics simulation analysis of membrane.pdf:application/pdf}
}

@article{ketterer_dna_2018,
	title = {{DNA} origami scaffold for studying intrinsically disordered proteins of the nuclear pore complex},
	volume = {9},
	copyright = {2018 The Author(s)},
	issn = {2041-1723},
	url = {https://www.nature.com/articles/s41467-018-03313-w},
	doi = {10.1038/s41467-018-03313-w},
	abstract = {The nuclear pore complex (NPC) is the gatekeeper for nuclear transport in eukaryotic cells. A key component of the NPC is the central shaft lined with intrinsically disordered proteins (IDPs) known as FG-Nups, which control the selective molecular traffic. Here, we present an approach to realize artificial NPC mimics that allows controlling the type and copy number of FG-Nups. We constructed 34 nm-wide 3D DNA origami rings and attached different numbers of NSP1, a model yeast FG-Nup, or NSP1-S, a hydrophilic mutant. Using (cryo) electron microscopy, we find that NSP1 forms denser cohesive networks inside the ring compared to NSP1-S. Consistent with this, the measured ionic conductance is lower for NSP1 than for NSP1-S. Molecular dynamics simulations reveal spatially varying protein densities and conductances in good agreement with the experiments. Our technique provides an experimental platform for deciphering the collective behavior of IDPs with full control of their type and position.},
	language = {en},
	number = {1},
	urldate = {2020-05-30},
	journal = {Nature Communications},
	author = {Ketterer, Philip and Ananth, Adithya N. and Laman Trip, Diederik S. and Mishra, Ankur and Bertosin, Eva and Ganji, Mahipal and van der Torre, Jaco and Onck, Patrick and Dietz, Hendrik and Dekker, Cees},
	month = mar,
	year = {2018},
	pages = {902},
	file = {Full Text PDF:/home/gbubnis/Zotero/storage/GN8ZTMX7/Ketterer et al. - 2018 - DNA origami scaffold for studying intrinsically di.pdf:application/pdf;Snapshot:/home/gbubnis/Zotero/storage/74TMUQ4T/s41467-018-03313-w.html:text/html}
}

@article{chanturiya_flickering_1997,
	title = {Flickering fusion pores comparable with initial exocytotic pores occur in protein-free phospholipid bilayers},
	volume = {94},
	copyright = {© 1997},
	issn = {0027-8424, 1091-6490},
	url = {https://www.pnas.org/content/94/26/14423},
	doi = {10.1073/pnas.94.26.14423},
	abstract = {For the act of membrane fusion, there are two competing, mutually exclusive molecular models that differ in the structure of the initial pore, the pathway for ionic continuity between formerly separated volumes. Because biological “fusion pores” can be as small as ionic channels or gap junctions, one model posits a proteinaceous initial fusion pore. Because biological fusion pore conductance varies widely, another model proposes a lipidic initial pore. We have found pore opening and flickering during the fusion of protein-free phospholipid vesicles with planar phospholipid bilayers. Fusion pore formation appears to follow the coalescence of contacting monolayers to create a zone of hemifusion where continuity between the two adherent membranes is lipidic, but not aqueous. Hypotonic stress, causing tension in the vesicle membrane, promotes complete fusion. Pores closed soon after opening (flickering), and the distribution of fusion pore conductance appears similar to the distribution of initial fusion pores in biological fusion. Because small flickering pores can form in the absence of protein, the existence of small pores in biological fusion cannot be an argument in support of models based on proteinaceous pores. Rather, these results support the model of a lipidic fusion pore developing within a hemifused contact site.},
	language = {en},
	number = {26},
	urldate = {2020-05-30},
	journal = {Proceedings of the National Academy of Sciences},
	author = {Chanturiya, Alexandr and Chernomordik, Leonid V. and Zimmerberg, Joshua},
	month = dec,
	year = {1997},
	keywords = {exocytosis, membrane fusion, membranes, viral fusion},
	pages = {14423--14428},
	file = {Full Text PDF:/home/gbubnis/Zotero/storage/DUZUEFPQ/Chanturiya et al. - 1997 - Flickering fusion pores comparable with initial ex.pdf:application/pdf;Snapshot:/home/gbubnis/Zotero/storage/JZ8AL8KE/14423.html:text/html}
}

@article{kozlovsky_lipid_2002,
	title = {Lipid intermediates in membrane fusion: formation, structure, and decay of hemifusion diaphragm.},
	volume = {83},
	issn = {0006-3495},
	shorttitle = {Lipid intermediates in membrane fusion},
	url = {https://www.ncbi.nlm.nih.gov/pmc/articles/PMC1302349/},
	abstract = {Lipid bilayer fusion is thought to involve formation of a local hemifusion connection, referred to as a fusion stalk. The subsequent fusion stages leading to the opening of a fusion pore remain unknown. The earliest fusion pore could represent a bilayer connection between the membranes and could be formed directly from the stalk. Alternatively, fusion pore can form in a single bilayer, referred to as hemifusion diaphragm (HD), generated by stalk expansion. To analyze the plausibility of stalk expansion, we studied the pathway of hemifusion theoretically, using a recently developed elastic model. We show that the stalk has a tendency to expand into an HD for lipids with sufficiently negative spontaneous splay, ({\textasciitilde})J(s){\textless} 0. For different experimentally relevant membrane configurations we find two characteristic values of the spontaneous splay. ({\textasciitilde})J*(s) and ({\textasciitilde})J**(s), determining HD dimension. The HD is predicted to have a finite equilibrium radius provided that the spontaneous splay is in the range ({\textasciitilde})J**(s){\textless} ({\textasciitilde})J(s){\textless}({\textasciitilde})J*(s), and to expand infinitely for ({\textasciitilde})J(s){\textless}({\textasciitilde})J**(s). In the case of common lipids, which do not fuse spontaneously, an HD forms only under action of an external force pulling the diaphragm rim apart. We calculate the dependence of the HD radius on this force. To address the mechanism of fusion pore formation, we analyze the distribution of the lateral tension emerging in the HD due to the establishment of lateral equilibrium between the deformed and relaxed portions of lipid monolayers. We show that this tension concentrates along the HD rim and reaches high values sufficient to rupture the bilayer and form the fusion pore. Our analysis supports the hypothesis that transition from a hemifusion to a fusion pore involves radial expansion of the stalk.},
	number = {5},
	urldate = {2020-05-30},
	journal = {Biophysical Journal},
	author = {Kozlovsky, Yonathan and Chernomordik, Leonid V and Kozlov, Michael M},
	month = nov,
	year = {2002},
	pages = {2634--2651},
	file = {PubMed Central Full Text PDF:/home/gbubnis/Zotero/storage/PQAJ6G3A/Kozlovsky et al. - 2002 - Lipid intermediates in membrane fusion formation,.pdf:application/pdf}
}

@article{nikolaus_direct_2010,
	title = {Direct {Visualization} of {Large} and {Protein}-{Free} {Hemifusion} {Diaphragms}},
	volume = {98},
	issn = {0006-3495},
	url = {https://www.ncbi.nlm.nih.gov/pmc/articles/PMC2849057/},
	doi = {10.1016/j.bpj.2009.11.042},
	abstract = {Fusion of cellular membranes is a ubiquitous biological process requiring remodeling of two phospholipid bilayers. We believe it is very likely that merging of membranes proceeds via similar sequential intermediates. Contacting membranes form a stalk between the proximal leaflets that expands radially into an hemifusion diaphragm (HD) and subsequently open to a fusion pore. Although considered to be a key intermediate in fusion, direct experimental verification of this structure is difficult due to its transient nature. Using confocal fluorescence microscopy we have investigated the fusion of giant unilamellar vesicles (GUVs) containing phosphatidylserine and fluorescent virus derived transmembrane peptides or membrane proteins in the presence of divalent cations. Time-resolved imaging revealed that fusion was preceded by displacement of peptides and fluorescent lipid analogs from the GUV-GUV adhesion region. A detailed analysis of this area being several μm in size revealed that peptides were completely sequestered as expected for an HD. Lateral distribution of lipid analogs was consistent with formation of an HD but not with the presence of two adherent bilayers. Formation and size of the HD were dependent on lipid composition and peptide concentration.},
	number = {7},
	urldate = {2020-05-30},
	journal = {Biophysical Journal},
	author = {Nikolaus, Jörg and Stöckl, Martin and Langosch, Dieter and Volkmer, Rudolf and Herrmann, Andreas},
	month = apr,
	year = {2010},
	pages = {1192--1199},
	file = {PubMed Central Full Text PDF:/home/gbubnis/Zotero/storage/TYX2IH6M/Nikolaus et al. - 2010 - Direct Visualization of Large and Protein-Free Hem.pdf:application/pdf}
}

@book{berkowitz_biomembrane_2019,
	title = {Biomembrane {Simulations}: {Computational} {Studies} of {Biological} {Membranes}},
	isbn = {978-1-351-06029-5},
	shorttitle = {Biomembrane {Simulations}},
	abstract = {Due to recent advancements in the development of numerical algorithms and computational hardware, computer simulations of biological membranes, often requiring use of substantial computational resources, are now reaching a mature stage. Since molecular processes in membranes occur on a multitude of spatial and time scales, molecular simulations of membranes can also serve as a testing ground for use of multi-scale simulation techniques.   This book addresses some of the important issues related to understanding properties and behavior of model biological membranes and it    Shows how simulations improve our understanding of biological membranes and makes connections with experimental results.     Presents a careful discussion of the force fields used in the membrane simulations including detailed all-atom fields and coarse-grained fields.     Presents a continuum description of membranes.     Discusses a variety of issues such as influence of membrane surfaces on properties of water, interaction between membranes across water, nanoparticle permeation across the membrane, action of anesthetics and creation of inhomogeneous regions in membranes.     Discusses important methodological issues when using simulations to examine phenomena such as pore creation and permeation across membranes.     Discusses progress recently achieved in modeling bacterial membranes.   It will be a valuable resource for graduate students, researchers and instructors in biochemistry, biophysics, pharmacology, physiology, and computational biology.},
	language = {en},
	publisher = {CRC Press},
	author = {Berkowitz, Max L.},
	month = apr,
	year = {2019},
	keywords = {Medical / Biochemistry, Science / Chemistry / Physical \& Theoretical, Science / Physics / General}
}

@incollection{awasthi_free-energy_2019,
	address = {Boca Raton},
	edition = {1},
	title = {Free-{Energy} {Calculations} of {Pore} {Formation} in {Lipid} {Membranes}},
	isbn = {978-1-351-06031-8},
	language = {en},
	booktitle = {Biomembrane {Simulations}: {Computational} {Studies} of {Biological} {Membranes}},
	publisher = {CRC Press},
	author = {Awasthi, N and Hub, J.S., J. P. M.},
	year = {2019},
	keywords = {Effective Pair Potential, Mole Cular Cage, Oxygen Position, Protein Hydration, Spectral CORREl Density}
}

@article{kalienkova_stepwise_2019,
	title = {Stepwise activation mechanism of the scramblase {nhTMEM16} revealed by cryo-{EM}},
	volume = {8},
	issn = {2050-084X},
	url = {https://doi.org/10.7554/eLife.44364},
	doi = {10.7554/eLife.44364},
	abstract = {Scramblases catalyze the movement of lipids between both leaflets of a bilayer. Whereas the X-ray structure of the protein nhTMEM16 has previously revealed the architecture of a Ca2+-dependent lipid scramblase, its regulation mechanism has remained elusive. Here, we have used cryo-electron microscopy and functional assays to address this question. Ca2+-bound and Ca2+-free conformations of nhTMEM16 in detergent and lipid nanodiscs illustrate the interactions with its environment and they reveal the conformational changes underlying its activation. In this process, Ca2+ binding induces a stepwise transition of the catalytic subunit cavity, converting a closed cavity that is shielded from the membrane in the absence of ligand, into a polar furrow that becomes accessible to lipid headgroups in the Ca2+-bound state. Additionally, our structures demonstrate how nhTMEM16 distorts the membrane at both entrances of the subunit cavity, thereby decreasing the energy barrier for lipid movement.},
	urldate = {2020-06-02},
	journal = {eLife},
	author = {Kalienkova, Valeria and Clerico Mosina, Vanessa and Bryner, Laura and Oostergetel, Gert T and Dutzler, Raimund and Paulino, Cristina},
	editor = {Aldrich, Richard and Swartz, Kenton Jon and Ballesteros, Angela and Meyerson, Joel and Long, Stephen Barstow},
	month = feb,
	year = {2019},
	keywords = {activation mechanism, cryo-EM, lipid scrambling, membrane interaction, TMEM16},
	pages = {e44364},
	file = {Full Text PDF:/home/gbubnis/Zotero/storage/A4IE3Z3Q/Kalienkova et al. - 2019 - Stepwise activation mechanism of the scramblase nh.pdf:application/pdf}
}

@article{oostenbrink_biomolecular_2004,
	title = {A biomolecular force field based on the free enthalpy of hydration and solvation: the {GROMOS} force-field parameter sets {53A5} and {53A6}},
	volume = {25},
	issn = {0192-8651},
	shorttitle = {A biomolecular force field based on the free enthalpy of hydration and solvation},
	doi = {10.1002/jcc.20090},
	abstract = {Successive parameterizations of the GROMOS force field have been used successfully to simulate biomolecular systems over a long period of time. The continuing expansion of computational power with time makes it possible to compute ever more properties for an increasing variety of molecular systems with greater precision. This has led to recurrent parameterizations of the GROMOS force field all aimed at achieving better agreement with experimental data. Here we report the results of the latest, extensive reparameterization of the GROMOS force field. In contrast to the parameterization of other biomolecular force fields, this parameterization of the GROMOS force field is based primarily on reproducing the free enthalpies of hydration and apolar solvation for a range of compounds. This approach was chosen because the relative free enthalpy of solvation between polar and apolar environments is a key property in many biomolecular processes of interest, such as protein folding, biomolecular association, membrane formation, and transport over membranes. The newest parameter sets, 53A5 and 53A6, were optimized by first fitting to reproduce the thermodynamic properties of pure liquids of a range of small polar molecules and the solvation free enthalpies of amino acid analogs in cyclohexane (53A5). The partial charges were then adjusted to reproduce the hydration free enthalpies in water (53A6). Both parameter sets are fully documented, and the differences between these and previous parameter sets are discussed.},
	language = {eng},
	number = {13},
	journal = {Journal of Computational Chemistry},
	author = {Oostenbrink, Chris and Villa, Alessandra and Mark, Alan E. and van Gunsteren, Wilfred F.},
	month = oct,
	year = {2004},
	keywords = {Algorithms, Amino Acids, Computer Simulation, Cyclohexanes, Energy Transfer, Models, Chemical, Protein Folding, Solvents, Thermodynamics, Water},
	pages = {1656--1676}
}

@article{shirts_statistically_2008,
	title = {Statistically optimal analysis of samples from multiple equilibrium states},
	volume = {129},
	issn = {0021-9606},
	url = {https://aip.scitation.org/doi/10.1063/1.2978177},
	doi = {10.1063/1.2978177},
	number = {12},
	urldate = {2020-06-07},
	journal = {The Journal of Chemical Physics},
	author = {Shirts, Michael R. and Chodera, John D.},
	month = sep,
	year = {2008},
	pages = {124105},
	file = {Full Text:/home/gbubnis/Zotero/storage/M7Z7W2PS/Shirts and Chodera - 2008 - Statistically optimal analysis of samples from mul.pdf:application/pdf;Snapshot:/home/gbubnis/Zotero/storage/5KWJ2A5L/1.html:text/html}
}
