## Supplementary figures and images for "Sequential water and headgroup merger: Membrane poration paths and energetics from MD simulations"

### fig-06-tsm-snapshots-Igizmo.png

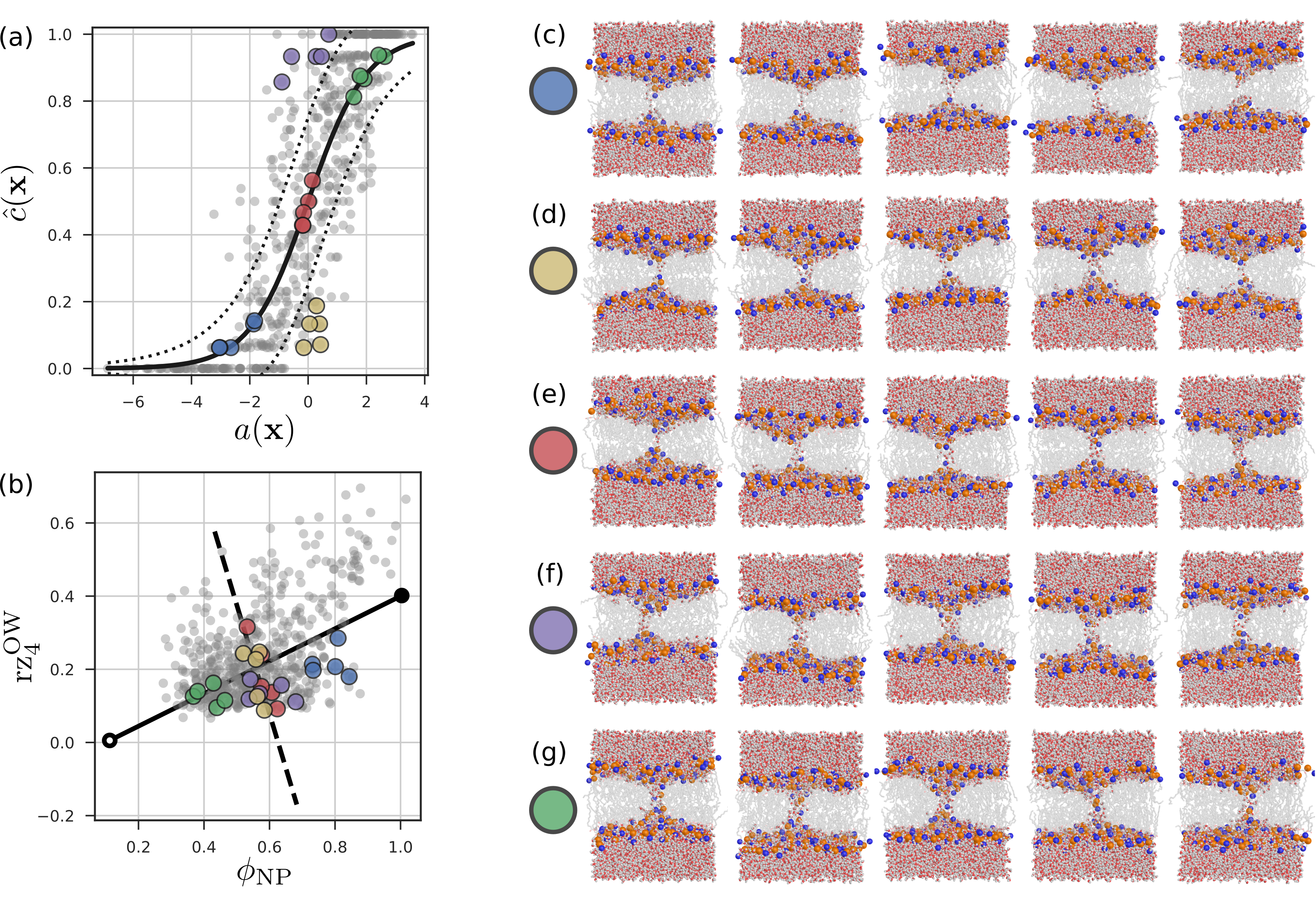
